# Large-scale analysis of interindividual variability in single and paired-pulse TMS data: results from the ‘Big TMS Data Collaboration’

**DOI:** 10.1101/2021.01.24.428014

**Authors:** Daniel T. Corp, Hannah G. K. Bereznicki, Gillian M. Clark, George J. Youssef, Peter J. Fried, Ali Jannati, Charlotte B. Davies, Joyce Gomes-Osman, Melissa Kirkovski, Natalia Albein-Urios, Paul B. Fitzgerald, Giacomo Koch, Vincenzo Di Lazzaro, Alvaro Pascual-Leone, Peter G. Enticott, the Big TMS Data Collaboration

## Abstract

**Objective:** Interindividual variability of single and paired-pulse TMS data has limited the clinical and experimental applicability of these methods. This study brought together over 60 TMS researchers to create the largest known sample of individual participant single and paired-pulse TMS data to date, enabling a more comprehensive evaluation of factors driving response variability.

**Methods:** 118 corresponding authors provided deidentified individual TMS data. Mixed-effects regression investigated a range of individual and study level variables for their contribution to variability in response to single and pp TMS data.

**Results:** 687 healthy participant’s TMS data was pooled across 35 studies. Target muscle, pulse waveform, neuronavigation use, and TMS machine significantly predicted an individual’s single pulse TMS amplitude. Baseline MEP amplitude, M1 hemisphere, and biphasic AMT significantly predicted SICI response. Baseline MEP amplitude, test stimulus intensity, interstimulus interval, monophasic RMT, monophasic AMT, and biphasic RMT significantly predicted ICF response. Age, M1 hemisphere, and TMS machine significantly predicted motor threshold.

**Conclusions:** This large-scale analysis has identified a number of factors influencing participants’ responses to single and paired pulse TMS. We provide specific recommendations to increase the standardisation of TMS methods within and across laboratories, thereby minimising interindividual variability in single and pp TMS data.

**Highlights:**

- 687 healthy participant’s TMS data was pooled across 35 studies
- Significant relationships between age and resting motor threshold
- Significant relationships between baseline MEP amplitude and SICI/ICF

## 1. Introduction

Single and paired-pulse (pp) TMS protocols are used to measure neural excitability within the primary motor cortex (M1) (Hallett 2000). However, these measures of M1 excitability have been shown to vary significantly between individuals (Iscan et al. 2016, Orth et al. 2003). A lack of understanding of the factors driving this variability has restricted greater application of single and pp TMS as a clinical and experimental tool (Iscan et al. 2016). Many studies have investigated this issue, yet there are conflicting findings in relation to the role of individual factors such as age (Cahn et al. 2003, Peinemann et al. 2001) and gender (Cahn et al. 2003, Shibuya et al. 2016), and also methodological factors such as the stimulus intensity used (Cosentino et al. 2018, Ibáñez et al. 2020, Ilić et al. 2002), and the hemisphere stimulated (Ilic et al. 2004, Maeda et al. 2002). Some of these conflicting findings are likely caused by small sample sizes inherent to most single-site studies (Fried et al. 2017a, Gilbert et al. 2005). To attempt to overcome this limitation, we recently formed the ‘Big TMS Data collaboration’ (Supplementary file 1) to combine individual participant TMS data across multiple studies. In the first instance, we used mixed-model regression to analyse data across 22 distinct datasets and demonstrate the variables driving interindividual variability in response to theta-burst stimulation (TBS) (Corp et al. 2020). Here we employ the same method, combining data from 35 TMS studies, to investigate the factors accounting for interindividual variability in response to single and pp TMS. The collation of multiple data-sets allowed us to more thoroughly examine sources of variability demonstrated by previous single and pp TMS studies, such as age, gender, and baseline MEP amplitude (Cahn et al. 2003, Shibuya et al. 2016, Strube et al. 2015), and also to further explore the possible influence of less examined variables on single and pp response, such as TMS machine, target muscle, and neuronavigation.

## 2. Methods

This project was deemed exempt from ethical review by the Deakin University Human Research Ethics Committee because it involved only the use of pre-existing, non-identifiable or re-identifiable data. All primary studies had been approved by local institutional review boards, and all participants had provided informed consent.

### 2.1 Article identification strategy

This analysis comes from a larger project collecting individual participant single and pp TMS data, input-output (I/O) curve data, and TBS data. Systematic search procedures are described in detail our companion paper (Corp et al. 2020), and the full search syntax is provided in Supplementary file 2. Inclusion criteria were: studies using a figure-of-eight coil; studies measuring TMS responses from intrinsic hand muscles of humans; and studies that collected baseline and conditioned MEP amplitudes. If an article met inclusion criteria, the corresponding authors of studies were emailed to ask for participants’ age, gender, motor threshold, and baseline and conditioned MEP amplitudes. Corresponding authors were asked to deidentify data prior to sending. A number of other studies were also included via informal data sharing with colleagues (Corp et al. 2020).

### 2.2 Variables of interest and data used for present analyses

Only healthy participant data were analysed within the present paper. To investigate interindividual variability for single pulse MEP amplitude, we used baseline MEP responses collected at 120% of RMT as our dependent variable (DV), collected across TBS, paired-pulse, and I/O curve datasets.

This intensity was chosen as the DV because it was the most commonly used single-pulse TMS intensity, enabling comparison across multiple studies (see Results, Table 3). We were not able to collect sufficient input/output curve data to analyse MEP amplitudes across a range of TS intensities. For SICI and ICF, each individual’s mean conditioned MEP amplitude was normalised to their mean baseline MEP amplitude (‘normalised MEP’) using the equation: (conditioned MEP amplitude / baseline MEP amplitude) x 100 (Amandusson et al. 2017, Di Lazzaro et al. 2006), where a value of 100% represents no change in conditioned MEP amplitudes. Note that the use of a ‘normalised MEP’ value or a percentage of change value (Fried et al. 2017b) (0% = no change in conditioned MEPs) provide the exact same results after regression analyses (Corp et al. 2020).

Because MT is extensively used as a measure of corticospinal excitability (Fried et al. 2017a, Kammer et al. 2001), we also investigated interindividual variability for four types of MT for which we had data: monophasic RMT, monophasic AMT, biphasic RMT and biphasic AMT. In addition to these four MTs being used as DVs (as above), MT may also predict single and pp TMS outcomes (Amandusson et al. 2017, Chen et al. 1998), thus these four MTs were also used as independent variables (IV) for our analyses of factors predicting single pulse MEP amplitude, and pp normalised MEP. Other IVs investigated were: age, gender, target muscle, M1 hemisphere, conditioning stimulus (CS) intensity, test stimulus (TS) intensity, pulse waveform (i.e. monophasic or biphasic), inter-stimulus interval (ISI), baseline MEP amplitude, the use/absence of neuronavigation, and TMS machine (Corp et al. 2020). Studies used either a Magstim 200^2^ TMS machine, a Magstim Rapid TMS machine, a Nexstim NBS TMS, or a MagPro TMS machine. We could not determine the specific MagPro model used in all studies, therefore these machines were grouped based on the brand. We controlled for pulse waveform in regression analyses to ensure that the effect of TMS machine was not due the differential use of monophasic or biphasic pulses. For TS intensity, studies used either 120% of RMT or a machine stimulus output evoking an MEP amplitude of 0.5 mV, 0.5 - 1 mV, 1 mV MEP, or 0.5 - 1.5 mV. To increase statistical power, we grouped these intensities into machine stimulus output evoking an MEP amplitude of 0.5 - 1.5 mV. Three studies did not use a TS intensity evoking 0.5 - 1.5 mV or 120% of RMT (Corp et al. 2015, Puri et al. 2016, Singh et al. 2016), and were therefore excluded from this comparison. We were not able to obtain baseline MEP amplitude data from one study (Munneke et al. 2013), thus these values were imputed as per the method of Corp et al. (2020). For studies that tested the effect of external interventions on TMS outcomes (e.g. exercise Singh et al. (2016)), only control/baseline data were analysed. We collected handedness data for 21 studies, yet there were only nine left handers represented across five studies, therefore this IV could not be analysed statistically.

We verified the accuracy of the data sent to us by comparing the results to group mean data in the corresponding published paper. In cases where we could not verify based on this group mean data, corresponding authors were contacted for clarification. In instances where data could not be verified, the study was excluded (n = 1).

All statistical analyses were conducted using Stata 13.0 (StataCorp, USA). First, data were checked for outliers using histograms and descriptive statistics. A number of outliers were detected in single and pp MEP data, therefore values falling outside of the 2^nd^ and 98^th^ percentiles were winsorized (Field 2009, Tukey 1962). Histograms prior to outlier winsorization are provided in Supplementary file 3.

### 2.3 Variability analyses

Prior to our main analyses investigating IVs predicting interindividual variability in single and pp TMS responses, we sought to characterise the variability of the data across our collected sample. As per the method of Brown et al. (2017), we calculated intraclass correlation coefficient (ICC), standard deviation (SD), and coefficient of variation (CV) (Brasil-Neto et al. 1992) values to assess within study, and between study variability of single and pp TMS data. Within study SDs and CVs were calculated using the mean MEP amplitude (or MT) of participants, and between study SDs and CVs were calculated using the mean MEP amplitude (or MT) of each study (Brown et al. 2017). ICC values < 0.50 were considered low; values 0.50 – 0.75 considered moderate; and > 0.75 considered high (Portney and Watkins 2009). High ‘within study’ ICC values reflect smaller variance within studies relative to larger variance between studies (Kline 2000).

Only one study (Beynel et al. 2014) assessed participants’ corticospinal excitability at multiple time-points, restricting an analysis of within-participant reliability over time. Yet, with the corresponding authors’ permission, we provide these (unpublished) data in Supplementary file 4.

### 2.4 Main regression analysis

Our main analyses investigated IVs predicting the aforementioned single, pp, and MT data. To do this, we employed the same regression analyses as described in detail in Corp et al. (2020). Briefly here, we used mixed-effects linear regression using a ‘one-step’ model as described by Riley et al. (2010), using ‘study ID’ as a random factor. Some data contained multiple entries by the same participants due to studies collecting multiple data-points across certain measures, such as ISI (e.g., 2 ms and 4 ms) (Croarkin et al. 2013). Thus, in these regressions we also included a random factor of ‘participant ID’ to maintain the nesting of these data-points within individual participants.

We used forward-stepwise regression in two stages for each TMS protocol (Bendel and Afifi 1977). Stage 1 regressions analysed the variance explained in the DV by each IV separately, while controlling for the age and gender of participants. IVs with p-values < 0.10 were added to the regression model in stage 2, while IVs with p-values > 0.10 were dropped (Corp et al. 2020). The stage 2 starting regression model comprised of all IVs that were p < 0.10 in stage 1. Consecutive regressions then iterated through IVs that were dropped in stage 1, to see whether these IVs now obtained a p-value < 0.10 controlling for IVs in the starting stage 2 model. Thus, the final regression model comprised of IVs that obtained a p-value < 0.10 in predicting the DV in either stage 1 or 2 regressions (Corp et al. 2020).

IVs were omitted from regression analyses for three possible reasons. First, an IV was omitted if it was not comprised of at least three studies within each IV level, given that unreliable estimates may have resulted from a smaller number of studies per level (Corp et al. 2020). For example, the IV ‘ISI’ was included only if all ISIs for which we had data (e.g. for SICI: 2 ms, 2.5 ms, 3 ms, and 4 ms) were used in at least three separate studies. Where some, but not all, levels of a given IV were represented across three or more studies, we compared these levels post-hoc (see below). Second, an IV was omitted if its inclusion led to a substantial reduction in the overall sample size of the regression analysis for that DV, due to that IV only being measured in a subset of studies. We defined a ‘substantial reduction of the regression sample size’ as cases where two or more studies were excluded from the regression analysis. Third, an IV was omitted because of collinearity, which occurred if two types of MTs were included in the same regression model. To avoid this, if two or more types of MTs had a p-value < 0.10 in stage 1 regressions, for stage 2 we included only the MT that was the strongest predictor of normalised MEP for that particular regression analysis.

Given the presence of non-linearity and non-normality, robust variance estimates were used for all regressions (Graubard and Korn 1996). Adjusted marginal means (just ‘marginal means’ henceforth) estimated the mean normalised MEP amplitude adjusted/controlled for all other variables in the regression model (Williams 2012). This allowed an interpretable estimate of the mean across the sample, and also for each level of categorical IVs (e.g. the levels ‘left’ and ‘right’ for the IV ‘M1 hemisphere’) (Williams 2012).

### 2.5 Post-hoc analyses

Where sufficient data, post-hoc analyses were run on IVs that were omitted from the main regression analyses for any of the three aforementioned reasons. In relation to reason three for omission (i.e. collinearity), different types of MT were always analysed in separate regression models, to assess their independent relationship to normalised MEP. Next, post-hoc pairwise comparisons were performed on significant IVs that had 3 or more levels (given that results from IVs with only 2 levels can be interpreted from the main regression output). Given their exploratory nature, these pairwise analyses were not corrected for multiple comparisons. Finally, scatterplots indicated possible non-linear relationships between normalised MEP and some continuous variables (e.g. age). Therefore, we re-analysed all (continuous variable) relationships that were included in the final regression model, or were significant in post-hoc analyses, using quadratic and cubic regression models (Davidson and MacKinnon 1993). All post-hoc analyses controlled for all other IVs in the final regression model.

### 2.6 Additional analyses

A number of additional analyses were performed to further explore the data. Marginal means following single pulse regression analysis indicated that 120% RMT MEP data did not reach 1 mV in amplitude. Therefore, we then assessed whether these MEP amplitudes were significantly lower in comparison to MEP amplitudes collected using the 1 mV method (i.e. stimulus intensity required to evoke a 1 mV MEP amplitude). To do this, we performed two-stage mixed-effects linear regression analysis, as above, including TS intensity (with levels of 120 RMT method and 1 mV method) as an IV. Given that controlling for other IVs may cause unwanted influence on 1 mV values, which were already adjusted by TMS operators to attain a 1 mV amplitude regardless of age, gender etc., we also repeated this analysis without the inclusion of these IVs (i.e. including only the TS intensity IV, and ‘study ID’ and ‘Participant ID’ as a random factors). This analysis did not include the imputed data of Munneke et al. (2013).

We then assessed a possible difference in MEP amplitude *variance* between these TS intensity methods. Here we used the same method as in our ‘variability analysis’, calculating SD and CV values of single pulse MEP amplitudes, yet split the sample to analyse SD and CV separately for studies that used the 120% RMT method, and the 1 mV method. Significance between the TS intensity methods was assessed using Levene’s robust test for equality of variances (Levene 1961). While lower variance may be expected for the 1mV method, given that operators specifically set the machine intensity to evoke a 1mV amplitude, we still thought it valuable to quantify these (possible) differences.

Lastly, we analysed correlations between the four types of MT. Because different studies use different methods for obtaining MTs and therefore vary in their average MT values, we normalised MTs to z-values within study, then performed Pearson’s correlation analyses on these z-values across the sample. This gives similar results to correlating MT values within studies, then taking the average of these correlations (Supplementary file 5).

## 3. Results

See Corp et al. (2020) for the PRISMA flowchart describing our initial systematic search. In total, 38 studies contributed individual participant data. Three studies were removed because they either included clinical populations only (2) (Kuppuswamy et al. 2015, Murdoch et al. 2016), or we were unable verify the accuracy of the sent data through email correspondence (1) (Malcolm et al. 2015). MT and single-pulse data were drawn from this larger sample of 35 studies and 687 healthy participants, which included theta-burst stimulation and I/O curve datasets in addition to pp data (Table 1). Pp TMS data were drawn from 16 studies, including 15 SICI and 14 ICF datasets comprising 295 healthy participants. Figure 1 shows the distribution of single, pp, and MT data.

**Figure 1.**
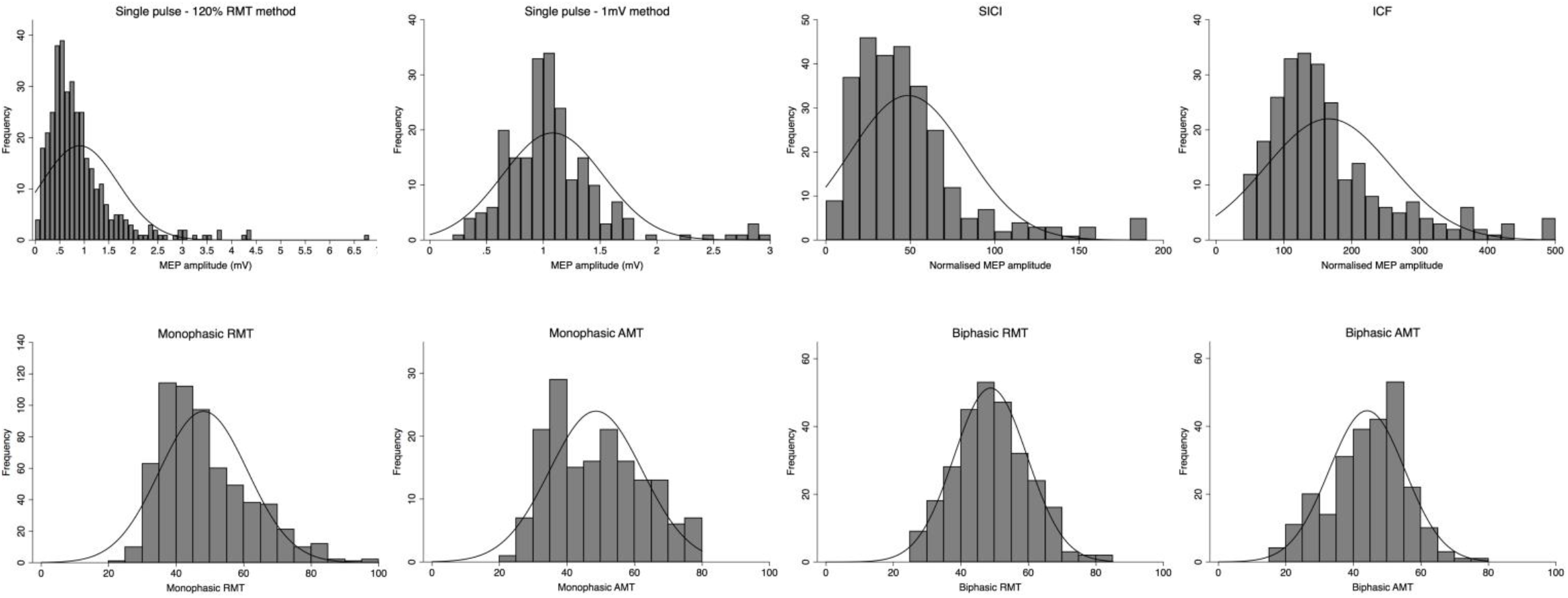
Distribution plots. Histograms of single pulse, paired pulse, and motor threshold data. 120% RMT data was used for single-pulse main regression analysis. These data were then compared to single pulse data using the 1 mV method in the ‘additional analyses’. In addition to differences in amplitude and variance (see Results), 120% RMT data appear positively skewed, also evidenced by low median value (0.73 mV). 1 mV method data median = 1.03 mV. So that each participant was only represented once within all histograms and scatterplots (multiple data points due to some studies using multiple ISIs, muscles, etc. – see Methods) we take each participant’s mean normalised MEP value across their multiple measurements. Note that in regression analyses, multiple measurements were dealt with by including ‘participant ID’ as a random factor – see Methods.

**Table 1.** Characteristics of included studies.

| **Study** | **Author/s** | **Participants** | **TMS protocols** |
| --- | --- | --- | --- |
| 1 | Barhoun (unp.) | 13 healthy (5F, 22.1 ± 3.0 y) | cTBS |
| 2 | Beynel et al. (2014) | 20 younger (14F, 26.4 ± 7.9 y), 19 older healthy (12F, 63.7 ± 1.7 y) | SICI, ICF |
| 3 | Busan et al., (2013) | 40 healthy adults (12F, 26.2 ± 6.6 y) | I/O curves |
| 4 | Capone et al. (2009) | 22 healthy (13F, 27.6 ± 9.0 y) | SICI, ICF |
| 5 | Corp et al. (2015) | 14 healthy (3F, 29.6 ± 6.7 y) | SICI, ICF |
| 6 | Cosentino et al. (2015) | 25 cluster headache patients (4F, 37.7 ± 10.5 y), 13 healthy (2F, 35.2 ± 11.2 y) | SICI, ICF |
| 7 | Croarkin et al. (2013) | 24 MDD (14F, 13.9 ± 2.1 y), 22 healthy (11F, 13.8 ± 2.2 y) | SICI, ICF |
| 8 | Di Lazzaro (unp.) | 17 healthy (5F, 23.9 ± 5.1 y) | SICI, ICF |
| 9 | Di Lazzaro et al. (2008) | 12 stroke patients (5F, 69.4 ± 9.5 y), 12 controls (2F, 63.2 ± 5.3 y) | iTBS & cTBS |
| 10 | Di Lazzaro et al. (2011) | 10 healthy (7F, 26.6 ± 4.1 y) | SICI, ICF, iTBS, cTBS |
| 11 | Dickins et al. (2015) | 20 younger (10F, 22.9 ± 2.5 y) and 20 older participants (10F, 70.2 ± 3.1 y) | iTBS |
| 12 | Dileone et al. (2016) | 16 healthy (10F, 23.2 ± 3.8 y) | iTBS |
| 13 | Do et al. (2018) | 20 healthy (14F, 26.5 ± 3.1 y) | cTBS |
| 14 | Fried et al. (2017) | 28 type 2 diabetes patients (12F, 65.8 ± 7.7 y), 22 AD patients (13F, 69.6 ± 7.4 y), 26 healthy (13F, 62.9 ± 8.9 y) | SICI, ICF, iTBS |
| 15 | Fuhl et al., (2015) | 10 healthy (1F, 24.6 ± 3.9 y) | I/O curves |
| 16 | Goldsworthy et al. (2016) | 18 healthy (10F, 22.1 ± 4.4 y) | iTBS |
| 17 | Gomes-Osman (unp.) | 17 healthy (10F, 30.0 ± 12.9 y) | SICI, ICF, iTBS |
| 18 | Helm et al. (2015) | 11 healthy (2F, 25 ± 4.3 y) | ICF |
| 19 | Hoseini et al., (2016) | 18-40 y | I/O curves |
| 20 | Jannati et al. (2017) | 30 healthy (3F, 36.0 ± 14.4 y) | cTBS |
| 21 | Koch et al. (2016) | 40 AD patients (17F, 71.0 ± 6.4 y) and 24 healthy (12F, 69.3 ± 2.3 y) | iTBS, cTBS |
| 22 | Lee et al. (2014) | 18 healthy (12F, 73.8 ± 5.1 y) | cTBS |
| 23 | Li et al. (2017) | 26 GAD patients (13F, 42 ± 9.7 y), 35 controls (20F, 41 ± 10.6 y) | SICI, ICF |
| 24 | McDonnell et al. (2013) | 25 healthy (9F, 26.8 ± 8.1 y) | cTBS |
| 25 | Lücke et al., (2014) | 9 healthy (3F, 25 ± 4.2 y) | I/O curves |
| 26 | Morris (unp.) | 15 healthy (9F, 25 ± 2.7 y) | SICI, ICF, iTBS |
| 27 | Munneke et al. (2013) | 10 ALS patients (10M, 57.8 ± 1.8 y) and 10 controls (0F, 49.0 ± 3.6 y) | SICI, ICF, cTBS |
| 28 | Nettekoven et al. (2014) | 16 healthy (9F, 27.0 ± 3.0 y) | iTBS |
| 29 | Opie et al. (2013) | 13 sleep apnoea patients (2F, 42.6 ± 10.2 y), 11 controls (2F, 43.0 ± 10.3 y) | SICI, cTBS |
| 30 | Opie et al. (2015) | 13 younger (7F, 22.3 ± 3.8 y) and 15 older healthy (7F, 73.7 ± 4.0 y) | SICI |
| 31 | Puri et al. (2016) | 33 healthy (21F, 66.0 ± 4.8 y) | iTBS |
| 32 | Singh et al. (2016) | 10 healthy (6F, 25.4 ± 4.0 y) | SICI, ICF, cTBS |
| 33 | Vallence et al. (2015) | 18 healthy (10F, 23.1 ± 4.0 y) | cTBS |
| 34 | Vernet et al. (2014) | 10 healthy (5F, 33.0 ± 18.0 y) | cTBS |
| 35 | Young-Bernier et al. (2014) | 20 younger (13F, 22.3 ± 3.2 y) and 18 older healthy (9F, 70.1 ± 5.6 y) | iTBS |

Note: age mean and standard deviation are shown. Studies without paired-pulse data were used in single pulse and/or motor threshold analyses. Abbreviations: F = females; y = years old; GAD = generalised anxiety disorder; AD = Alzheimer's disease; ALS = amyotrophic lateral sclerosis; MDD = major depressive disorder; I/O = input/output; FDI = first dorsal interosseous; APB = abductor pollicis brevis.

### 3.1 Variability analyses

Table 2 shows measures of reliability for all TMS outcomes. 120% of RMT MEP amplitudes, SICI, and ICF demonstrated higher within, than between, study variance. This is also demonstrated by low ICC values for these outcomes, reflecting little grouping of within study values relative to the overall sample. Consistent with previous reports (Davila-Pérez et al. 2018, Fried et al. 2017a), within and between study reliability was higher for MTs than the aforementioned (120% of RMT) single pulse and pp TMS outcomes.

**Table 2.**
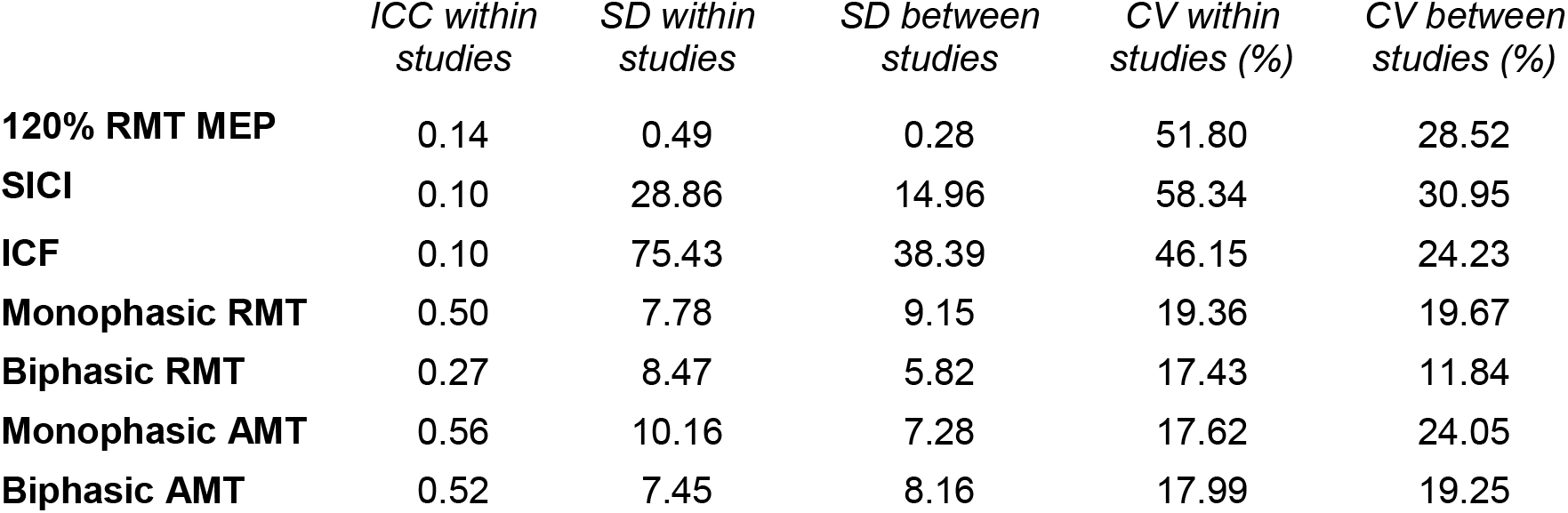
Variability of single and paired-pulse TMS data. ICC = intraclass correlation coefficient; SD = standard deviation; CV = coefficient of variation %.

### 3.2 Single pulse TMS regression analysis

The inclusion of any MT in the model would have substantially reduced the regression sample size. Thus, see post-hoc analyses for these relationships.

The final regression model showed that muscle, pulse waveform, the use of neuronavigation, and TMS machine were all significant predictors of 120% of RMT single-pulse MEP amplitude (Table 3). See Figure 2 for single pulse TMS marginal means.

**Table 3.**
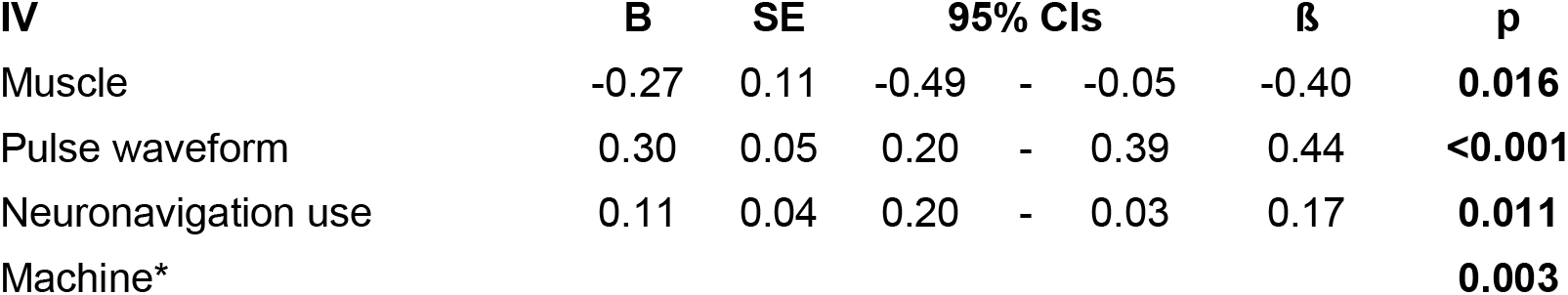
Final single pulse MEP amplitude regression model. B-values for categorical IVs show the differences between the IV levels in mV. e.g. the APB demonstrated 0.27 mV lower MEP amplitudes than the FDI. Bold denotes significance (p < 0.05). Participants = 341; studies = 17. *TMS machine had 3 levels (Magstim 200^2^, MagPro, and Nextstim), therefore main effect: *χ^2^* = 11.62, df = 2. See post-hocs for pairwise comparisons between levels.

**Figure 2.**
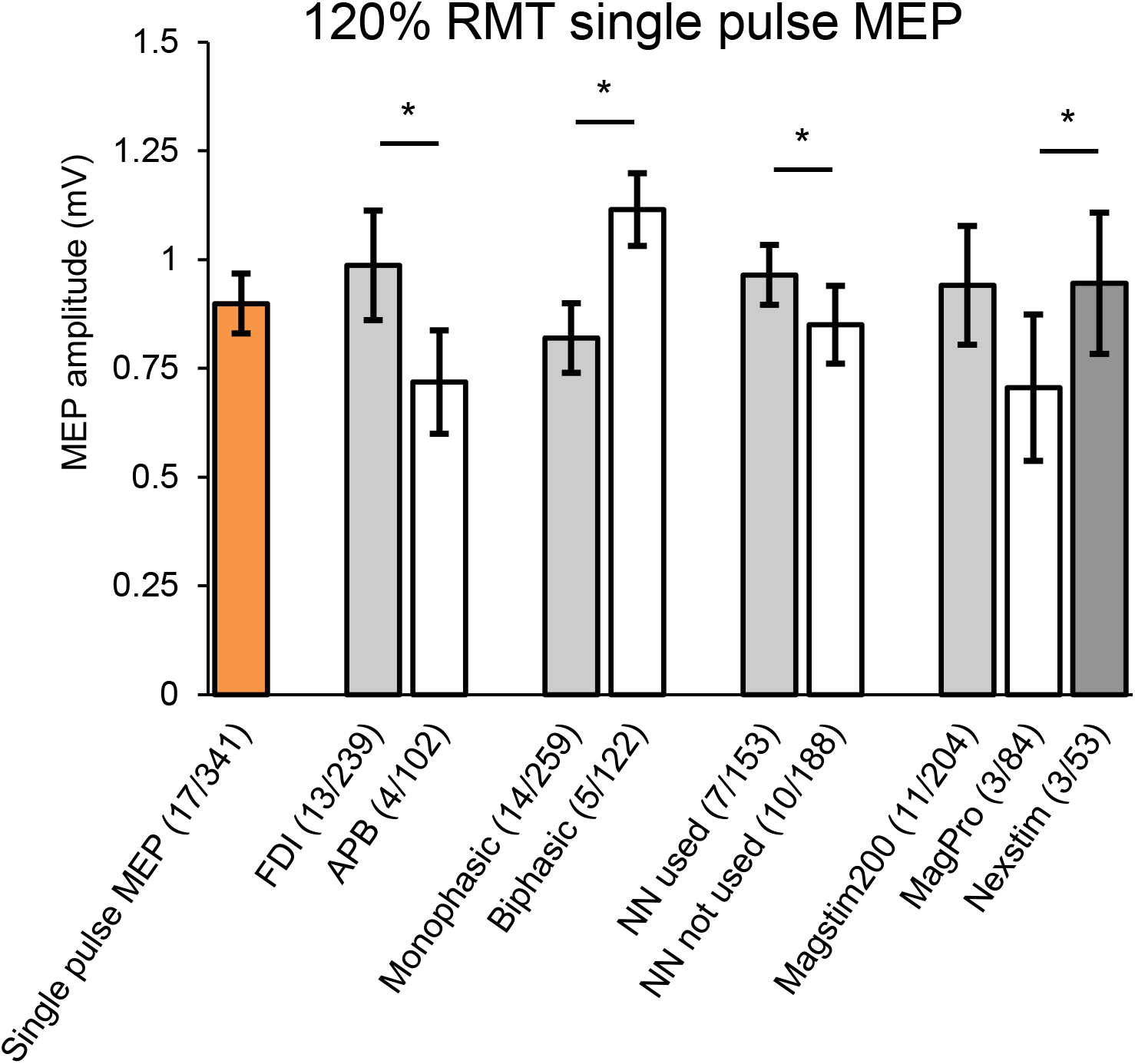
Marginal means for 120% RMT single pulse MEPs. Marginal means provide an estimate of normalised MEP, adjusted for all variables in the final model. Orange bar shows the overall marginal mean for single pulse MEPs. Grey and white bars show marginal means for each level of the IVs muscle, pulse waveform, neuronavigation (NN), and TMS machine. * denotes a significant difference between levels (p < 0.05). Error bars show 95% confidence intervals. Brackets show (studies/ participants). Difference between Magstim 2002 and MagPro was close to significance (p = 0.078).

Other IVs not included in final regression model had p-values > 0.10 in both stage 1 and 2 regressions (see Supplementary file 6 for all stage 1 and 2 results).

### 3.3 Single pulse TMS post-hoc analyses

When controlling for all IVs in the final regression model, all four types of MT were significantly negatively associated with single pulse MEP amplitude at 120% RMT. Monophasic RMT, B = −0.015; SE = 0.004; ß = 0.31; p < 0.001 (studies = 13; N = 248). Biphasic RMT, B = −0.020; SE = 0.005; ß = −0.31; p < 0.001 (studies = 8; N = 174). Monophasic AMT, B = −0.010; SE = 0.004; ß = − 0.20; p = 0.024 (studies = 3; N = 62). Biphasic AMT, B = −0.017; SE = 0.006; ß = −0.29; p = 0.005 (studies = 9; N = 174). Figure 3 shows bivariate relationship between single-pulse MEP amplitude and monophasic RMT.

**Figure 3.**
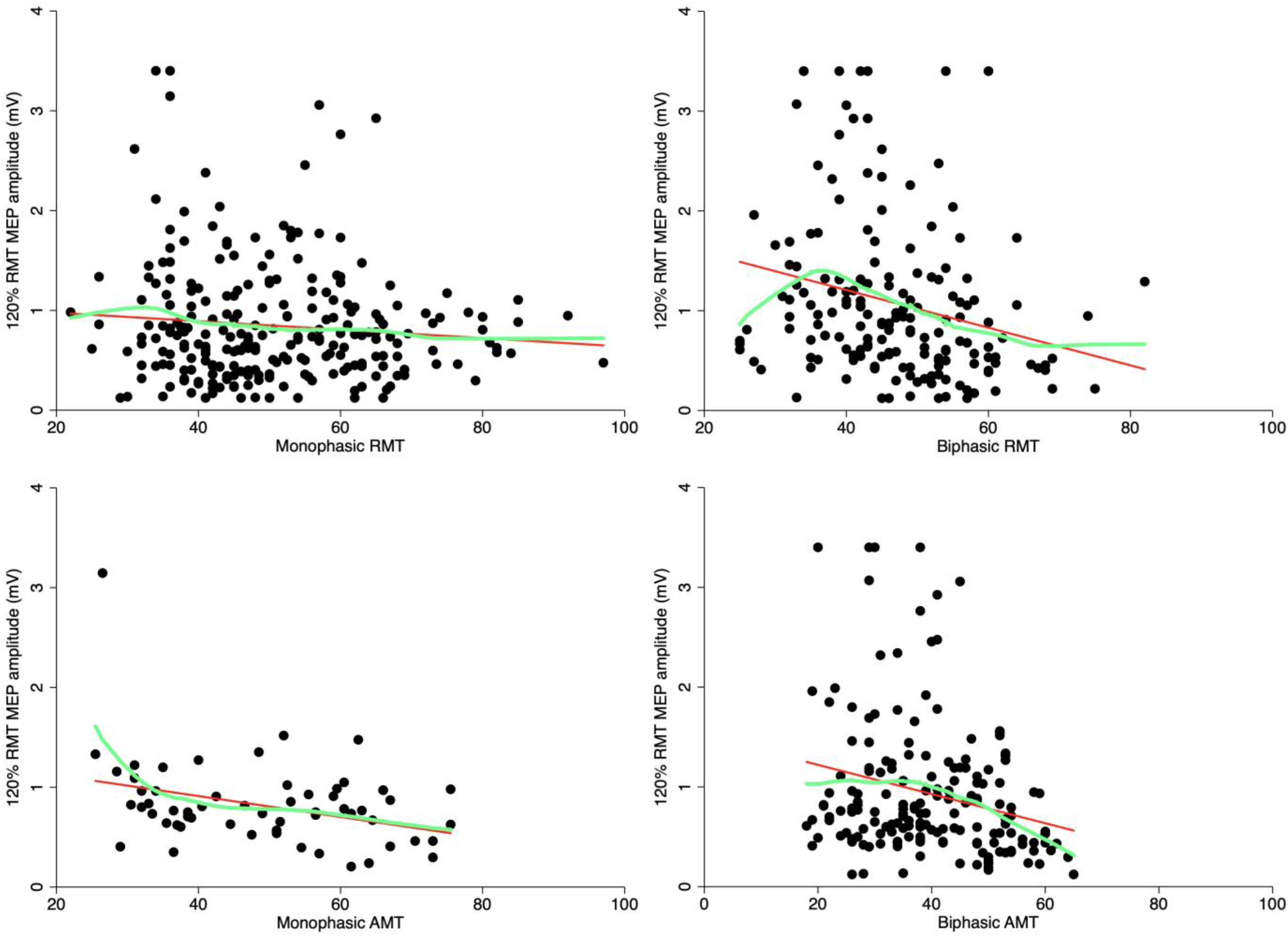
Relationships between 120% RMT single pulse MEPs and MTs. All relationships were significant in post-hoc regression analyses. Note that these scatterplots show raw bivariate relationships to give an indication of relationships only, see post-hoc section for results controlled for other IVs in the single pulse TMS model. Green lines fit a smoothed ‘lowess’ curve through data (smoothing level = 0.8, default).

In addition, non-linear analyses demonstrated a significant quadratic relationship between single pulse MEP amplitude and biphasic AMT (p = 0.042), and significant cubic relationships between single pulse MEP amplitude and biphasic RMT, and monophasic AMT (p = 0.001 and p = 0.010, respectively) (see Supplementary file 7 for scatterplots).

### 3.4 SICI regression analysis

IVs ‘TMS machine’, ‘CS intensity’, ‘pulse waveform’, and ‘ISI’ were omitted because they did not include at least three studies within each IV level, while biphasic AMT and biphasic AMT were p < 0.10 in stage 1 regressions but substantially reduced regression sample size, thus were analysed post-hoc. The final SICI regression model showed that baseline MEP and M1 hemisphere were both significant predictors of SICI normalised MEP (Table 4). M1 hemisphere was still significant when re-analysed including only data from only right handers (from the sample in which we had handedness data) (studies = 9; N = 144; B = −9.04; SE = 2.85; p = 0.002).

**Table 4.** Final SICI regression model. B-values for continuous IVs show the amount of increase in normalised MEP, for a one unit increase in the IV, after adjusting for all other variables in the model. i.e. a 1mV increase in baseline MEP resulted in a 23.29% reduction in SICI normalised MEP (greater inhibition). Bold denotes significance (p < 0.05). Participants = 283; studies = 15. See Figure 5 for IV levels.

| IV | B | SE | 95% CIs | | | $\beta$ | p |
| --- | --- | --- | --- | --- | --- | --- | --- |
| Age | 0.11 | 0.11 | -0.11 | - | 0.34 | 0.04 | 0.334 |
| Gender | 5.67 | 3.63 | -1.45 | - | 12.78 | 0.15 | 0.119 |
| Baseline MEP | -23.29 | 8.22 | -39.41 | - | -7.17 | -0.33 | **0.005** |
| Hemisphere | -4.01 | 1.73 | -7.41 | - | -0.62 | -0.10 | **0.021** |

Figure 4 shows bivariate relationships for continuous IVs baseline MEP and age, which were included in the final regression model. See Figure 5 for SICI marginal means.

**Figure 4.**
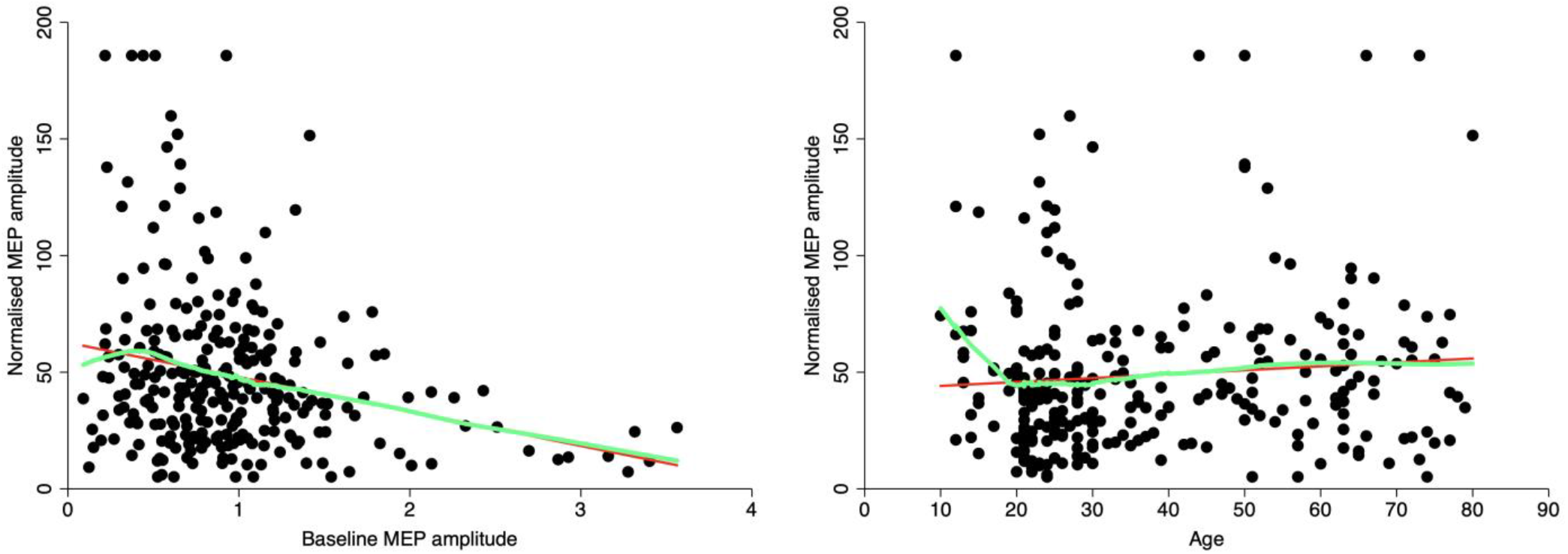
Relationships between continuous IVs and SICI. Baseline MEP amplitude was a significant predictor of SICI. Bivariate scatterplots give an indication of results only; see Table 4 for results controlled for other IVs. Green lines fit a smoothed ‘lowess’ curve through data. The appearance of a line of datapoints at the top (and to a lesser extent the bottom) of these (and other) scatterplots is due to winsorization; where small and large value outliers are converted to the value of the datapoint at the 2nd and 98th percentile (Field 2009, Tukey 1962) (see Methods).

**Figure 5.**
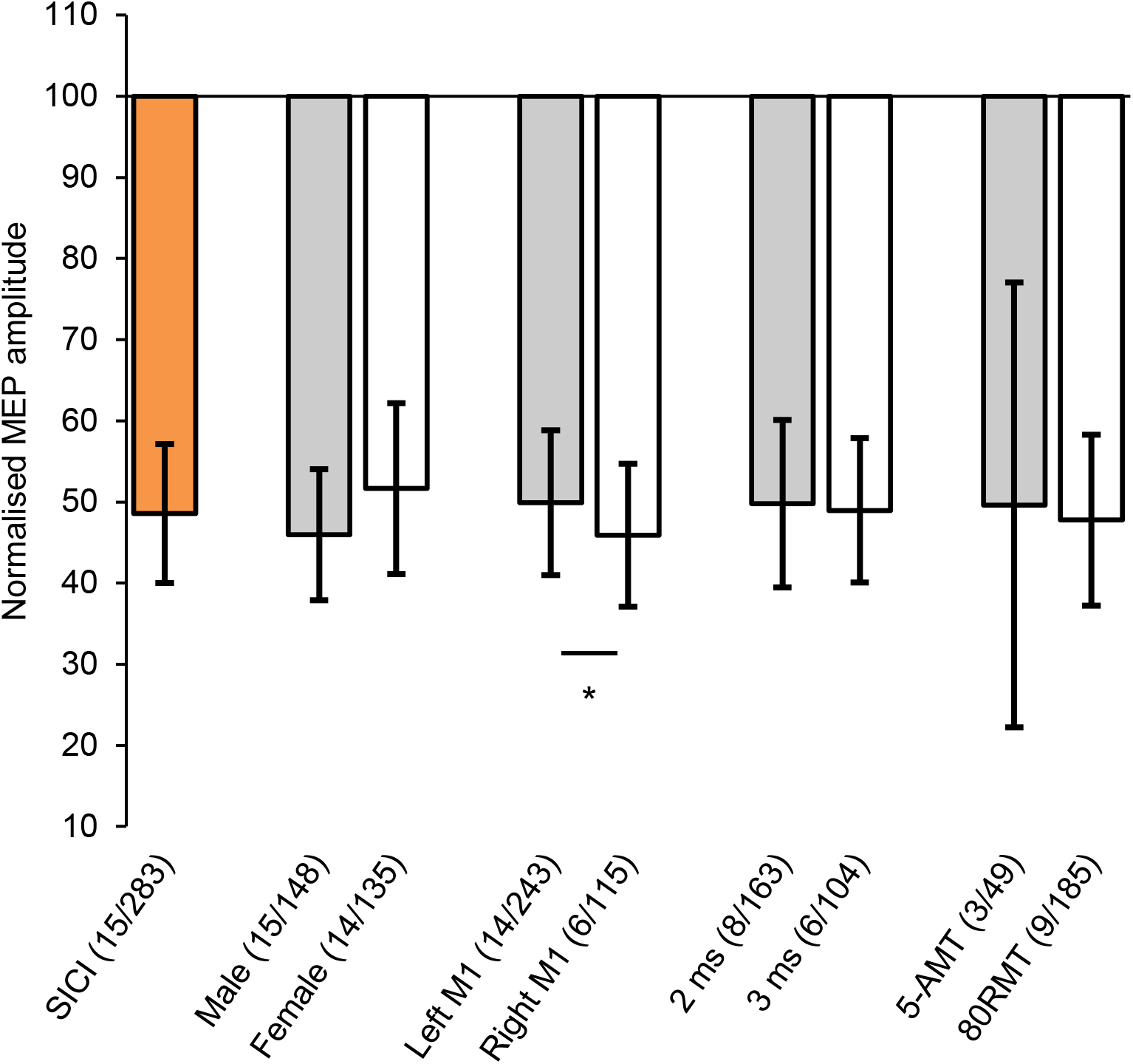
Marginal means for SICI normalised MEP. Orange bar shows the overall marginal mean for SICI. Grey and white bars show marginal means for each level of the IVs gender, M1 hemisphere, interstimulus interval and CS intensity (5% of machine intensity below AMT and 80% of RMT), which were included in the final model or post-hoc tests. * denotes a significant difference between levels (p < 0.05). All samples demonstrated significant inhibition (p < 0.001). Error bars show 95% confidence intervals. Brackets show (studies/ participants).

Other IVs not included in final regression model had p-values > 0.10 in both stage 1 and 2 regressions (see Supplementary file 8 for all stage 1 and 2 results).

### 3.5 SICI post-hoc analyses

CS intensity and ISI were omitted from the main analysis, yet we had sufficient data to compare SICI normalised MEP between studies that used an intensity of 80% of RMT to those that used a machine intensity 5% below AMT (5-AMT), and also ISI of 2 ms and 3 ms (> 3 studies for these levels). Neither comparison was significant (p = 0.900 and p = 0.778, respectively; Figure 5).

Biphasic AMT was a significant predictor of SICI normalised MEP when controlling for all IVs in the final model: 6 studies, 85 participants; B = −0.86; SE = 0.30; ß = −0.24; p = 0.004. Biphasic RMT was not a significant predictor of normalised MEP: 3 studies, 78 participants; B = 0.24; SE = 0.31; ß = 0.07; p = 0.426.

There were no significant non-linear relationships between SICI and age, baseline MEP amplitude, or biphasic AMT. Although the quadratic relationship between SICI and baseline MEP amplitude almost reached significance (p = 0.053).

### 3.6 ICF regression analysis

IVs ‘TMS machine’, ‘CS intensity’, ‘pulse waveform’, and ‘ISI’ were omitted from ICF regression due to insufficient data. The inclusion of any the MTs as IVs would have led to a substantial reduction in regression sample size, therefore these were analysed post-hoc.

The final regression model showed that baseline MEP amplitude and TS intensity (i.e. 120% RMT vs 0.5 - 1.5 mV methods) were significant predictors of ICF normalised MEP (Table 5 and Figure 6). See Figure 7 for ICF marginal means. Other IVs not included in final regression model had p-values > 0.10 in both stage 1 and 2 regressions (see Supplementary file 8 for all stage 1 and 2 results).

**Figure 6.**
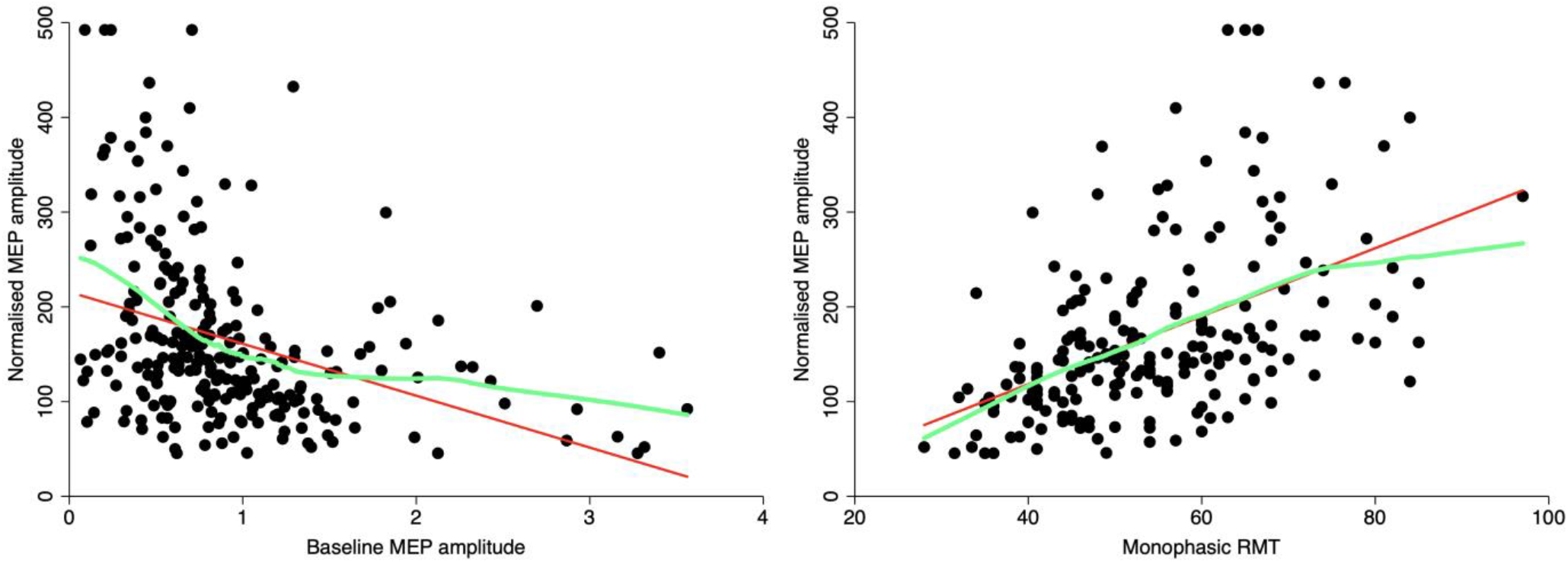
Relationships between continuous IVs and ICF. Baseline MEP and monophasic RMT were significant predictors of ICF MEP change. Bivariate scatterplots give an indication of results only; see Table 5 for results controlled for other IVs. Green lines fit a smoothed ‘lowess’ curve through data.

**Figure 7.**
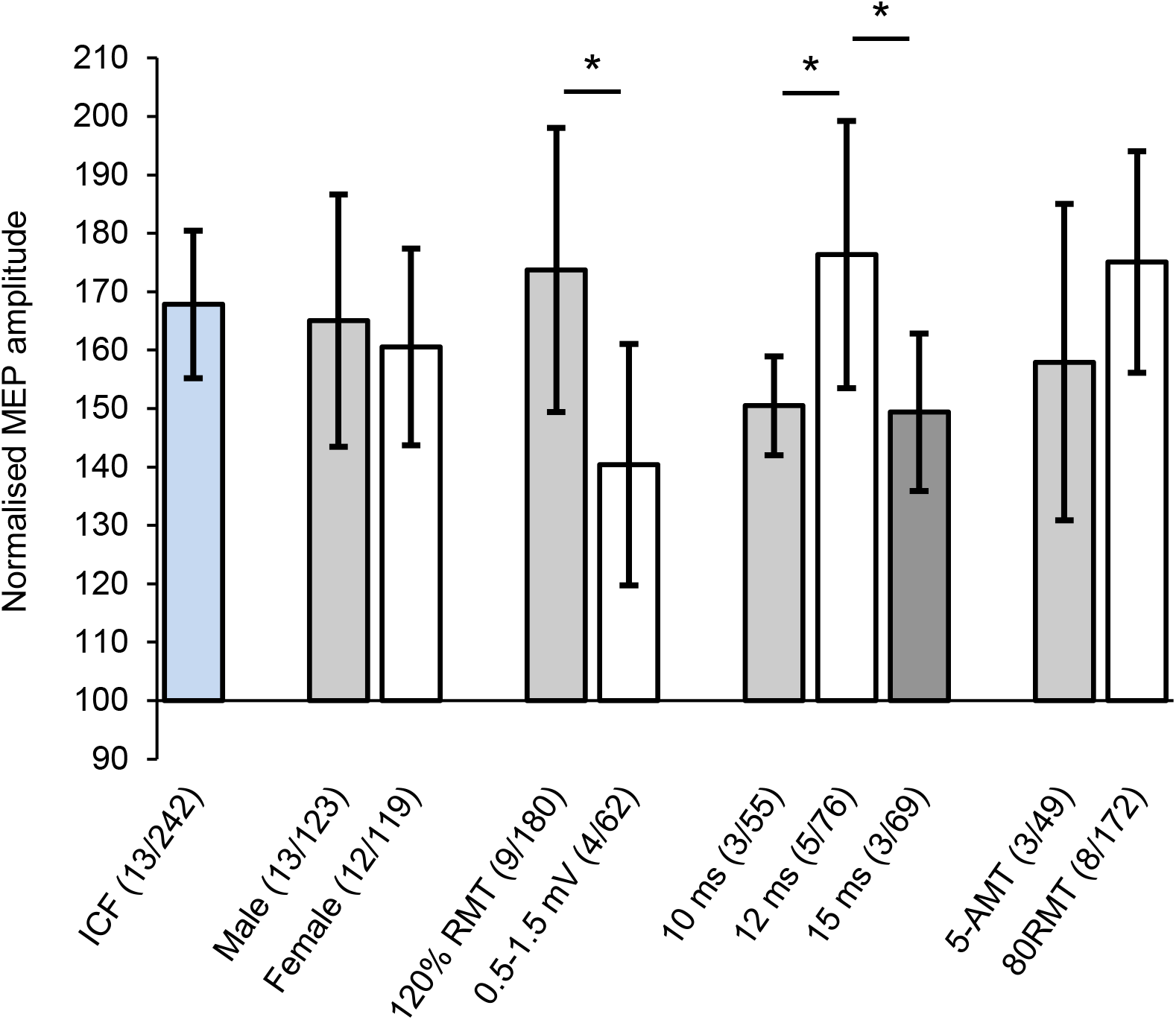
Marginal means for ICF normalised MEP. Blue bar shows the overall marginal mean for ICF. Grey and white bars show marginal means for each level of the IVs gender, TS intensity, ISI, and CS intensity (5% machine intensity below AMT vs. 80% of RMT) which were included in the final model or post-hoc tests. * denotes a significant difference between levels (p < 0.05). All samples demonstrated significant facilitation (p < 0.001). Error bars show 95% confidence intervals. Brackets show (studies/participants).

**Table 5.**
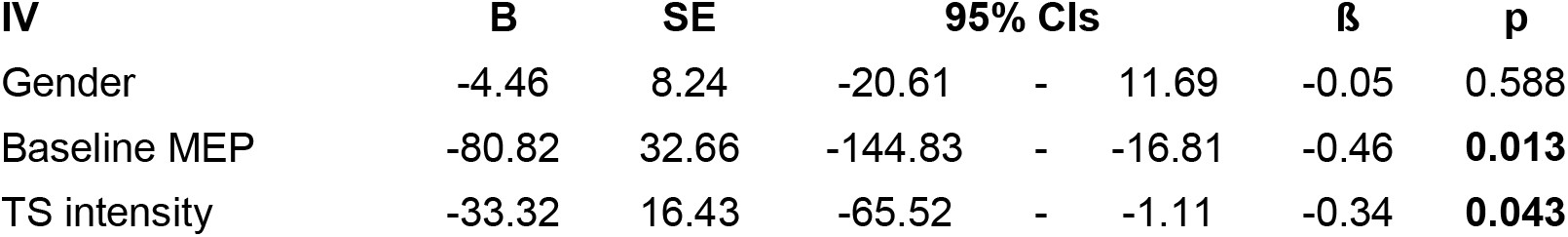
Final ICF regression model. Bold denotes significance (p < 0.05). Participants = 242; studies = 13. See Figure 7 for IV levels.

### 3.7 ICF post-hoc analyses

While CS intensity and ISI were omitted from the main analysis, we had sufficient data to compare 80% of RMT to 5-AMT CS intensities and to compare 10 ms, 12, ms, and 15 ms ISIs. The CS intensity comparison was not significant (p = 0.303), however for ISI, there was significantly higher ICF for 12 ms ISI data compared to both 10 ms (p = 0.043) and 15 ms ISI data (p = 0.042) (Figure 7).

Of the four types of MT, only biphasic AMT was not significantly positively associated with ICF normalised MEP. Monophasic RMT, B = 2.09; SE = 0.55; ß = 0.29; p < 0.001 (studies = 11; N = 193). Biphasic RMT, B = 1.46; SE = 0.30; ß = 0.16; p < 0.001 (studies = 3; N = 79). Monophasic AMT, B = 1.33; SE = 0.48; ß = 0.19; p < 0.005 (studies = 3; N = 84).

Non-linear analyses demonstrated a significant quadratic and cubic relationship between ICF and baseline MEP amplitude (p = 0.025 and p = 0.044, respectively) (Figure 6). There was also a significant quadratic relationship between ICF and monophasic AMT (p = 0.001), and a significant cubic relationship between ICF and biphasic RMT (scatterplots in Supplementary file 9).

### 3.8 MT regression analyses

Table 6 shows the four final regression models, demonstrating IVs predicting each type of MT (see captions for IVs omitted due to insufficient data). Age, M1 hemisphere, and TMS machine were significant predictors of different types of MT. There was still higher monophasic RMT for the left hemisphere when including only data from only right handers (from the restricted sample in which we had handedness data), however this effect was now non-significant (studies = 18; N = 319; B = −0.69; SE = 0.39; p = 0.079). Age demonstrated a significant positive relationship with monophasic RMT and biphasic RMT (Figure 8). See Figure 9 for marginal means of each IV level.

**Figure 8.**
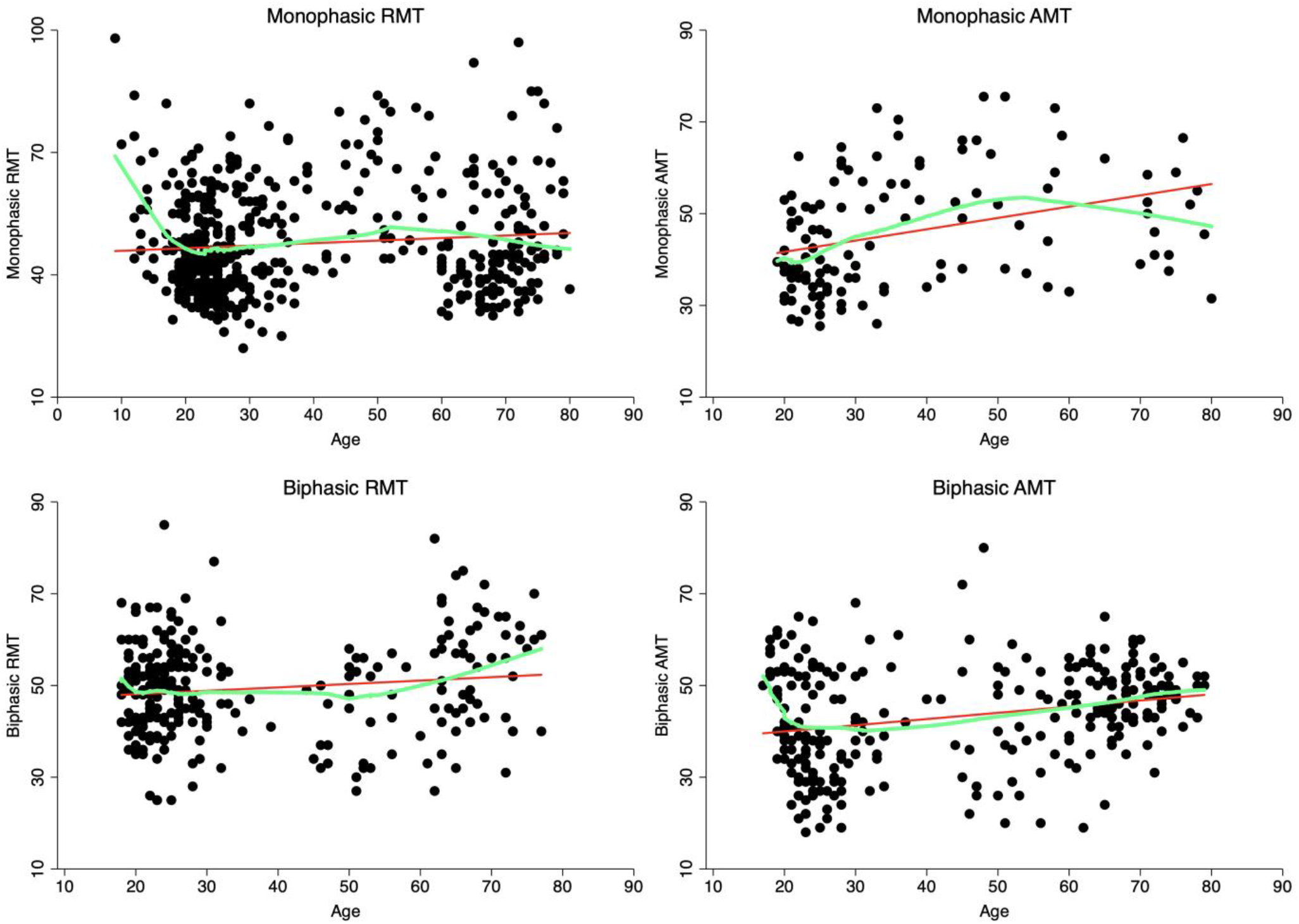
Relationship between age and motor threshold. Monophasic RMT and biphasic RMT showed a significant positive linear relationship with age (Table 6), indicating reduced corticospinal excitability in older adults. There were also significant non-linear relationships between age and monophasic AMT and biphasic AMT (see Results). Green lines fit a smoothed ‘lowess’ curve through data. Bivariate scatterplots give an indication of results only.

**Figure 9.**
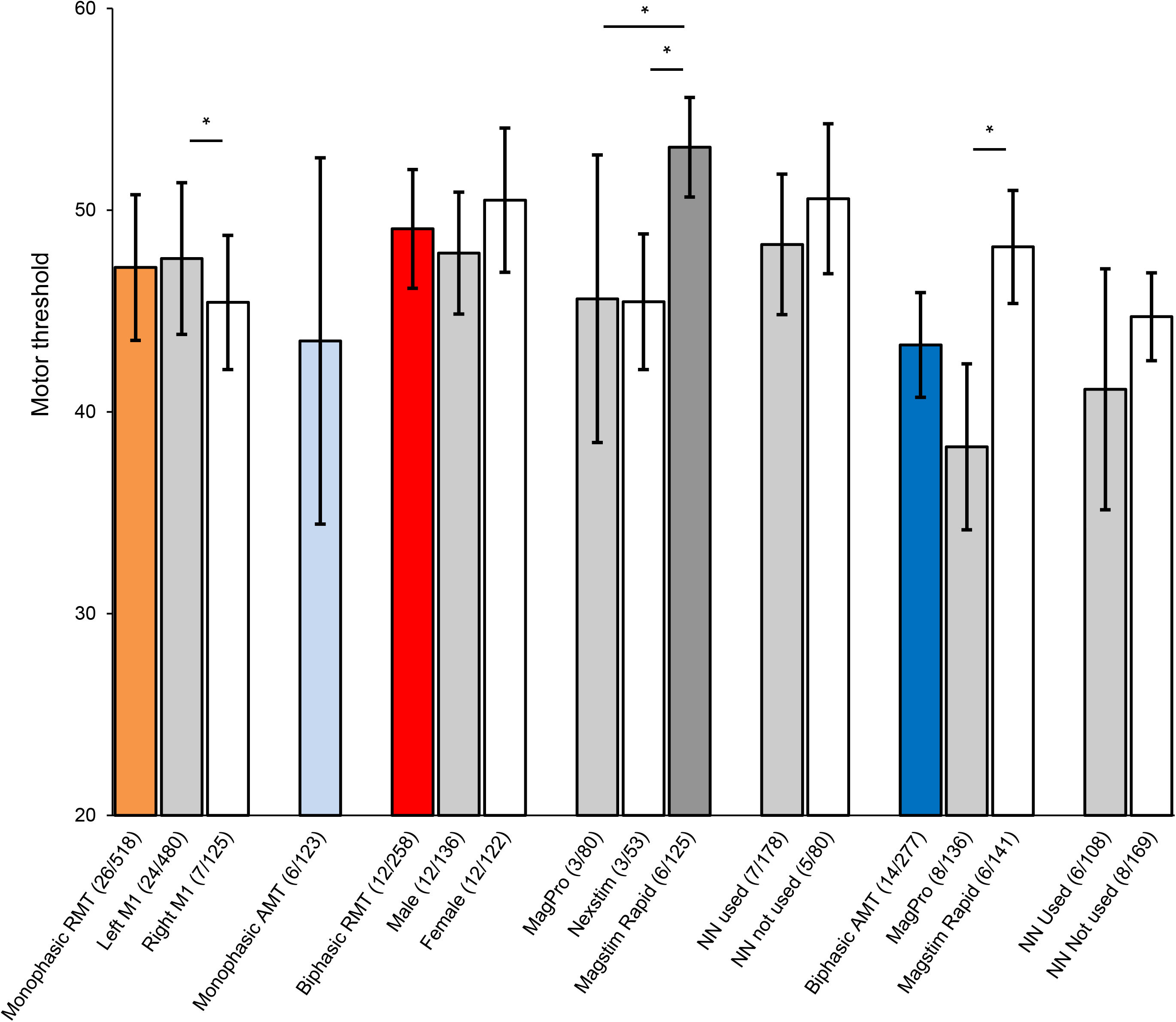
Marginal means for motor threshold. Coloured bars show overall marginal means for monophasic RMT, monophasic AMT, biphasic RMT, and biphasic AMT. Grey and white bars show marginal means of levels of the IVs M1 hemisphere, gender, TMS machine, and neuronavigation (NN), which were included in final regression models. * denotes a significant difference between levels (p < 0.05) Error bars show 95% confidence intervals. Brackets show (studies/participants).

**Table 6.**
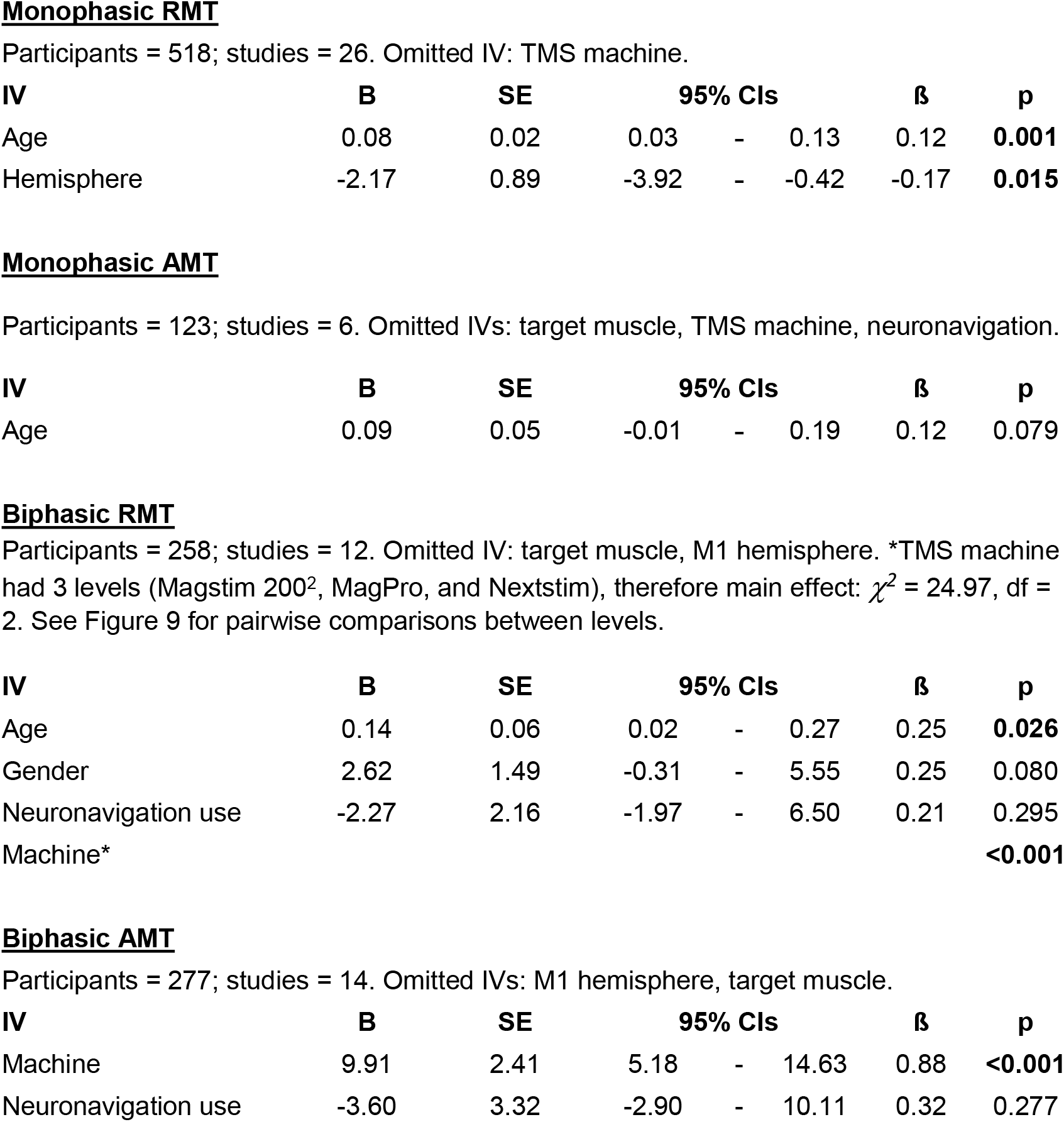
Final MT regression models. Separate analyses were conducted to investigate IVs explaining variability in each of the four types of MT. Bold denotes significance (p < 0.05). IVs omitted because of insufficient data are listed below. See Figure 9 for all IV levels.

Other IVs not included in the final regression models had p-values > 0.10 in both stage 1 and 2 regressions (see Supplementary file 10 for all stage 1 and 2 results).

### 3.9 MT post-hoc analyses

There was a significant quadratic and cubic relationship between monophasic AMT and age (p < 0.001 and p = 0.031, respectively). A cubic relationship between biphasic RMT and age did not reach significance (p = 0.070) (Figure 8).

### 3.10 Additional analyses

Two stage regression analysis demonstrated a significant difference between single pulse TMS MEP amplitudes collected using 120% of RMT, compared with those collected using the 1 mV method: 120% RMT marginal mean (studies = 17; N = 341) = 0.87 mV; 95% CIs = 0.78 – 0.96; 1 mV method marginal mean (studies = 9; N = 189) = 1.09 mV; 95% CIs = 0.97 – 1.21; B = 0.22; SE = 0.09; p = 0.015. This effect of TS intensity method was still significant when not controlling for any covariates (p = 0.013) (see Figure 1 for histograms of both methods).

Studies that employed the 120% RMT method also displayed higher average variance between participants’ MEP amplitudes: 120% RMT method studies average SD = 0.55 mV; average CV = 62.8%. 1 mV method studies average SD = 0.39 mV; average CV = 33.8%. Levene’s robust test demonstrated that the higher MEP amplitude variance for the 120% RMT method was significant (F = 23.35; df = 1, 573, p < 0.001). This lower variance for the 1mV method was expected, given that operators set the machine intensity to evoke this predefined 1mV amplitude output.

There were strong significant positive correlations between the four types of MT (all p < 0.001): monophasic RMT x biphasic RMT, N = 153, R = 0.856; monophasic RMT x monophasic AMT, N = 123, R = 0.933; monophasic RMT x biphasic AMT, N = 223, R = 0.659; biphasic RMT x biphasic AMT, N = 83, R = 0.749, monophasic AMT x biphasic AMT, N = 21, R = 0.916 (no observations for biphasic RMT x monophasic AMT).

## 4. Discussion

This study pooled data from 35 studies to demonstrate factors explaining interindividual variability in response to single and pp TMS. We suggest reasons for these observed sources of variability and propose specific methodological adjustments to reduce for their potential influence. We hope that these findings will lead to greater standardisation of single and pp TMS methods in the brain stimulation community, thereby increasing their utility as a clinical and experimental tool.

### 4.1 Baseline MEP amplitude

As in Corp et al. (2020), who applied the present method to TBS data, this study has demonstrated significant negative relationships between baseline MEP amplitude and (SICI and ICF) normalised MEP. That is, lower baseline responses resulted in higher amplitude conditioned MEPs, regardless of the pp TMS or TBS protocol. We suggest three main reasons as to why these relationships may occur in both pp TMS and TBS data (Corp et al. 2020): regression to the mean; floor and ceiling effects; and different cortical networks being probed between individuals. Regression to the mean is the statistical phenomenon by which an initial extreme measurement is more likely to be closer to the mean if measured for a second time (Bland and Altman 1994, Stigler 1997). By this logic, conditioned MEP responses are more likely to show facilitation (or ameliorated inhibition) if a person records extremely low baseline MEP amplitudes, and vice versa (Corp et al. 2020). Floor and ceiling effects occur when TMS intensities are too close to a floor (minimal activation) or ceiling (maximal activation of neurons), and thus further inputs fail to produce discernible changes in MEP amplitude (Devanne et al. 1997). While TS intensities are individualised, usually to 120% RMT or a 1 mV value, there can be substantial variability in relation to where these stimulus intensities occur in relation to each individual’s input/output curve (Goldsworthy et al. 2016b, Houdayer et al. 2008, Pitcher et al. 2015). In other words, these individualised TS intensities can be a relatively low or high between individuals. This can bias the effects of the CS, with ‘inhibition’ less likely for individuals with low relative TS intensities, and ‘facilitation’ less likely for those with high relative TS intensities (Amandusson et al. 2017, Goldsworthy et al. 2016b). If we assume that those with low baseline MEP amplitudes received TMS pulses at relatively low intensities, this would agree with the negative relationship in the present study, where low baseline MEP amplitudes resulted in greater ICF effects yet ameliorated SICI effects (Figures 4 & 6). However, this is speculative given that we could not directly assess the relative stimulus intensities at which the pulses were applied. Lastly, it has been shown that TS intensity influences the cortical circuits activated by the TMS pulse (Di Lazzaro et al. 1998). Thus, if the TS intensity used for an individual does not probe the circuits activated by the initial CS, SICI and ICF may not be revealed (Di Lazzaro et al. 1998, Garry and Thomson 2009). Based on this, the negative relationship for baseline MEP amplitude in the present study may suggest that SICI is best probed by high relative TS intensities and ICF best probed by low relative TS intensities. However, this does not agree with previous research showing that SICI and ICF are maximal at moderate TS intensities (Cosentino et al. 2018, Garry and Thomson 2009). This suggests that regression to the mean and floor and ceiling effects may have been stronger influences on SICI and ICF response, however again this is speculative, given that we could not directly test the relative intensities at which the pulses were applied within individuals.

### 4.2 Motor threshold predicts single and paired-pulse TMS response

Our data demonstrated that MT predicted single pulse MEP amplitude, SICI, and ICF response. For single pulse TMS, this is in agreement with Peterchev et al. (2013), who showed that individuals with lower MTs have steeper I/O slopes (Peterchev et al. 2013). We demonstrate a similar result here by showing that individuals with lower MTs have higher MEP amplitudes at one stimulus intensity along the I/O curve (120% RMT). For SICI and ICF, this phenomenon may be in part caused by the fact that the conditioning stimulus intensity (as a percentage of the machine output) is adjusted to an individual’s MT. This is designed to ensure the activation of a similar proportion of corticospinal neurons between individuals. However, SICI and ICF mechanisms are dependent on *intracortical*, rather than *corticospinal* neurons, and the threshold for activation of these two networks does not necessarily correlate (Chen et al. 1998). Thus, those with higher MTs receive a higher intensity CS (as a percentage of machine output), and this could cause stronger activation of intracortical mechanisms (Amandusson et al. 2017) (and thus an increased SICI and ICF effect, as demonstrated here). However, these relationships could also be caused by inherent differences in SICI and ICF for individuals with low or high MTs, with the differential effects of stimulus intensity and MT unable to be disentangled here due to machine output being adjusted to MT in all studies.

### 4.3 Effect of age on corticospinal excitability

Linear regression showed that, on average, monophasic RMT and biphasic AMT significantly increased with age. However, this reduction in corticospinal excitability does not appear to be linear across the lifespan, demonstrated by significant quadratic relationships for monophasic AMT, and biphasic AMT, and fitted ‘lowess’ lines through MT data indicating curved patterns at particular age points (Figure 8). These fitted lines suggest an initial stage of hypoexcitability for people under ∼20 years of age, with MT then reaching its lowest point at about the age of 25. After this age, there seemed to be different patterns in monophasic and biphasic data, with monophasic MTs increasing through middle age, then reducing again in older age, as opposed to biphasic MTs - which continued to increase with age. The divergent patterns observed in monophasic and biphasic data could be due to different cortical mechanisms activated by these pulse waveforms; biphasic pulses may activate later I-waves compared to monophasic posterior-anterior stimulation (Di Lazzaro et al. 2001). However, the pattern of activation may also depend on stimulus intensity, and the initial current direction of the biphasic pulse (Di Lazzaro et al. 2001), for which we had incomplete information. The curved pattern of response for monophasic MTs is similar to that of Shibuya et al. (2016), who demonstrated the lowest monophasic RMTs for 20-25 year olds and older adults (study age range: 20-83), and maximal RMT at approximately 50 years of age, and a significant quadratic effect.

Interestingly, the higher monophasic RMT for < 20 year olds (Figure 8) did not translate to a significant quadratic or cubic effect. This may be because the majority of these observations came from one study (Croarkin et al. 2013), and these values would have been adjusted given that we included ‘study ID’ as a random variable to account for the fact that data came from different studies. However, the relationships between corticospinal excitability and age observed in the present study should be interpreted with caution given the relative dearth of data for adolescents and middle-aged adults (Figure 7).

### 4.4 Effect of hemisphere on cortical excitability

Our results demonstrated reduced SICI and increased monophasic RMT in the left hemisphere. These effects were similar when including only data from right handers from our restricted sample for which we had handedness data (although the effect became non-significant for monophasic RMT, p = 0.079). Thus, while we do observe these effects in right handers, we cannot say whether they are driven by the fact that the left hemisphere is the dominant M1, or whether it is simply an effect of the left hemisphere across both right and left handers. The collection of additional data from left handers will be required to answer this question. In regards to previous literature, Ilic et al. (2004), also showed reduced SICI in the left M1 in right handed participants. These authors suggested that less SICI in the dominant hemisphere for right handers may provide an advantage for the readiness and ease to carry out movements with the dominant hand (Ilic et al. 2004). In contrast, our monophasic RMT findings differ to Ilic et al. (2004), who showed *reduced* monophasic RMT in the left hemisphere for right handers. It is not clear as to why we obtained conflicting MT results. However, given our non-significant results when only including right handers, and the small sample size of Ilic et al. (2004) (9 right handers), these effects are not conclusive, and additional hemisphere and handedness data needs to be gathered.

### 4.5 Effect of machine on corticospinal excitability

We found that Nexstim machines were more powerful than MagPro machines for single pulse MEP amplitude, yet observed higher biphasic RMT and biphasic AMT for the Magstim Rapid machine than MagPro and Nexstim machines. Much of this effect is likely due to the use of Magstim Rapid machines for biphasic MT assessment prior to repetitive TMS protocols (delivered with biphasic pulses), which have a reduced power output in comparison to Magstim 200^2^ (Kammer et al. 2001), and MagPro X100 machines (Koponen et al. 2020). These differential effects highlight the importance of the inclusion of TMS machine (and study location if applicable) as a covariate in statistical analyses on data that are pooled collaboratively using different machines. Researchers should also be aware that the various configurations of the Magstim BiStim machine (i.e. two connected Magstim 200^2^ machines) produce different power outputs, which may confound electrophysiological results if configured incorrectly (Do et al. 2019). We did not collect information on these configurations in the present study, which may have affected results.

### 4.6 Limitations

A number of limitations should be acknowledged. First, we were limited to analysing the variables that were available to us, and so could not measure the impact of IVs such as menstrual cycle (Hattemer et al. 2007), or neuroimaging markers (Silbert et al. 2006) on corticospinal excitability. Second, our approach pooled data from separate studies, and thus does not have the precision of a repeated-measures design. Pooling different studies’ results increases the risk of between-study variability being caused by factors such as sampling error, study setting, and experimenter behaviour (Higgins and Green 2011). Next, of the nine studies using neuronavigation, none reported coordinates of the motor hotspot, nor coil shift data from the motor hotspot. Thus, unaccounted for differences in coil position may have explained some unobserved intraindividual variability in TMS outcomes. Next, we were limited by the incomplete dataset that we could gather for handedness, and also the small number of left-handers within that dataset. Thus, we do not know whether our ‘hemisphere’ effects were driven by hemispheric differences between left and right handers, or by handedness. Next, we did not measure the potential impact of TMS machine coil size or type, or initial waveform direction (i.e. AP or PA), on cortical excitability. Finally, it should be acknowledged that a portion of interindividual variability in MEP amplitudes occurs due to differences in the excitability of spinal circuits (Kiers et al. 1993, Lackmy and Marchand-Pauvert 2010), and we could not account for this given that the included studies did not measure sub-cortical responses such as the M-max or H-reflex.

### 4.7 Recommendations

We first propose some steps to counter the significant relationships observed between baseline MEP amplitude and SICI/ICF. To avoid regression to the mean caused by chance occurrences of high or low MEP amplitudes, we recommend that investigators: 1) collect a sufficient number (20-30) of MEPs in their TMS blocks (Chang et al. 2016, Goldsworthy et al. 2016a); 2) avoid possible initial states of hyperexcitability within TMS sessions (Brasil-Neto et al. 1994, Schmidt et al. 2009); and 3) include baseline MEP amplitude as a covariate in statistical analyses. To avoid floor and ceiling effects, the CS could be normalised to 50% of maximal inhibition/facilitation (McAllister et al. 2009), while the TS could be normalised to 50% of maximal MEP amplitude (Goldsworthy et al. 2016b, Houdayer et al. 2008). This would also circumvent the aforementioned issues with normalising the CS to MT (Chen et al. 1998). However, it has previously been suggested that the use of this TS intensity may still result in substantial between-subject differences in the in the neural circuits probed by the TMS pulse (i.e. relative D- and I-wave contributions to the MEP) (Goldsworthy et al. 2016b). Until this can be empirically investigated (most likely through recordings from the cervical epidural space, e.g. Di Lazzaro et al. (2001)), we recommend that researchers minimise the aforementioned biases by collecting data across a range of stimulus intensities (i.e. pp input/output curves) (Ilić et al. 2002, Orth et al. 2003). However, in addition to the increased complexity in analysing pp input/output curve data, their collection is time consuming, especially if varying both CS and TS intensities. Thus, further effort should be directed towards the formulation of time effective methods of collection of (single and) pp TMS curve data, and increased standardisation in their analysis. Next, in order to reduce possible variability due to coil position, we suggest that where neuronavigation can be used, researchers should report the coordinates of the motor hotspot, and report or analyse the impact of shifts from the motor hotspot for individual participants. Lastly, when making age comparisons, investigators should be aware that the relationship between age and corticospinal excitability may not be linear across the lifespan.

### 4.8 Conclusions

The present study pooled individual participant data across 35 studies to demonstrate sources of interindividual variability in single and pp TMS measurements, including baseline MEP amplitude, age, TS intensity, M1 hemisphere, ISI, TMS machine, and MT. We have highlighted possible reasons for these sources of variability and made specific methodological recommendations to reduce their influence. These findings highlight the need for increased standardisation of single and pp TMS methods across the brain stimulation community, which we hope will be facilitated through this collaborative approach. We are currently expanding the ‘Big TMS Data Collaboration’ through the construction of an individual participant TMS data repository at www.bigtmsdata.com, and welcome additional brain stimulation researchers to contribute to this database.

## Abbreviations and nomenclature

TMS: Transcranial magnetic stimulation
MEP: motor evoked potential
pp: paired-pulse
SICI: short-interval intracortical inhibition
ICF: intracortical facilitation
IV: independent variable
DV: dependent variable
Normalised MEP: DV for SICI and ICF analyses (conditioned MEP amplitude expressed as a percentage of the baseline MEP amplitude)
CS: conditioning stimulus (initial pulse for paired-pulse TMS protocols)
TS: test stimulus (second pulse for pp TMS protocols, or unconditioned / baseline MEPs for pp protoocol)
ISI: interstimulus interval
RMT: resting motor threshold
AMT: active motor threshold
Pulse waveform: monophasic or biphasic pulse waveforms

## Acknowledgments

We would like to thank all of the researchers who were kind enough to share the data that they worked so hard to collect.

## Collaboration members

Daniel T. Corp, Hannah G. K. Bereznicki, Gillian M. Clark, George J. Youssef, Peter J. Fried, Ali Jannati, Charlotte B. Davies, Joyce Gomes-Osman, Julie Stamm, Sung Wook Chung, Steven J. Bowe, Nigel C. Rogasch, Paul B. Fitzgerald, Giacomo Koch, Vincenzo Di Lazzaro, Alvaro Pascual-Leone, Peter G. Enticott, Pamela Barhoun, Natalia Albein-Urios, Melissa Kirkovski, Lysianne Beynel, Hannah J. Block, Filippo Brighina, Pierpaolo Busan, Fioravante Capone, Benjamin W. X. Chong, Guiseppe Cosentino, Paul E. Croarkin, Zafiris J. Daskalakis, Daina S. E. Dickins, Michele Dileone, Michael Do, Luciano Fadiga, Anna Fuhl, Mitchell R. Goldsworthy, Christian Grefkes, Fabian Helm, Najmeh Hoseini, Annapoorna Kuppuswamy, Cheng-Ta Li, Wei-Chen Lin, Caroline Lucke, Christian Marendaz, Alessandro Martorana, Michelle N. McDonnell, Moniek A. M. Munneke, Kate Murdoch, Charlotte Nettekoven, George M. Opie, Paolo Profice, Rohan Puri, Federico Ranieri, Michael C. Ridding, Stephan Riek, John G. Semmler, Amaya M Singh, W Richard Staines, Dick F. Stegeman, Cathy M. Stinear, Jeffery J. Summers, Pietro A. Tonali, François Tremblay, Liam J. Wallace, Nick S. Ward, Ann-Maree Vallence, Marine Vernet, Woo-Kyoung Yoo, Marielle Young-Bernier, Juho Joutsa, Andris Cerins, Ulf Ziemann, Alison Canty, Mark R. Hinder, Timothy P. Morris, Christian Hyde, Wolnei Caumo

---

**Supplementary file 2. Search syntax.**

Search ((intermittent theta-burst stimulation OR intermittent theta burst stimulation OR iTBS)) AND (Transcranial magnetic stimulation OR TMS) Filters: Publication date from 2013/01/01 to 2016/12/31. Results = 126

((continuous theta-burst stimulation OR continuous theta burst stimulation OR cTBS)) AND (Transcranial magnetic stimulation OR TMS) Filters: Publication date from 2012/01/01 to 2016/12/31 Results = 239

((short-interval intracortical inhibition OR short interval intracortical inhibition OR SICI)) AND (Transcranial magnetic stimulation OR TMS) Filters: Publication date from 2014/01/01 to 2016/12/31. Results = 218

((intracortical facilitation OR ICF)) AND (Transcranial magnetic stimulation OR TMS) Filters: Publication date from 2014/01/01 to 2016/12/31. Results = 152

((input-output curve* OR stimulus-response curve* OR I-O curve* OR IO curve* OR S-R curve* OR SR curve*)) AND (Transcranial magnetic stimulation OR TMS) Filters: Publication date from 2013/01/01 to 2016/12/31. Results = 69

Supplementary file 3. Distribution plots. Histograms show distribution of MEP data for single pulse, SICI and ICF protocols, prior to outlier winsorization.

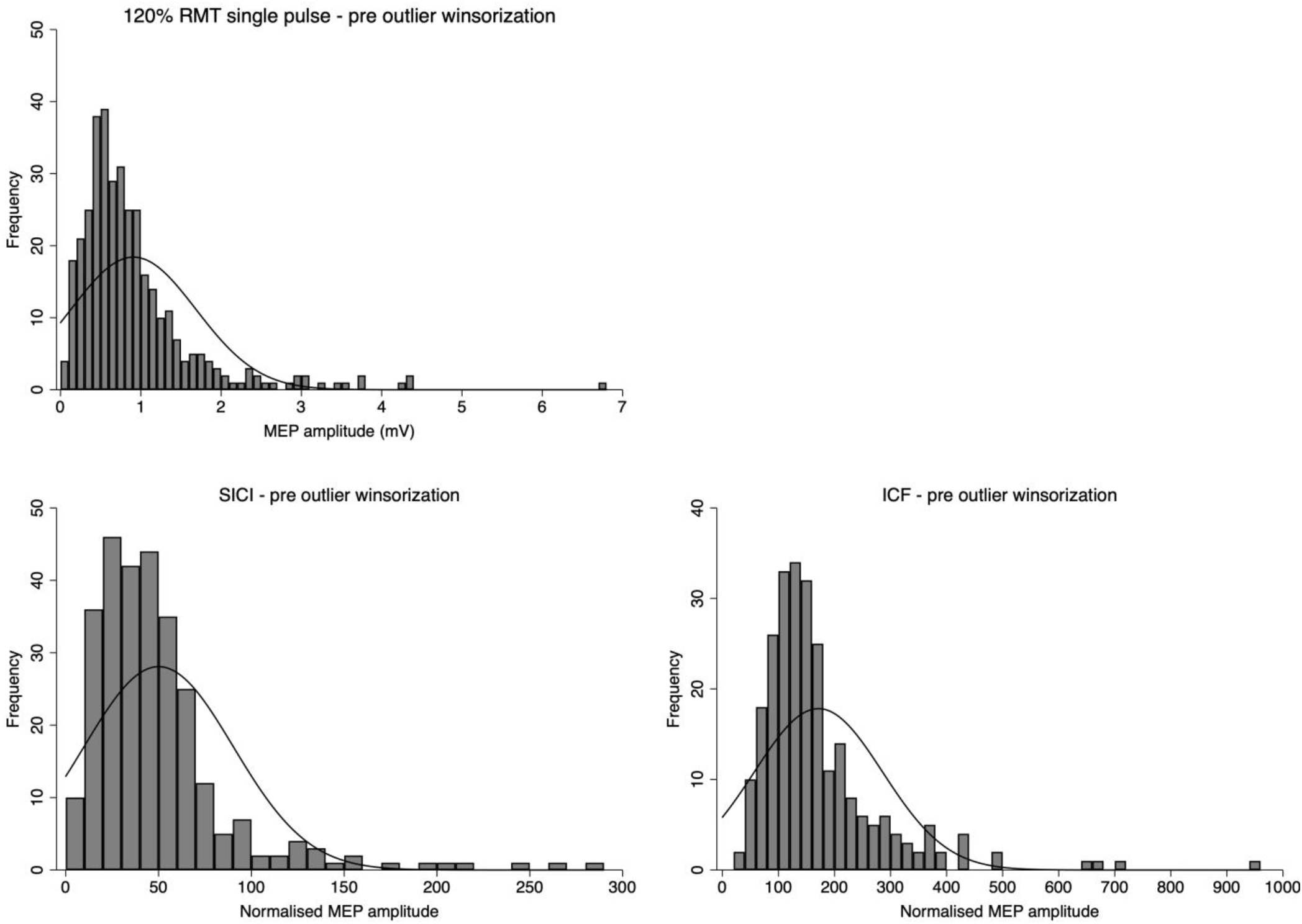

Supplementary file 4: Reproducibility data from Beynel et al. (2014)

## Methods

Test-retest data were taken from 35 healthy participants (19 females; mean age: 44.67 ± 20.12) at a month interval. Single pulse MEP data were assessed at 120% of RMT, while SICI and ICF were assessed at 80% and 120% of RMT, for conditioning and test stimuli, respectively, with interstimulus intervals of 2 ms (SICI) and 15 ms (ICF). Ten MEPs were collected per condition, per session. Please see the published study (Beynel et al., 2014) for further methodological details. As in the main manuscript (Corp et al.), for SICI and ICF, each individual’s mean conditioned MEP amplitude was normalised to their mean baseline MEP amplitude.

## Results

The intraclass correlation coefficients (McGraw et al., 1996) for each TMS protocol were as follows: biphasic RMT = 0.845; single pulse MEP amplitude = 0.375; ICF = 0.376; and SICI = 0.367.

Beynel L, Chauvin A, Guyader N, Harquel S, Marendaz C. Age-related changes in intracortical inhibition are mental-cognitive state-dependent. Biol Psychol 2014; 101: 9-12.

McGraw KO, Wong S. Forming inferences about some intraclass correlation coefficients. Psychological methods 1996; 1: 30.

**Supplementary file 5.** The use of z-scores grouped by study to run correlation analyses. Table shows an example of this method, using the correlations between monophasic RMT and biphasic RMT.

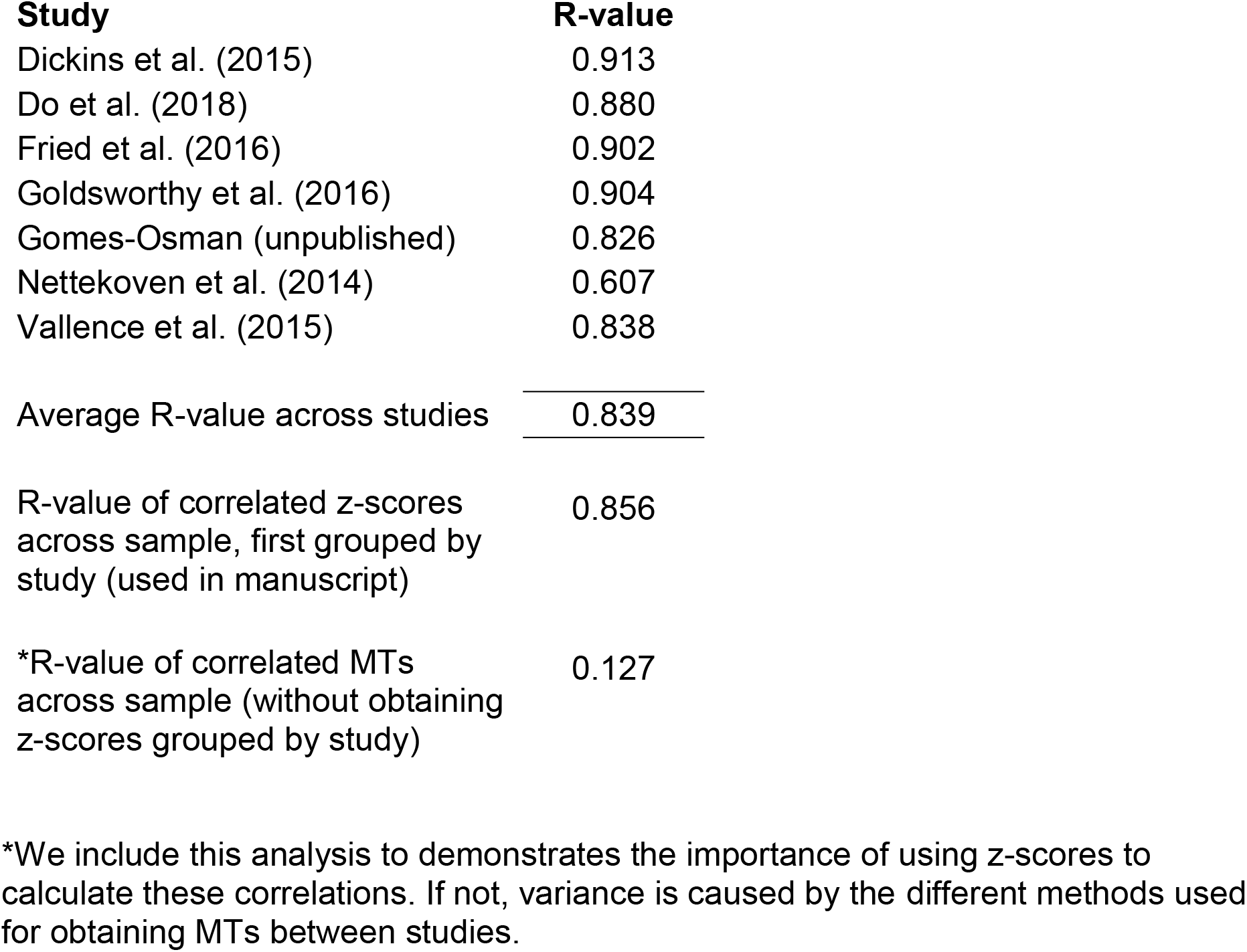

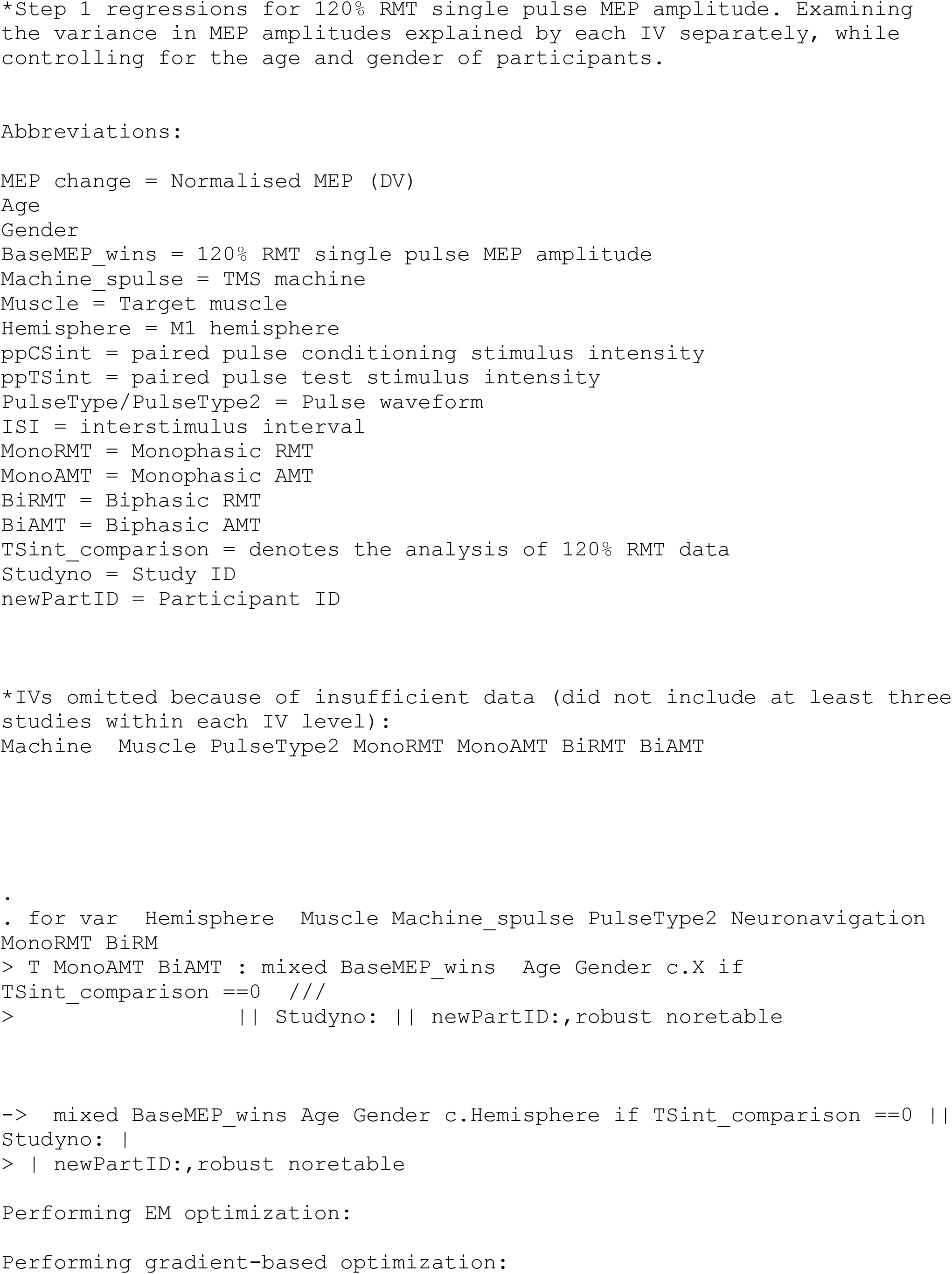
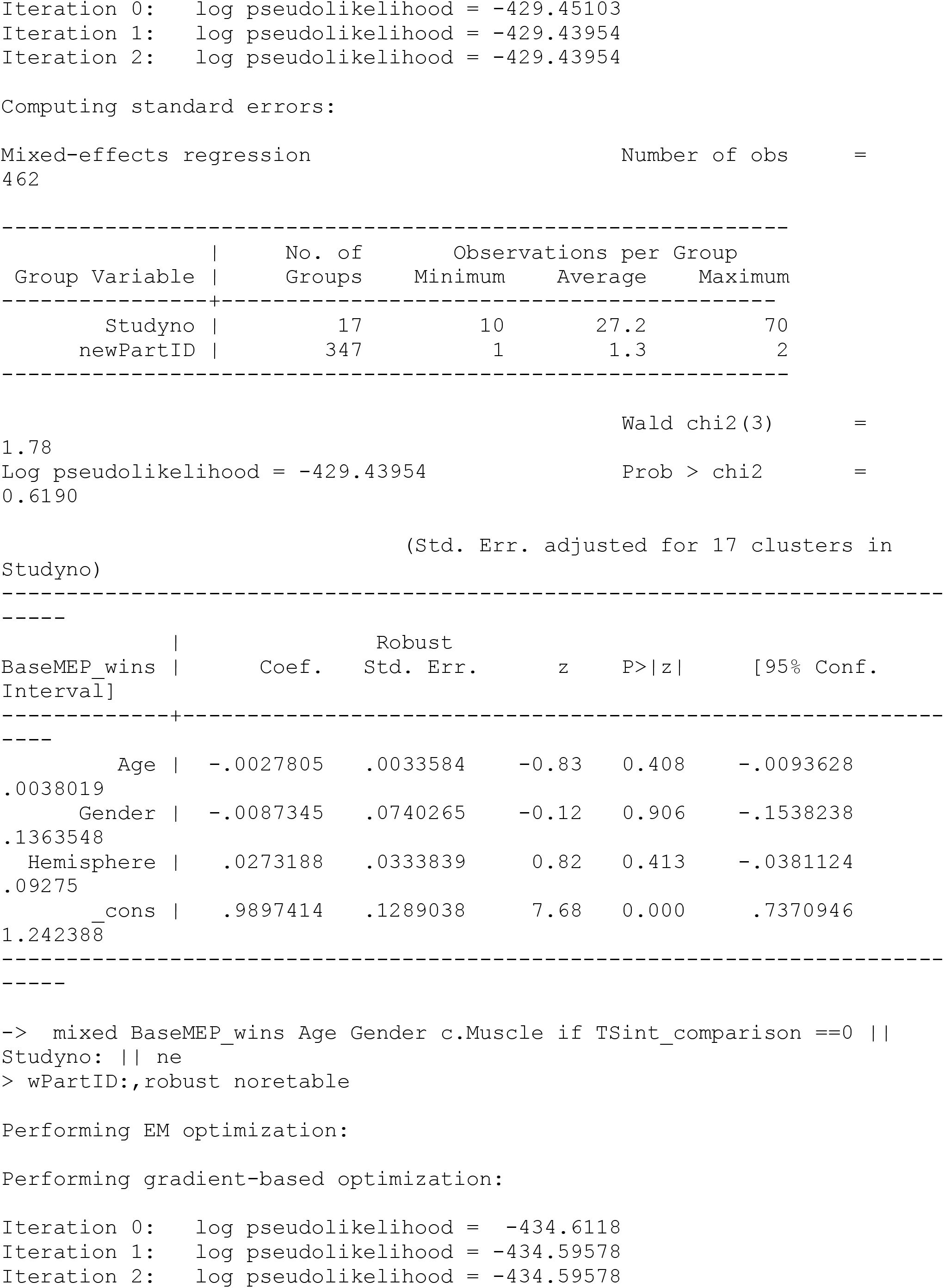
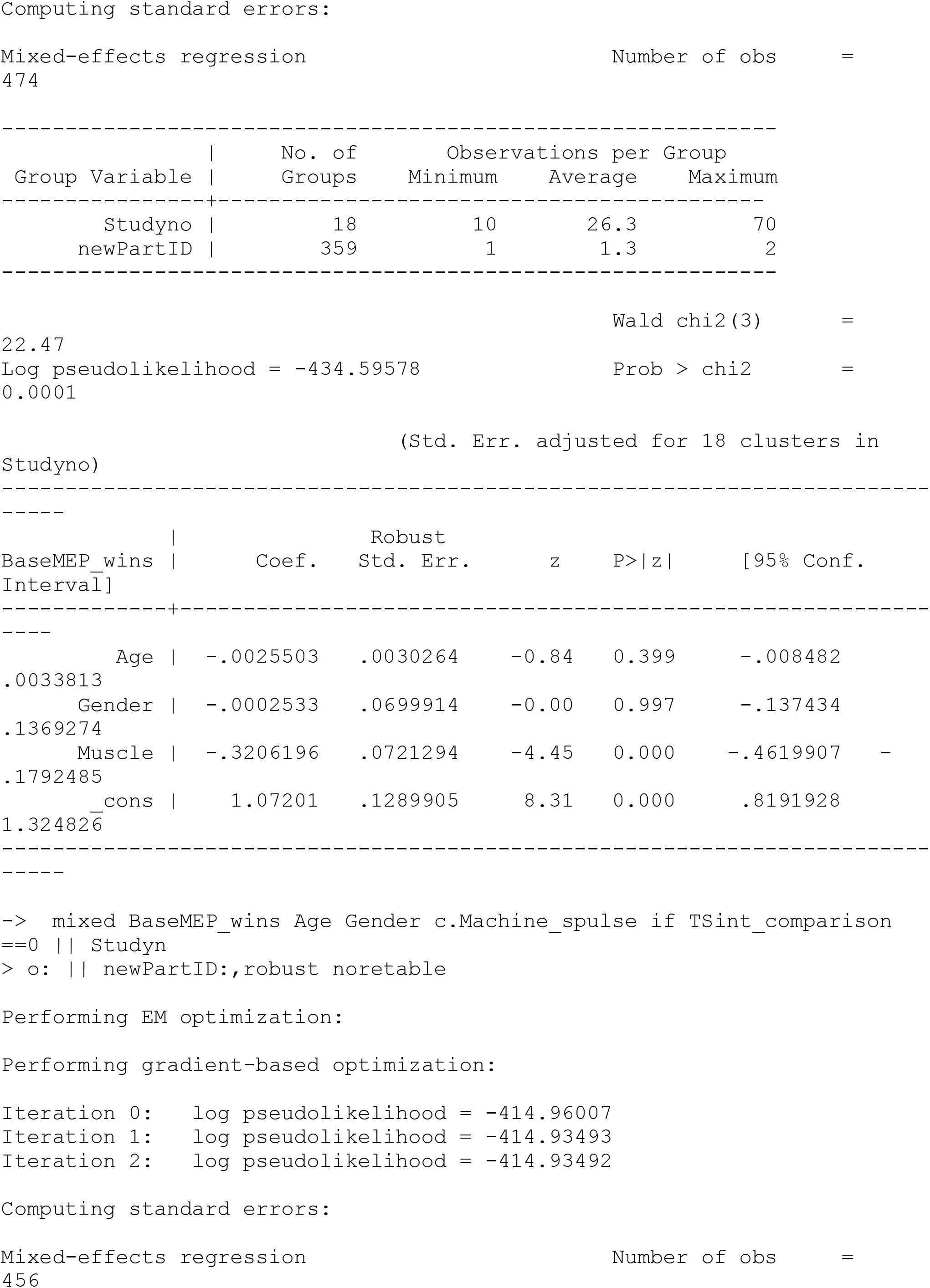
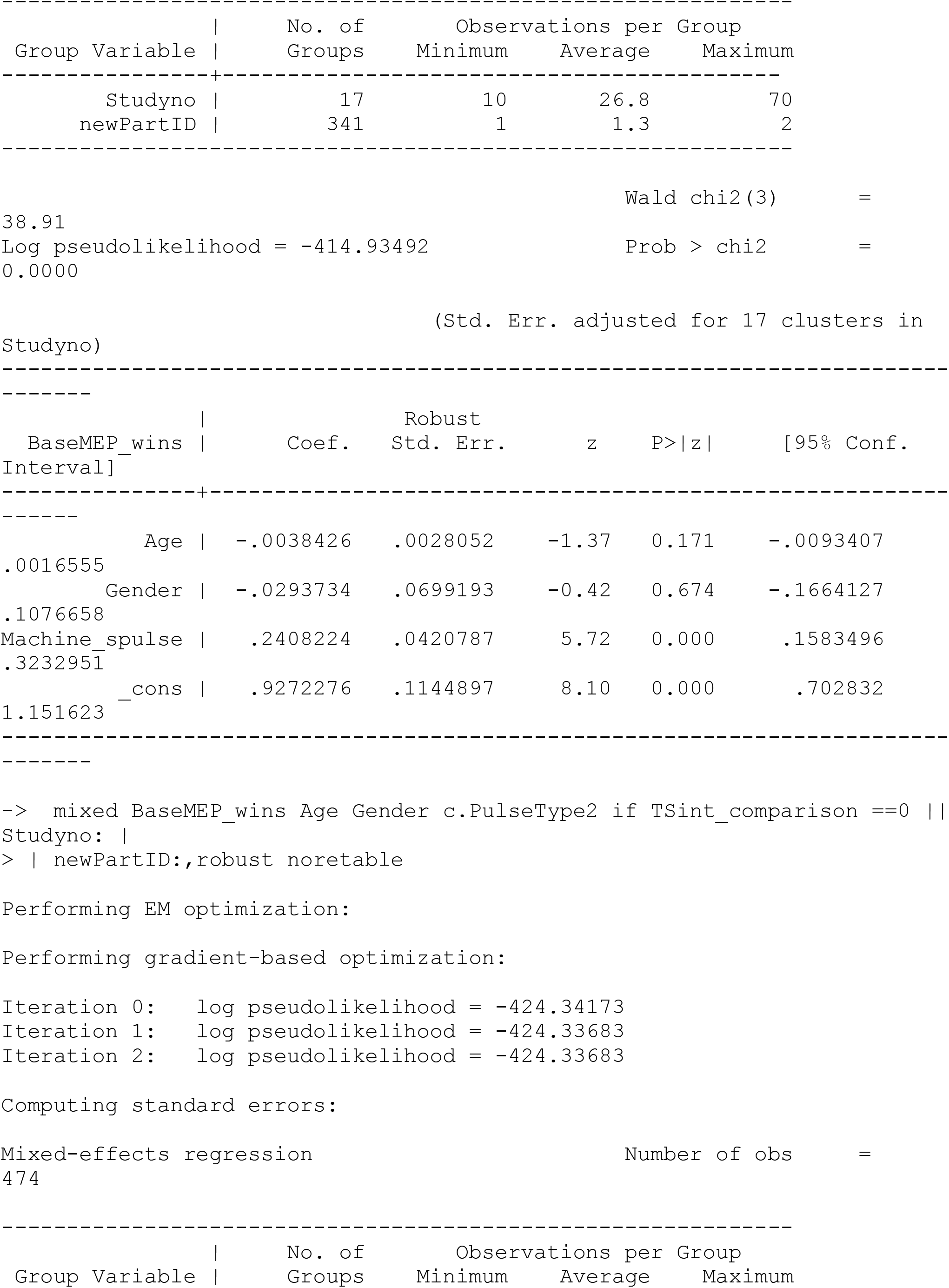
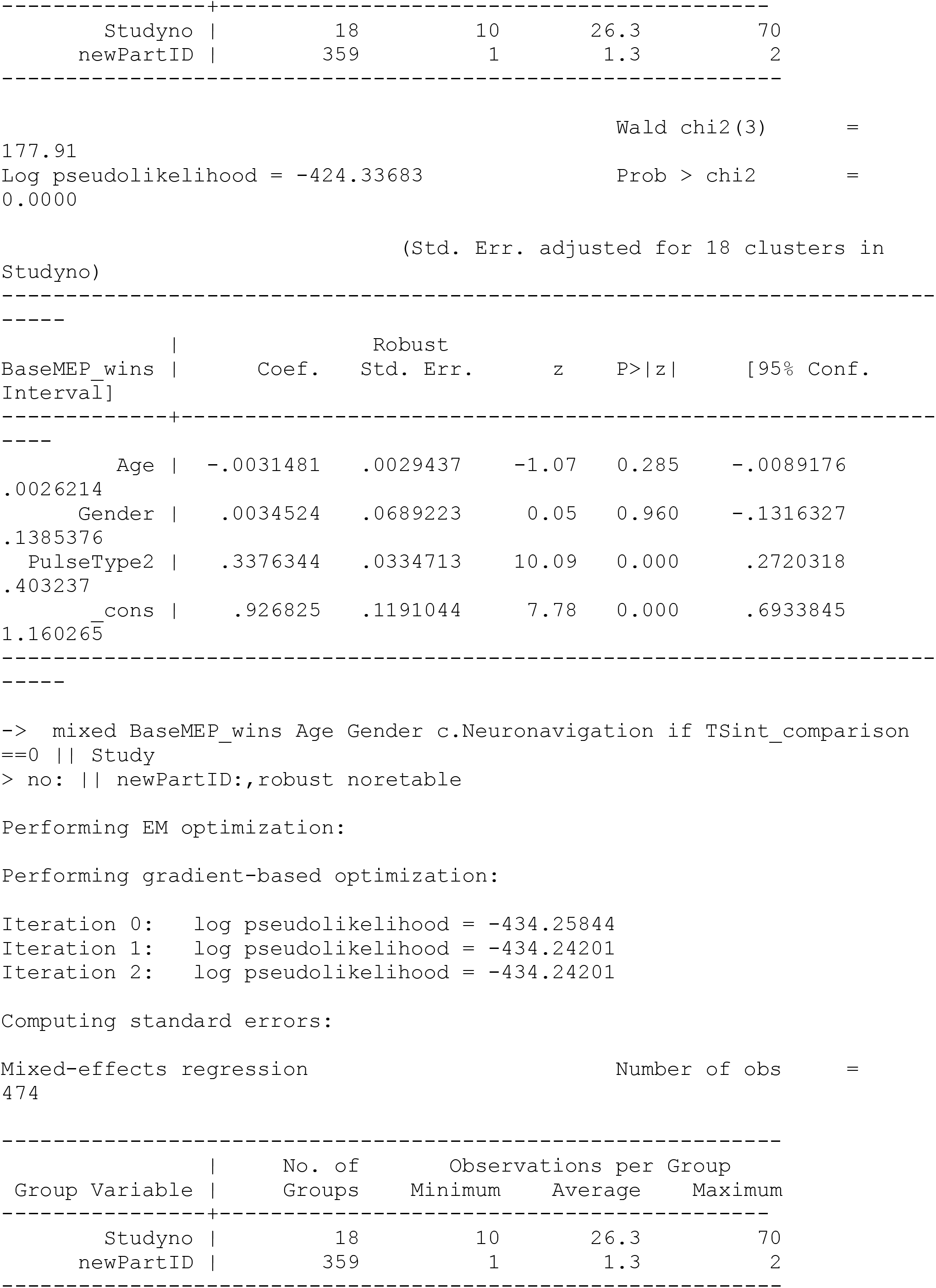
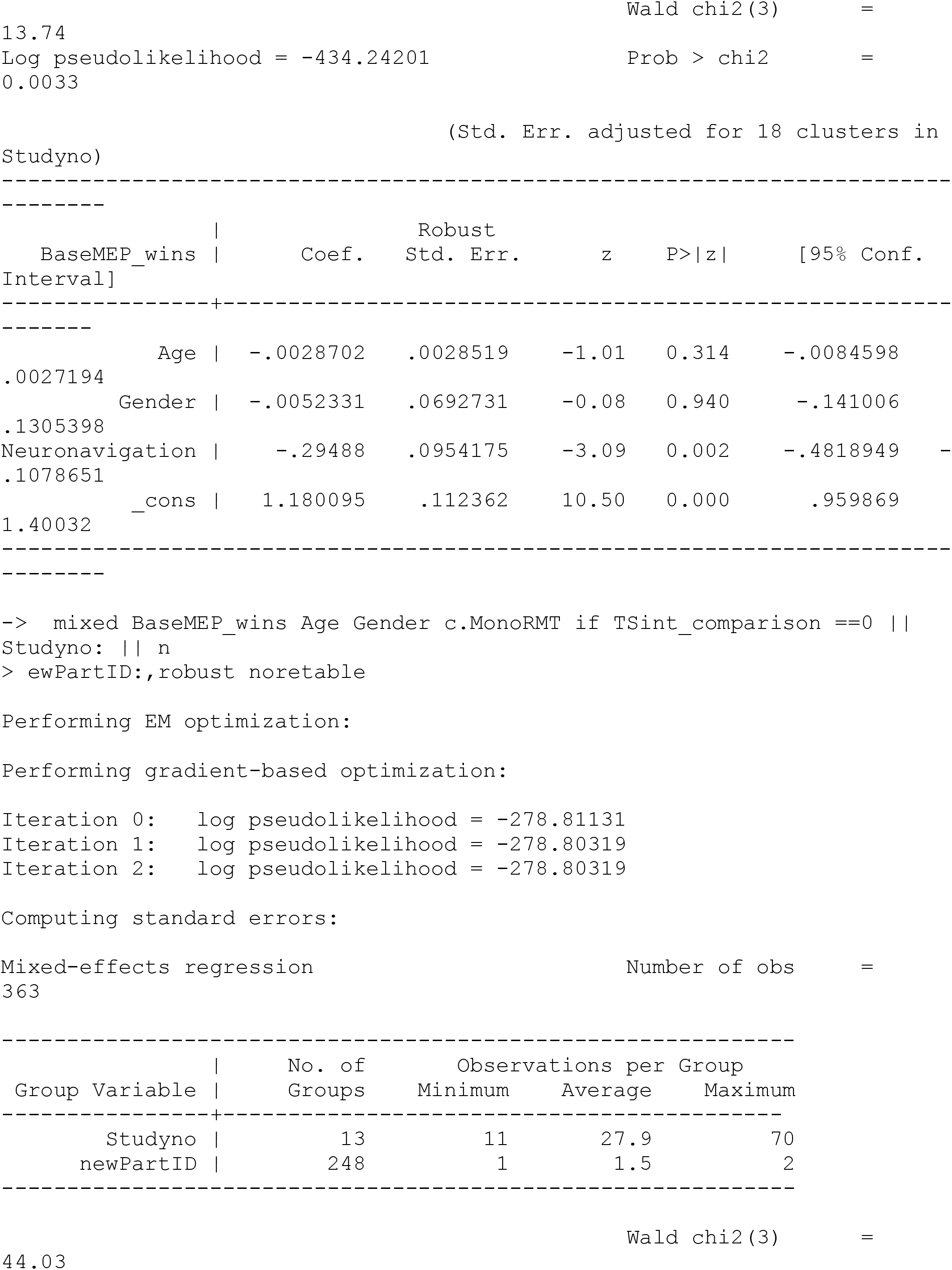
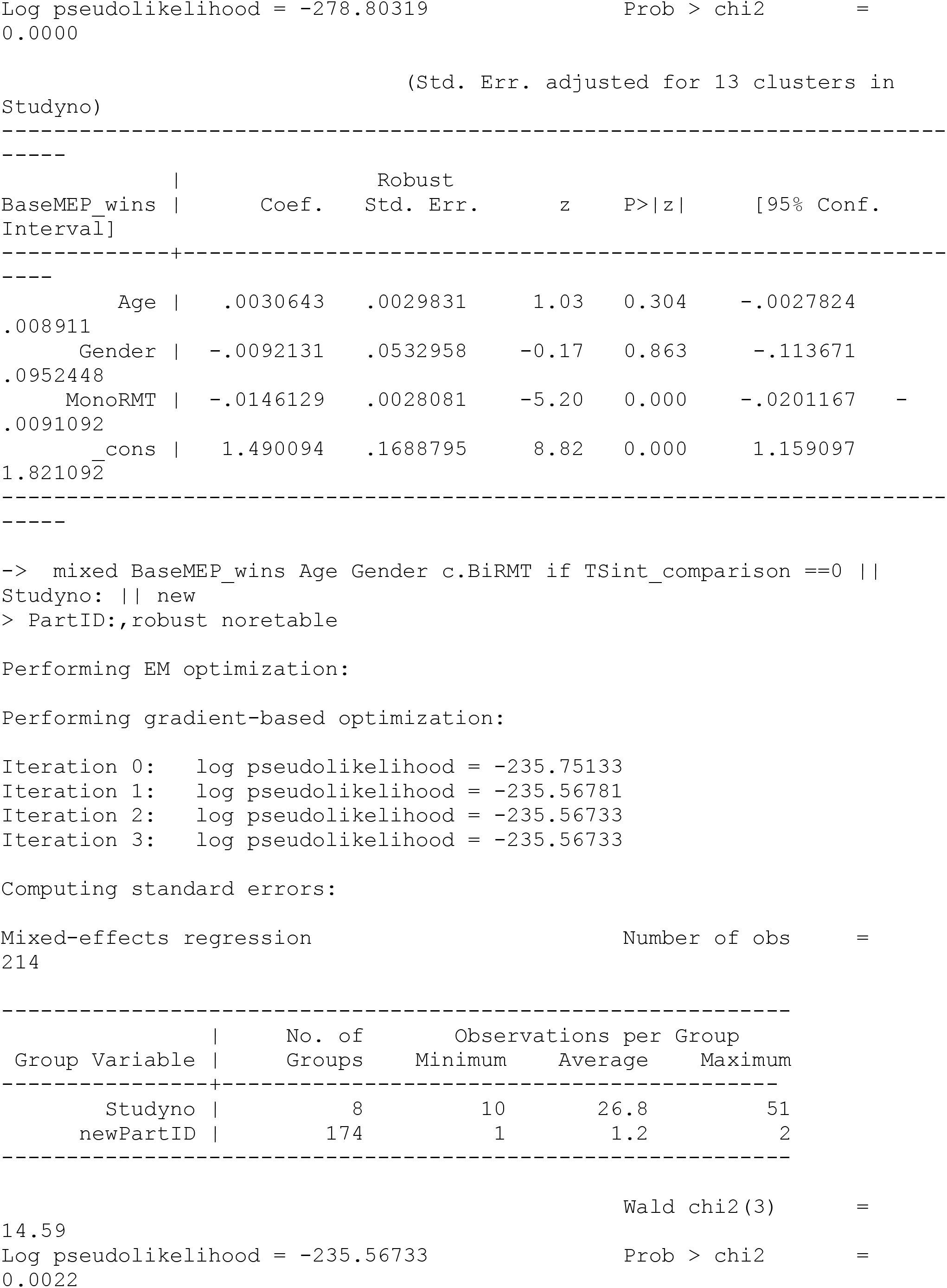
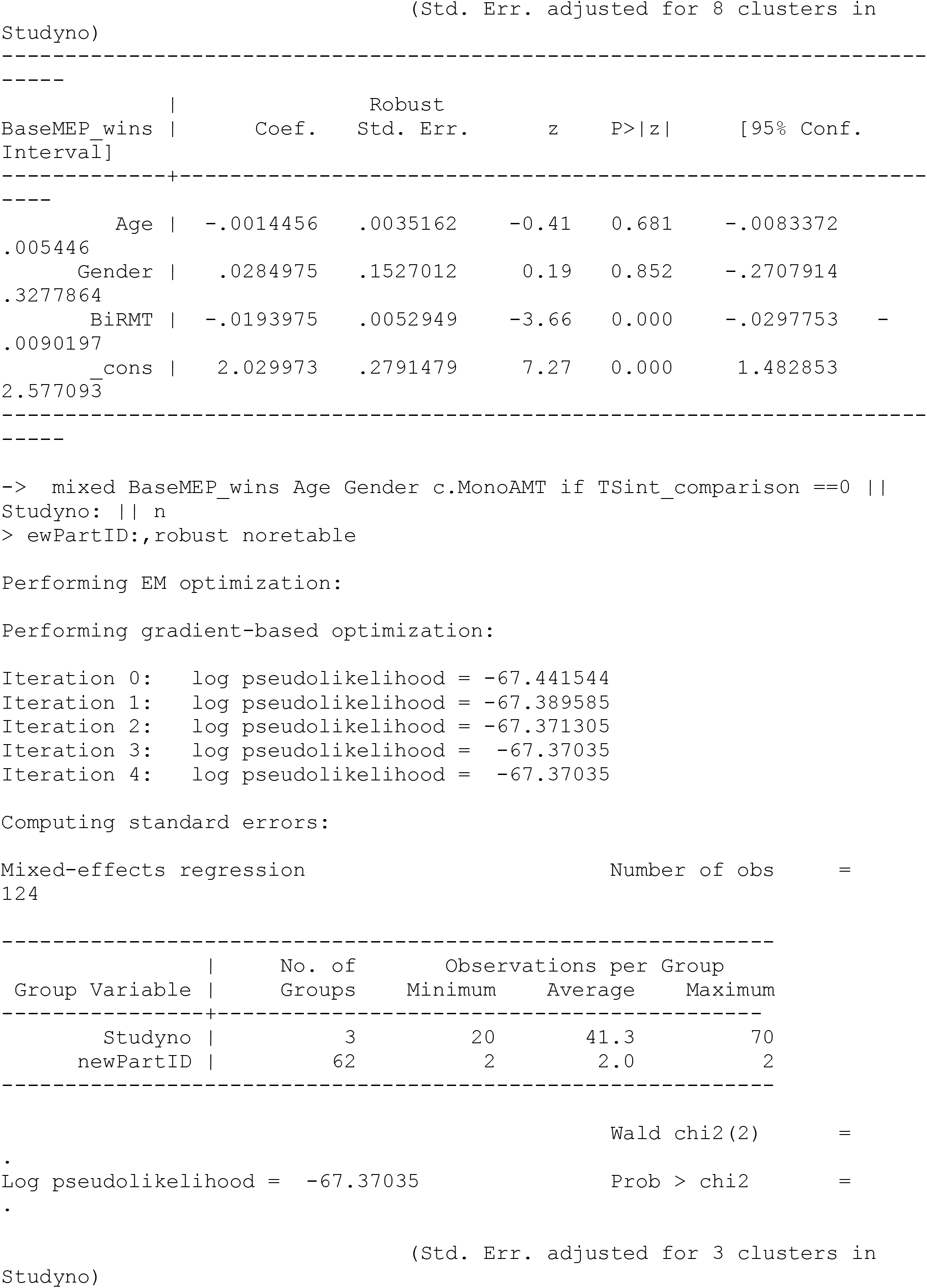
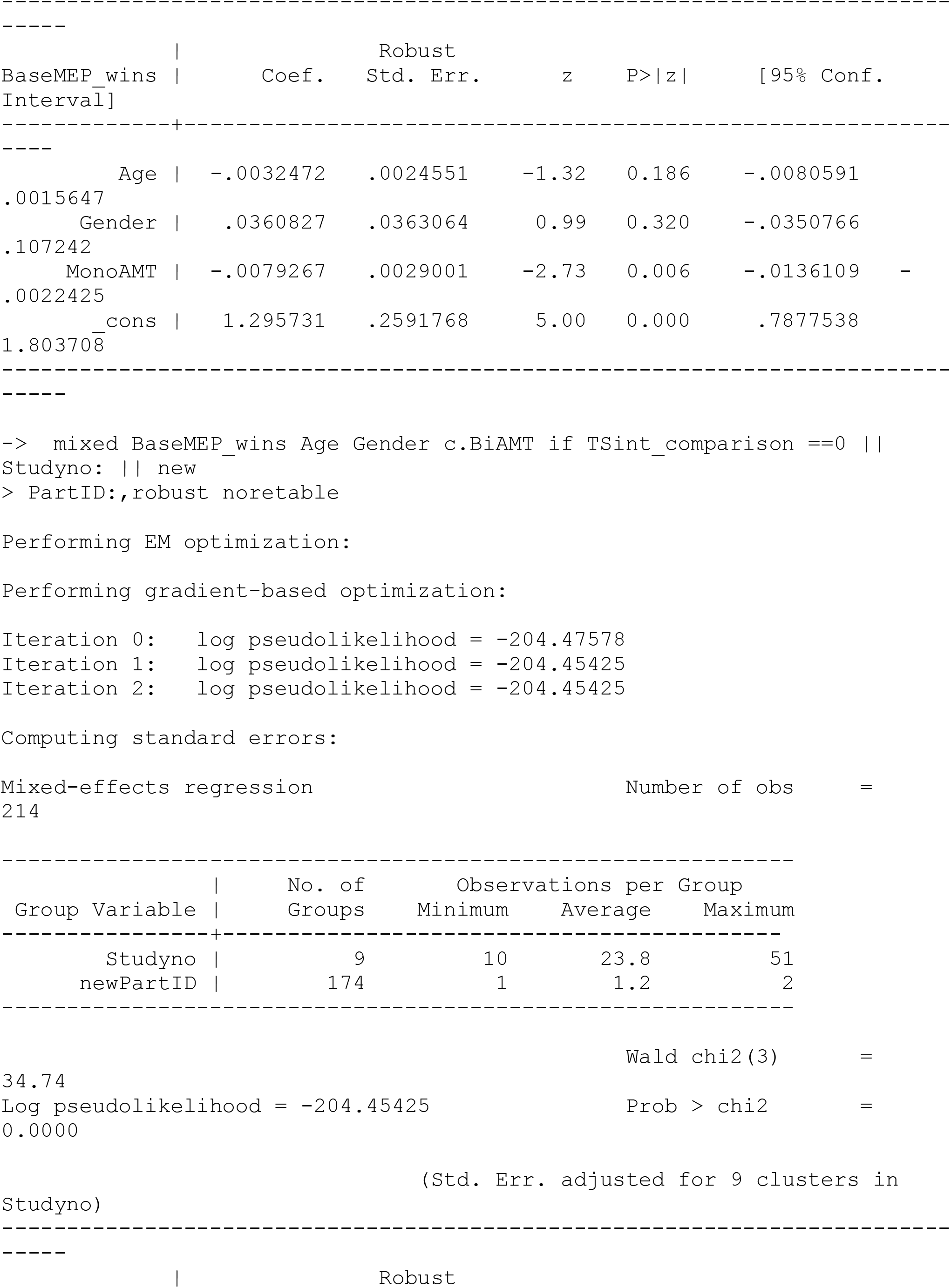
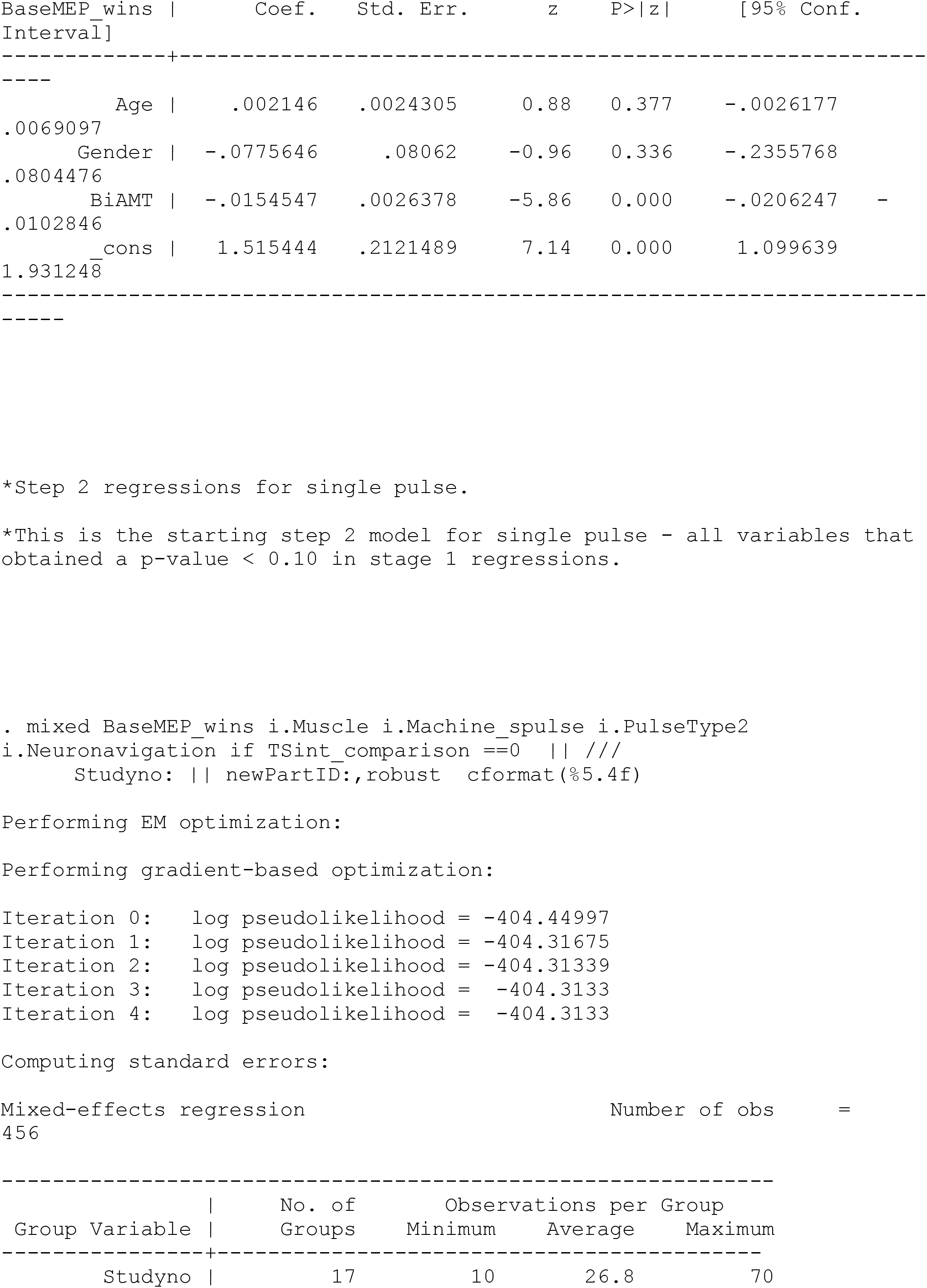
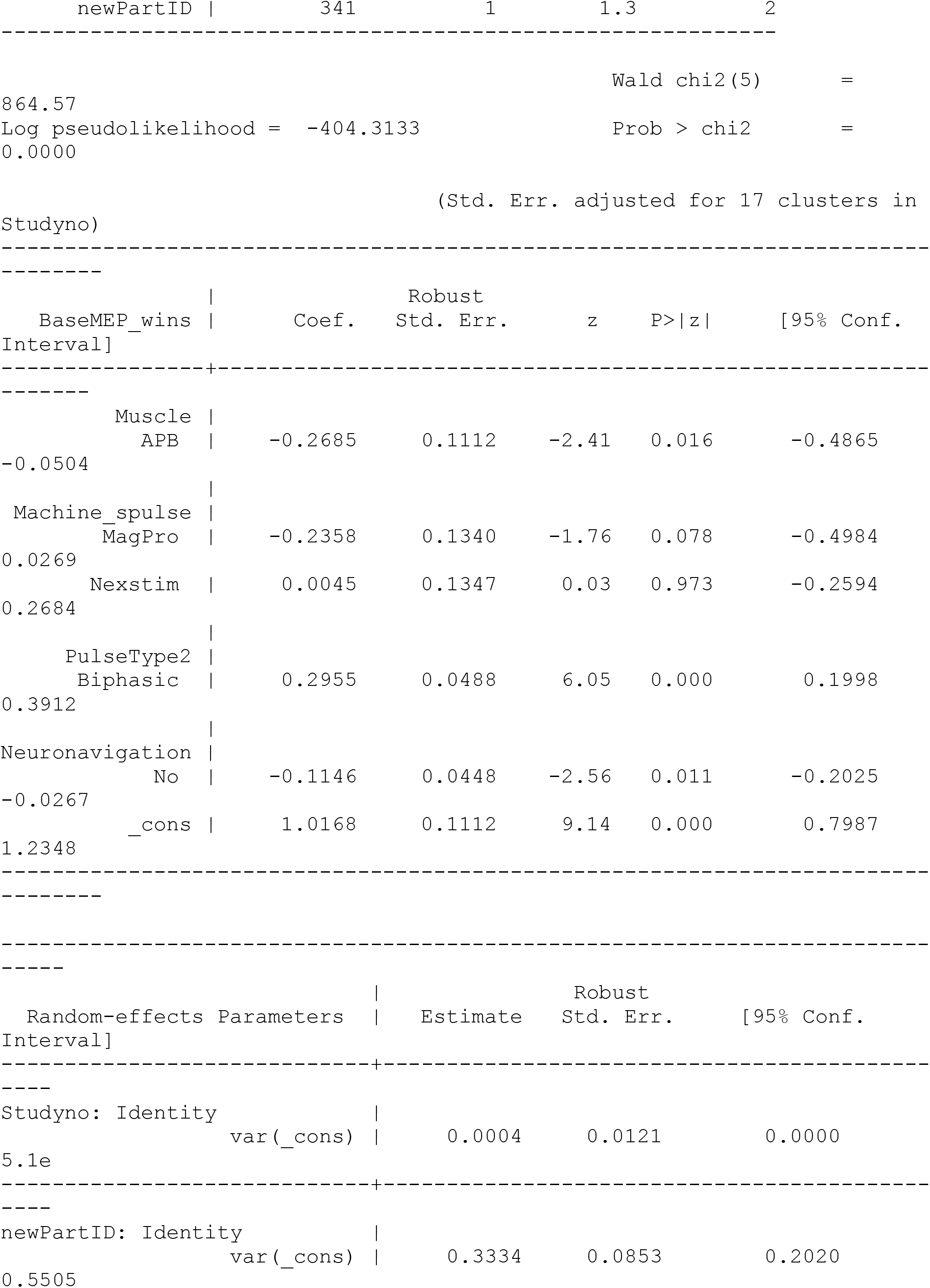
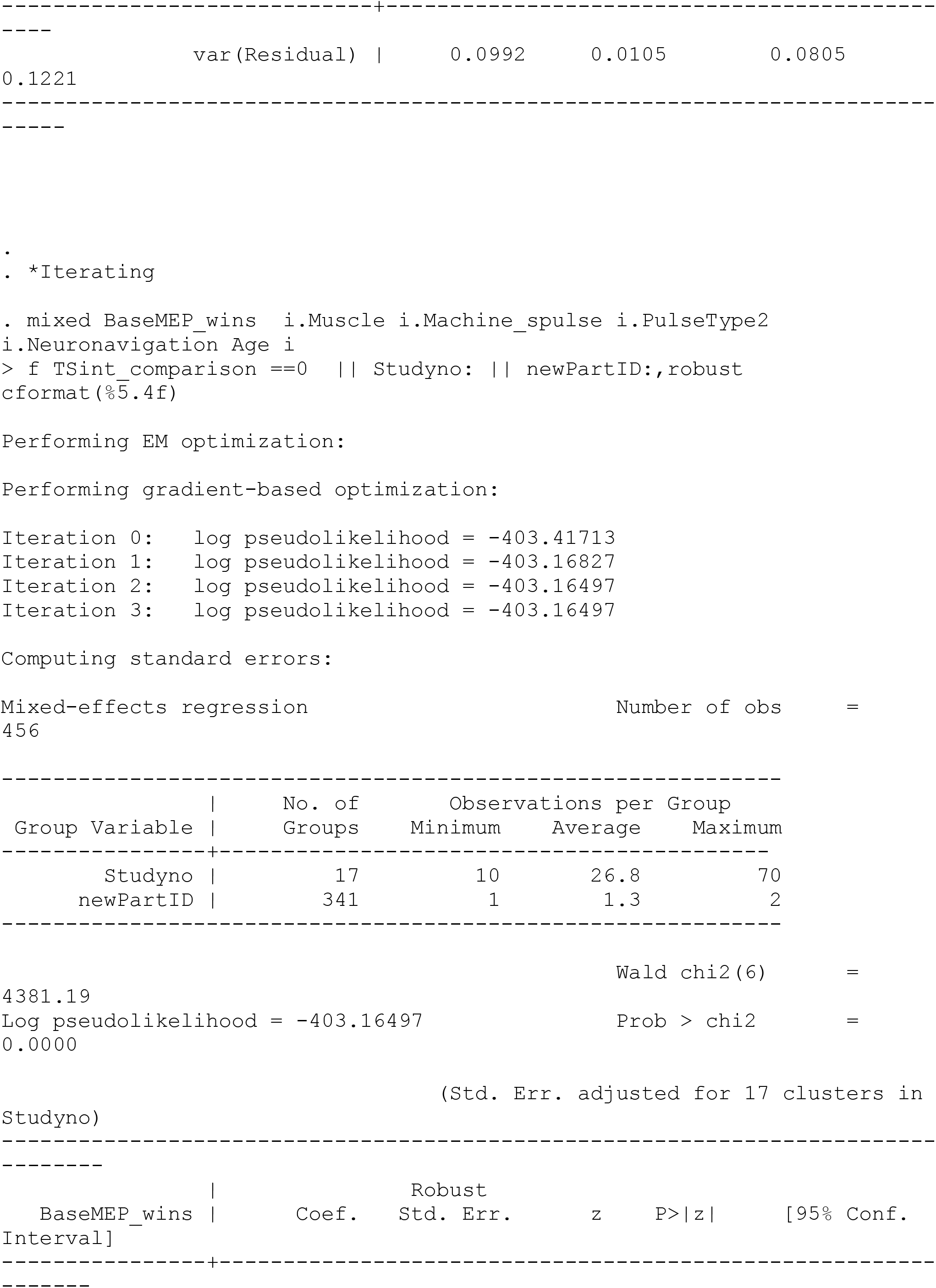
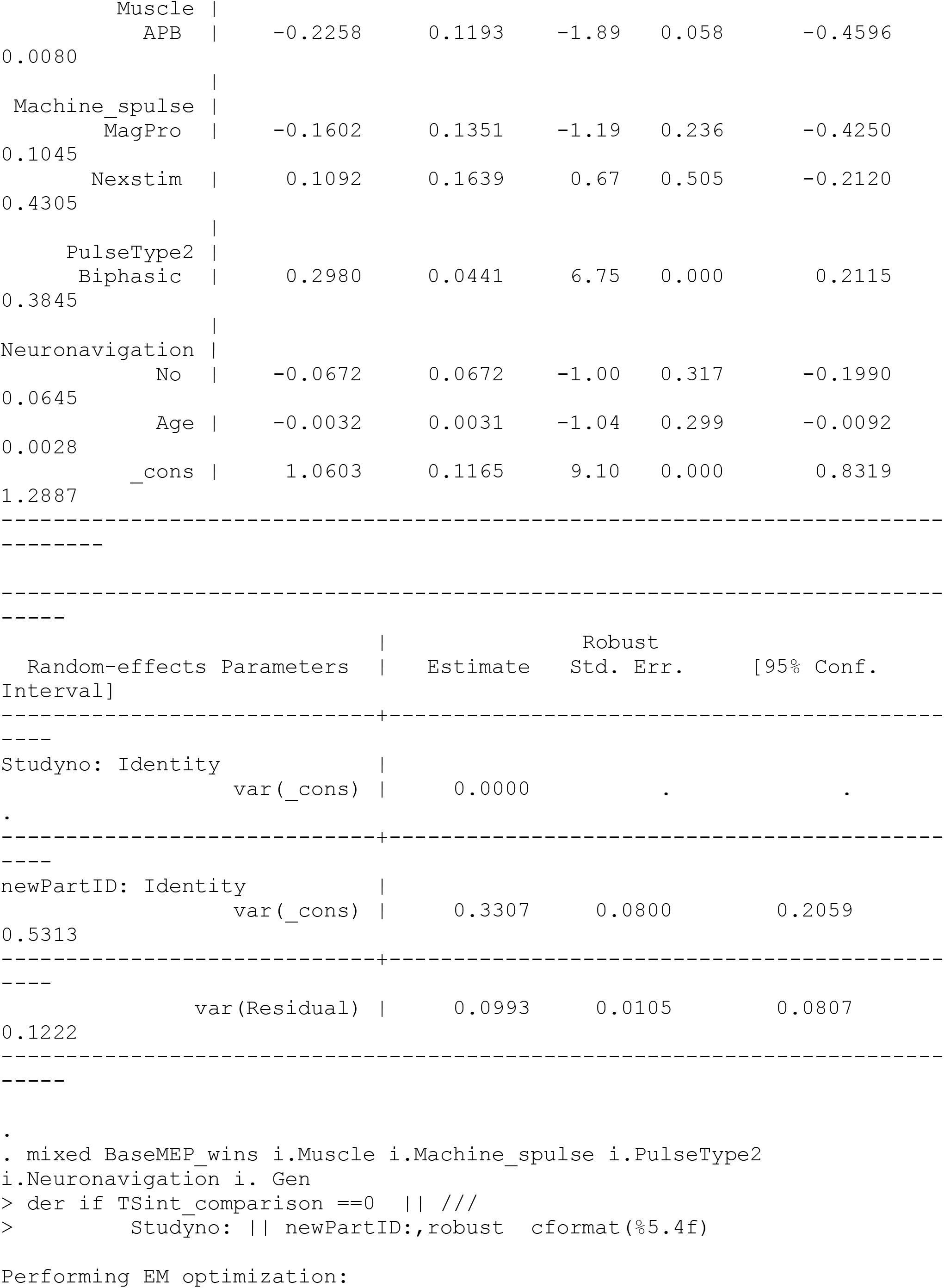
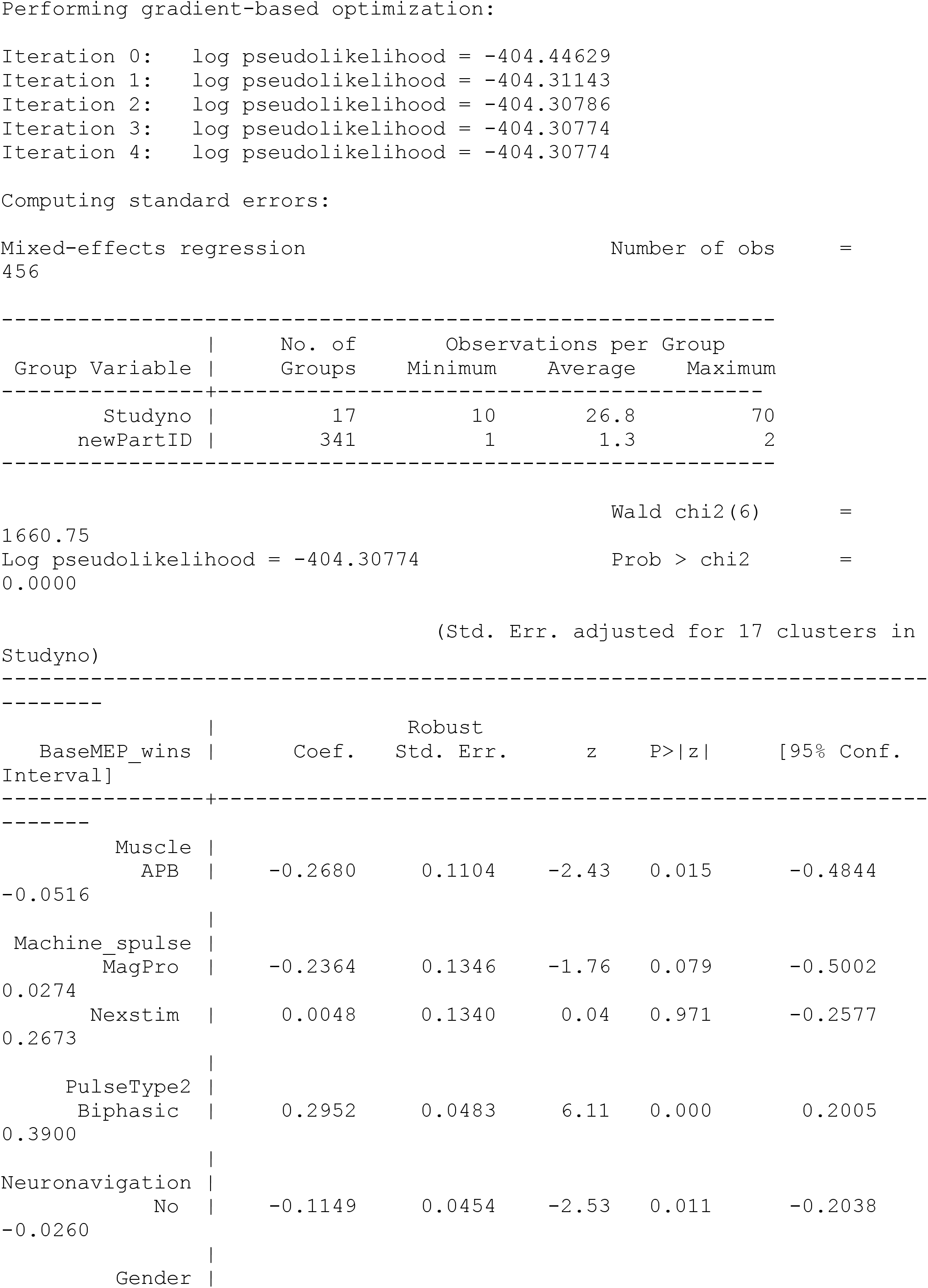
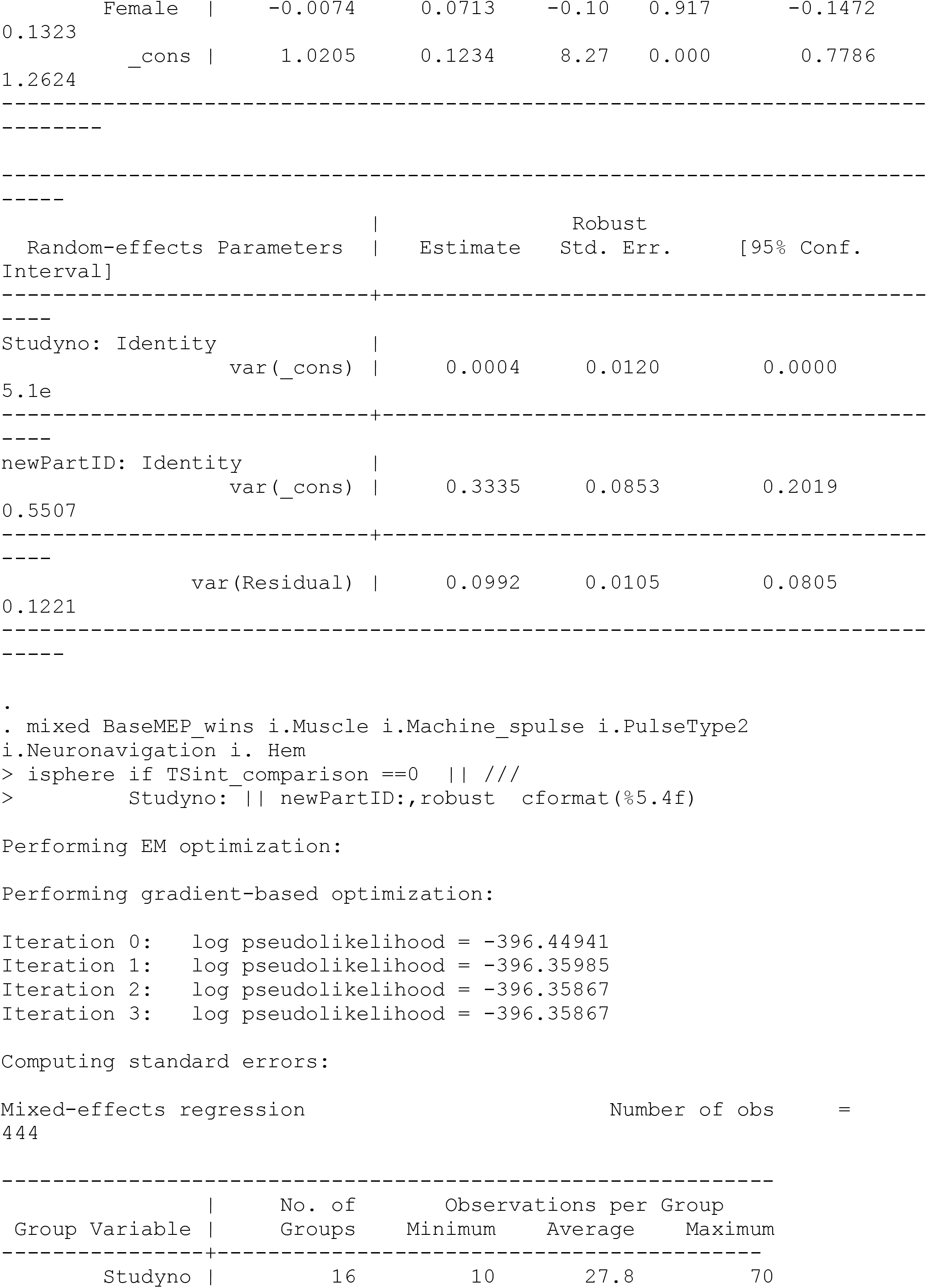
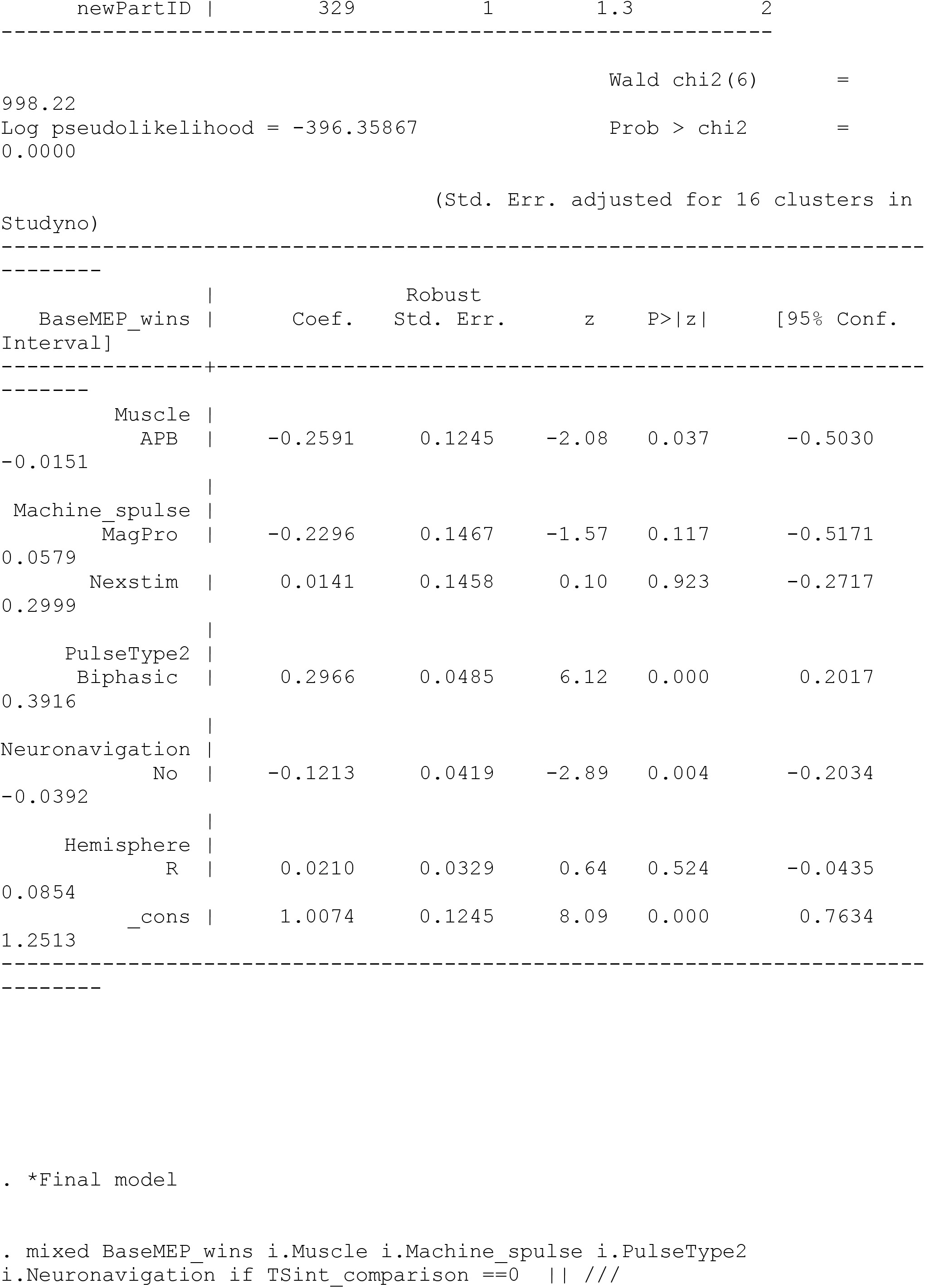
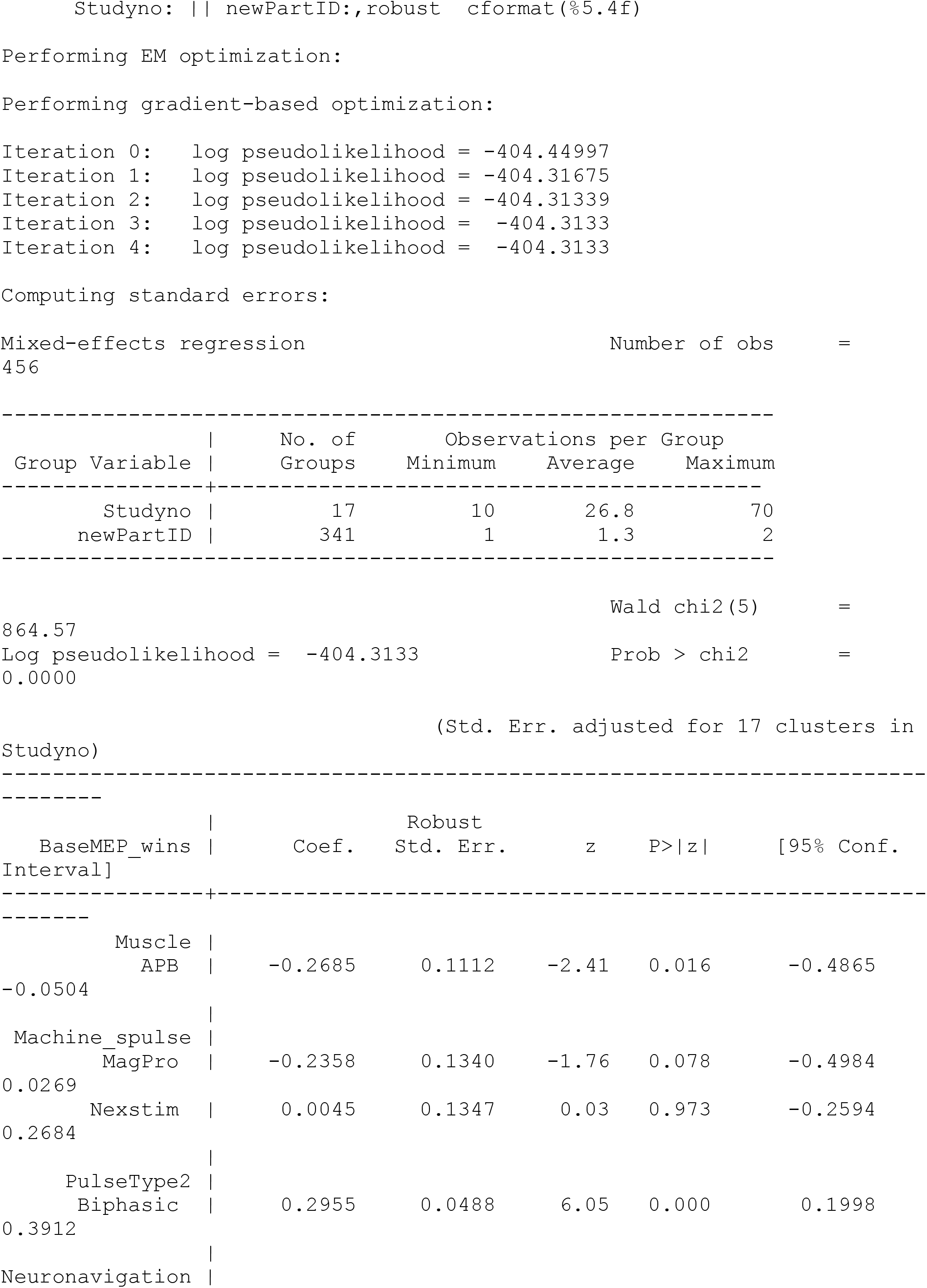
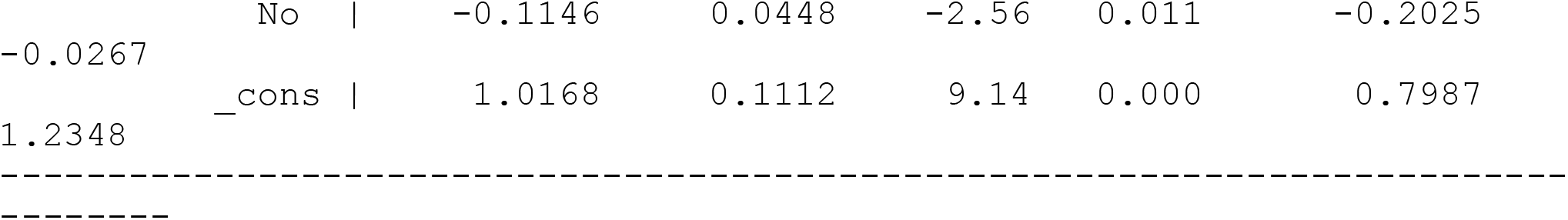

**Supplementary file 7. Non-linear relationships for 120% RMT single pulse MEP amplitude.** Post-hoc analyses demonstrated significant non-linear relationships between single pulse MEP amplitude and monophasic AMT, biphasic RMT, and biphasic AMT.

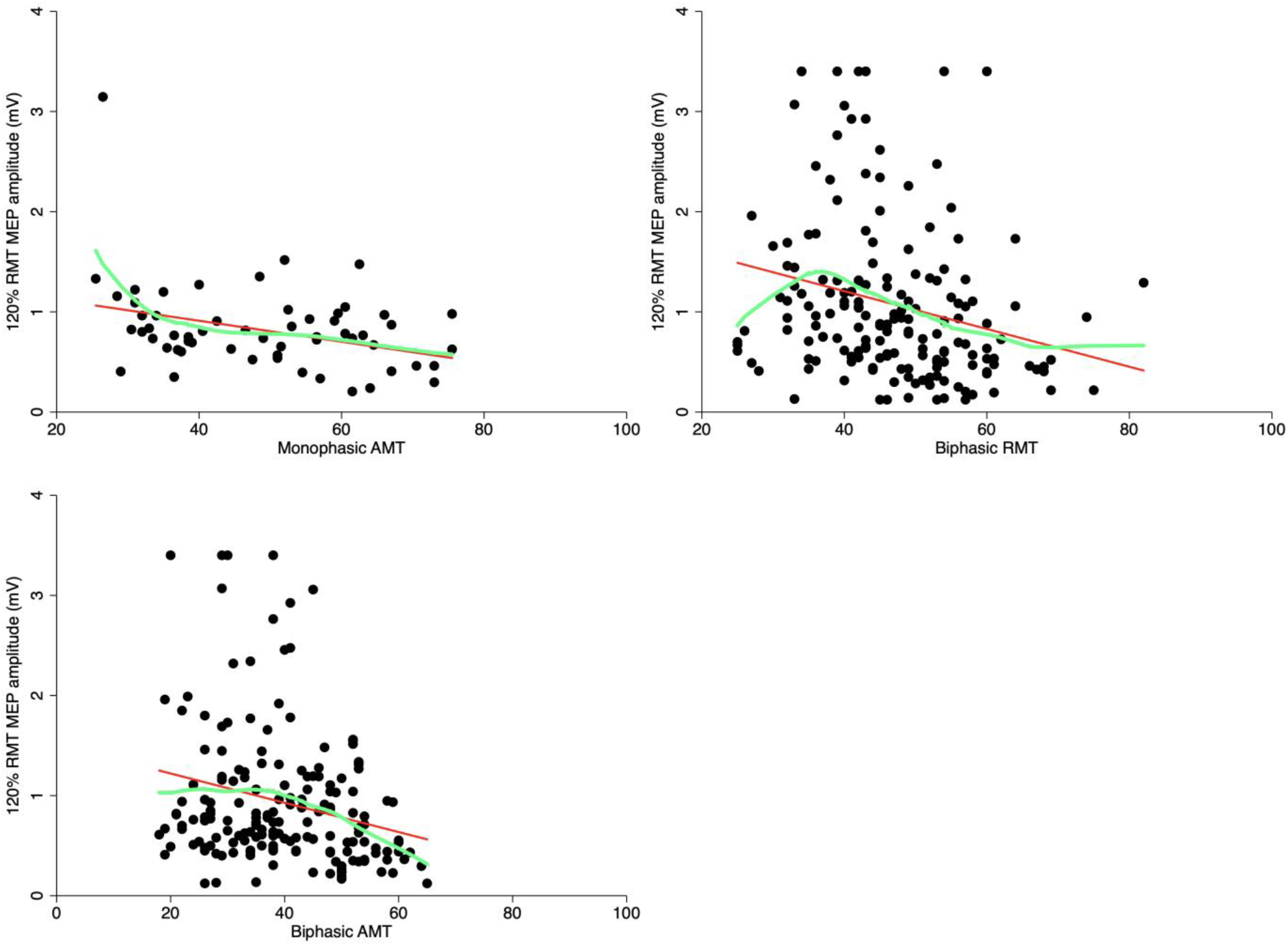

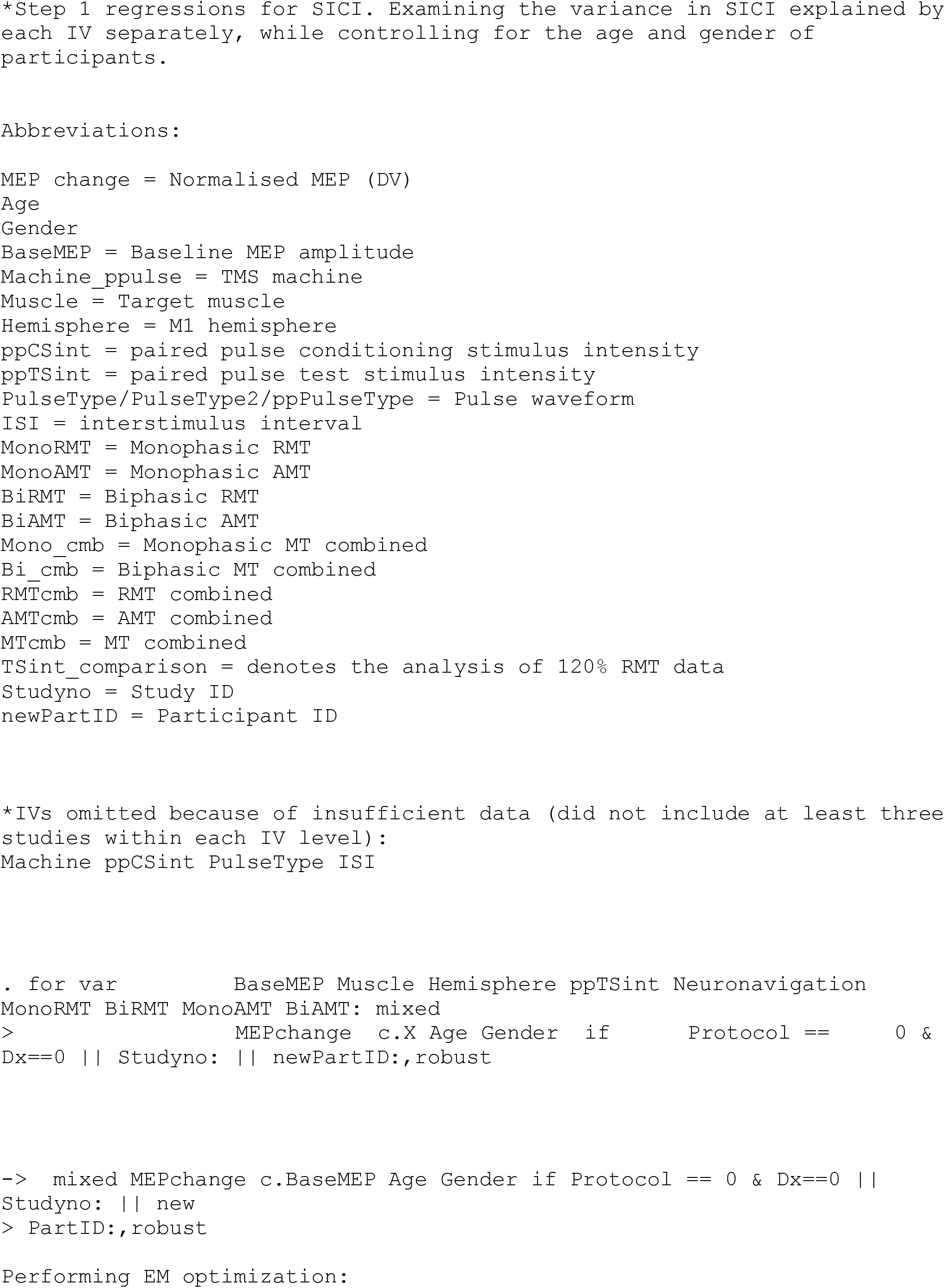
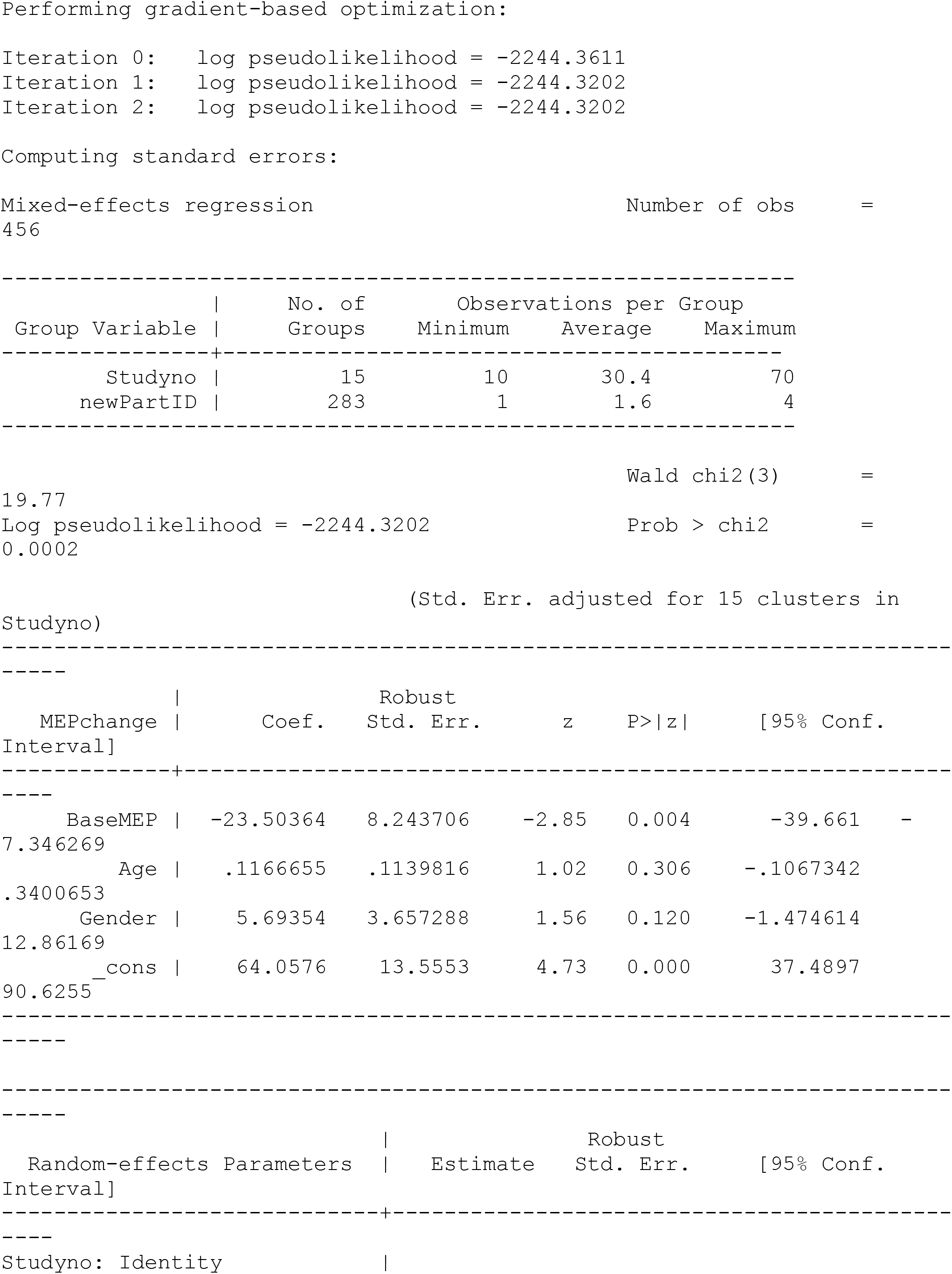
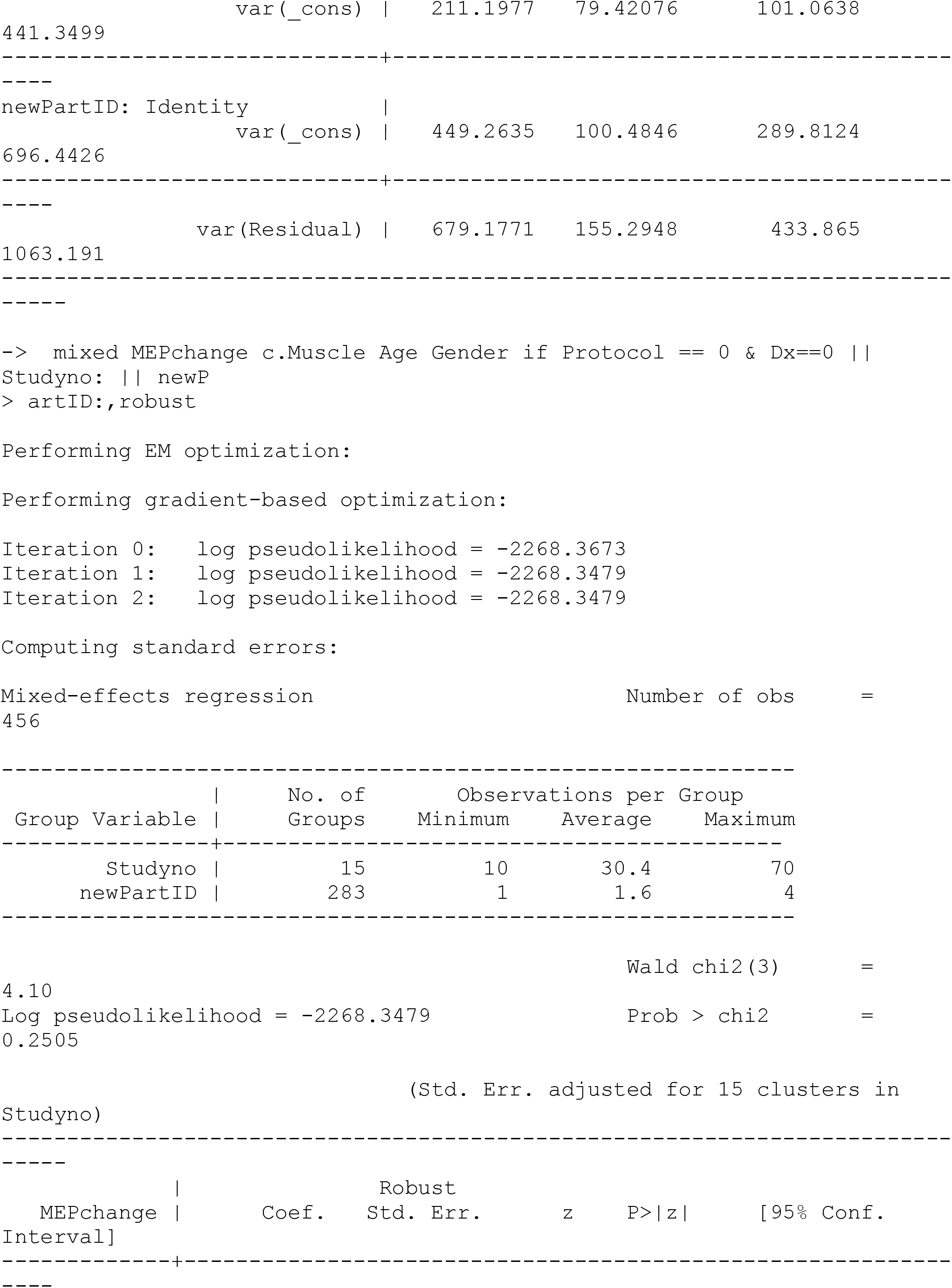
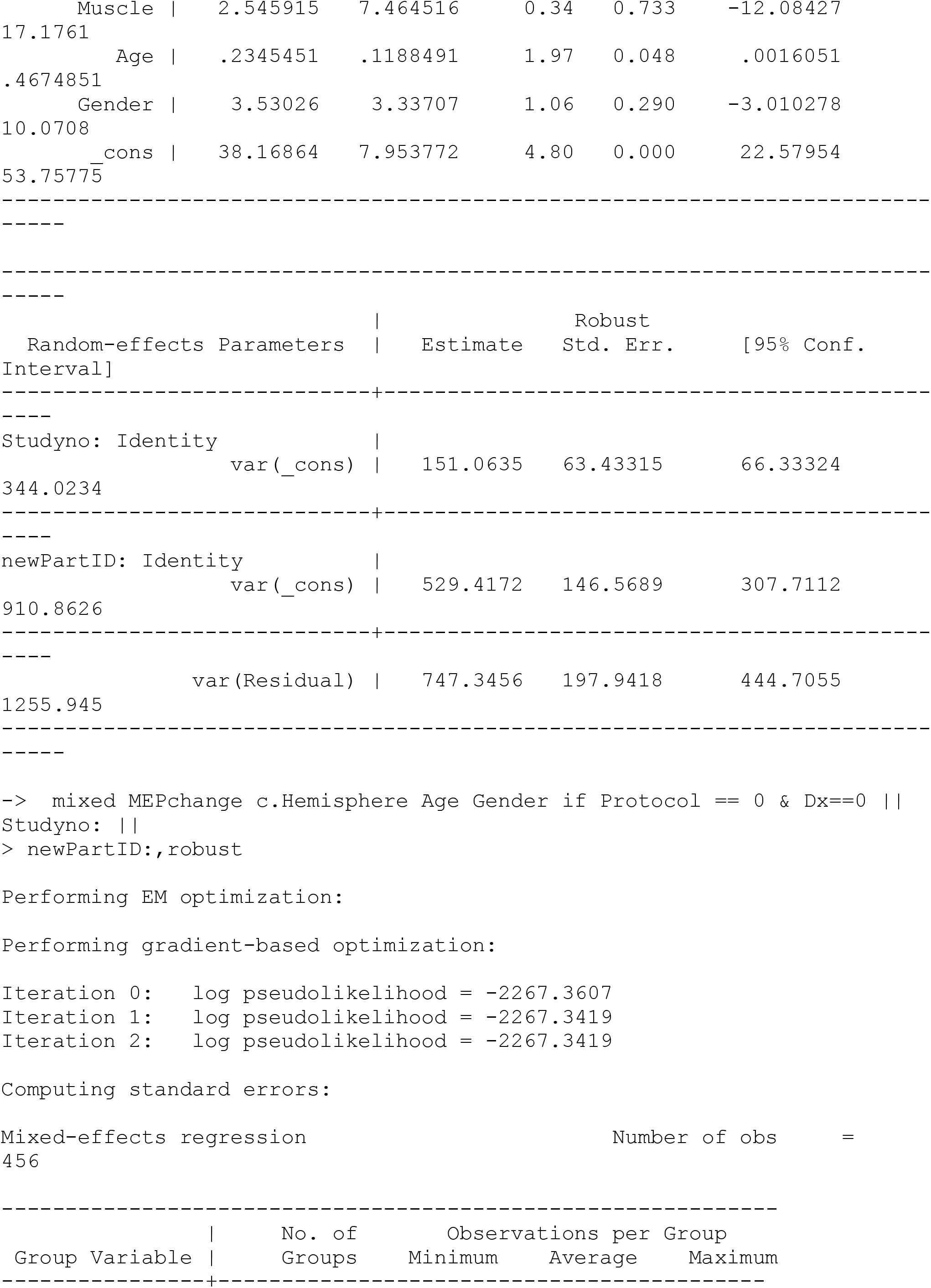
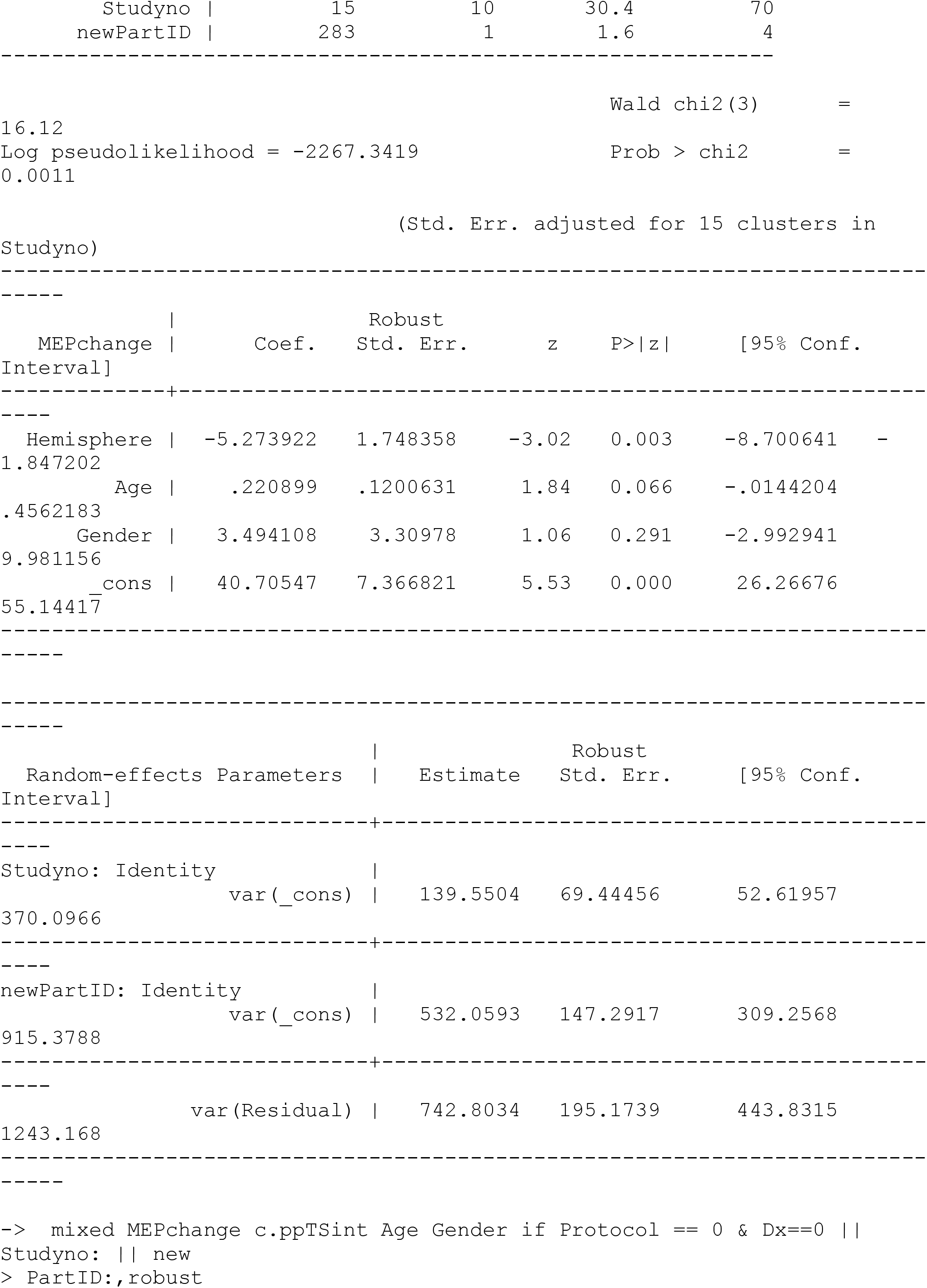
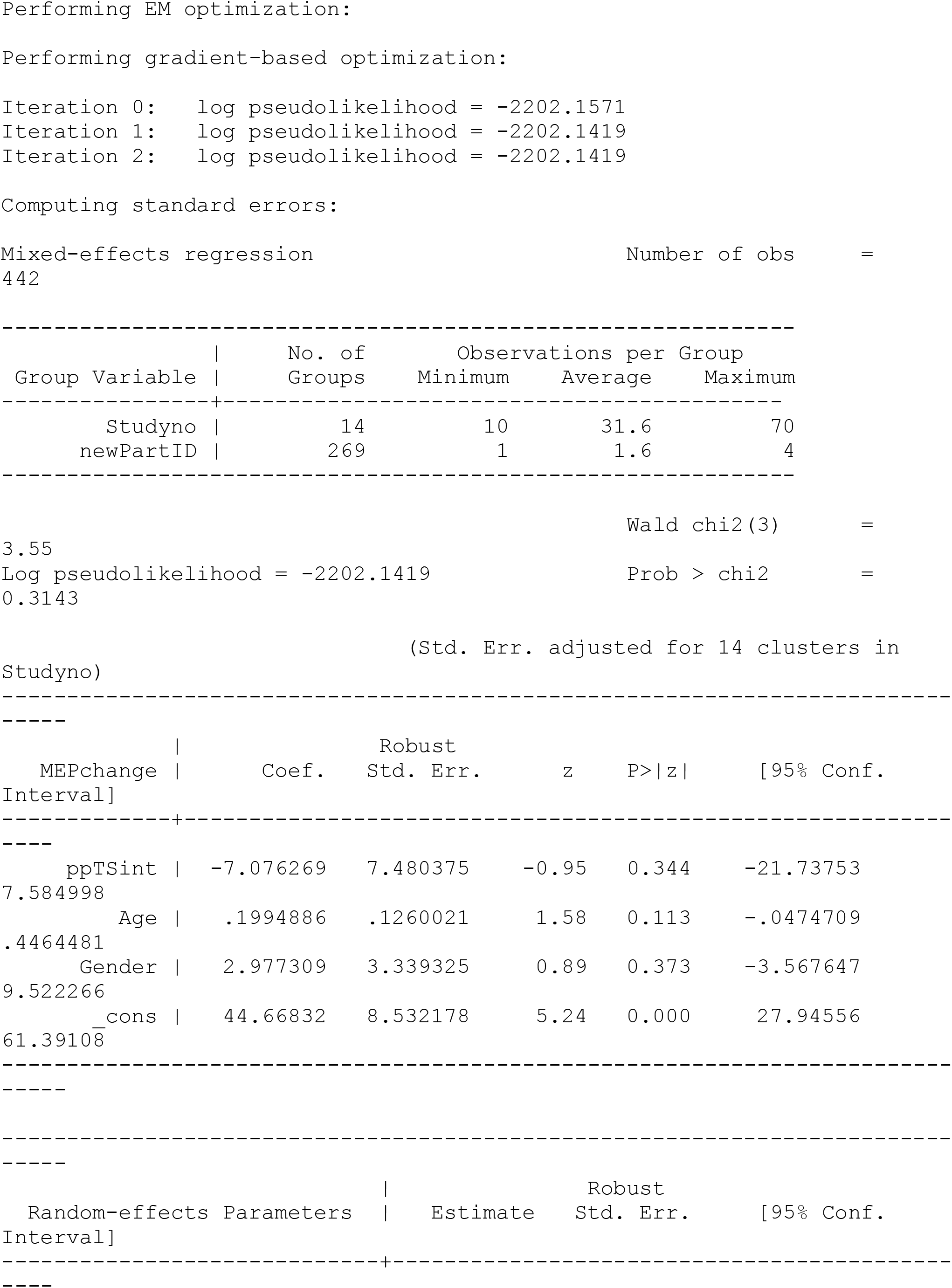
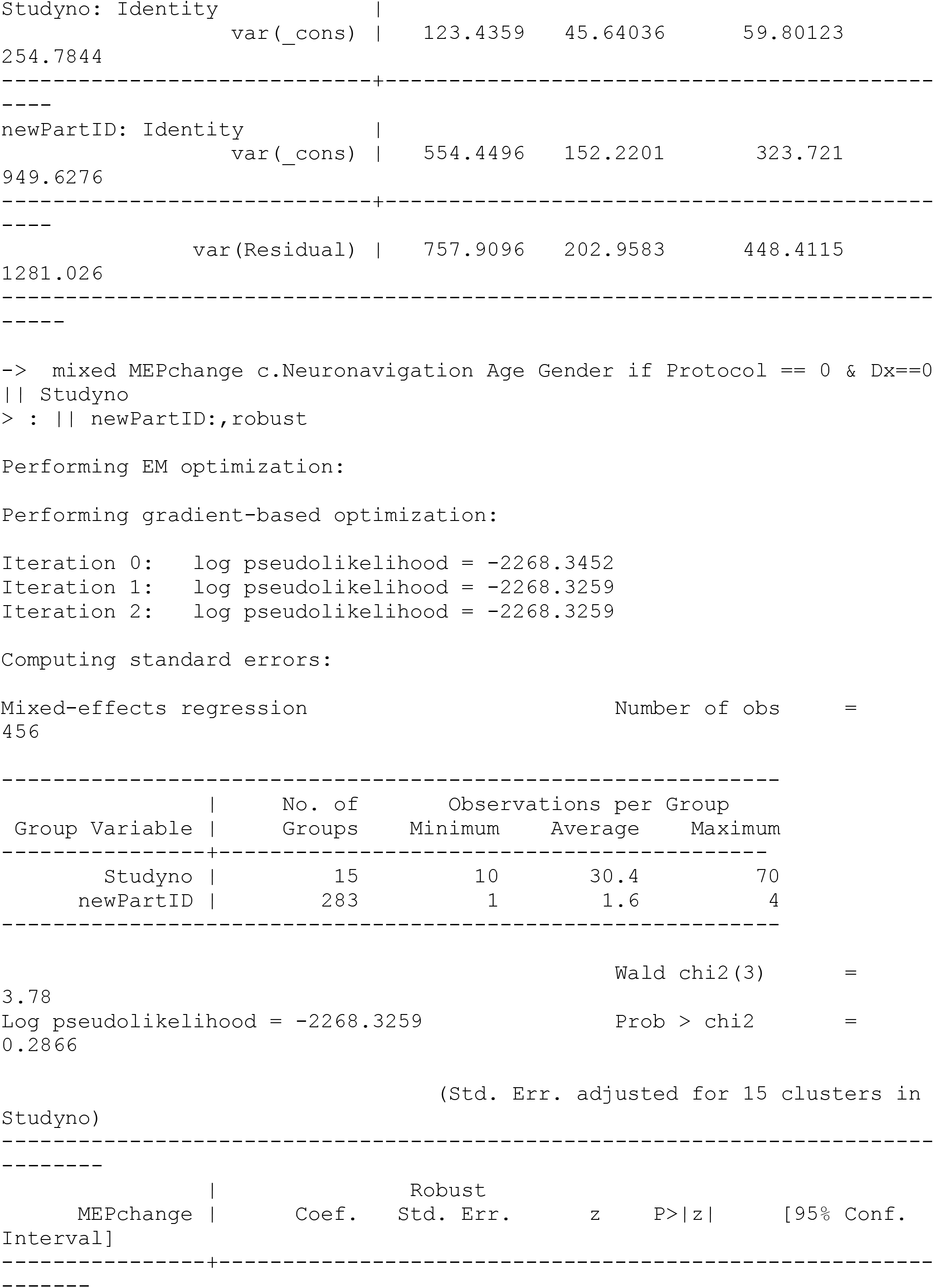
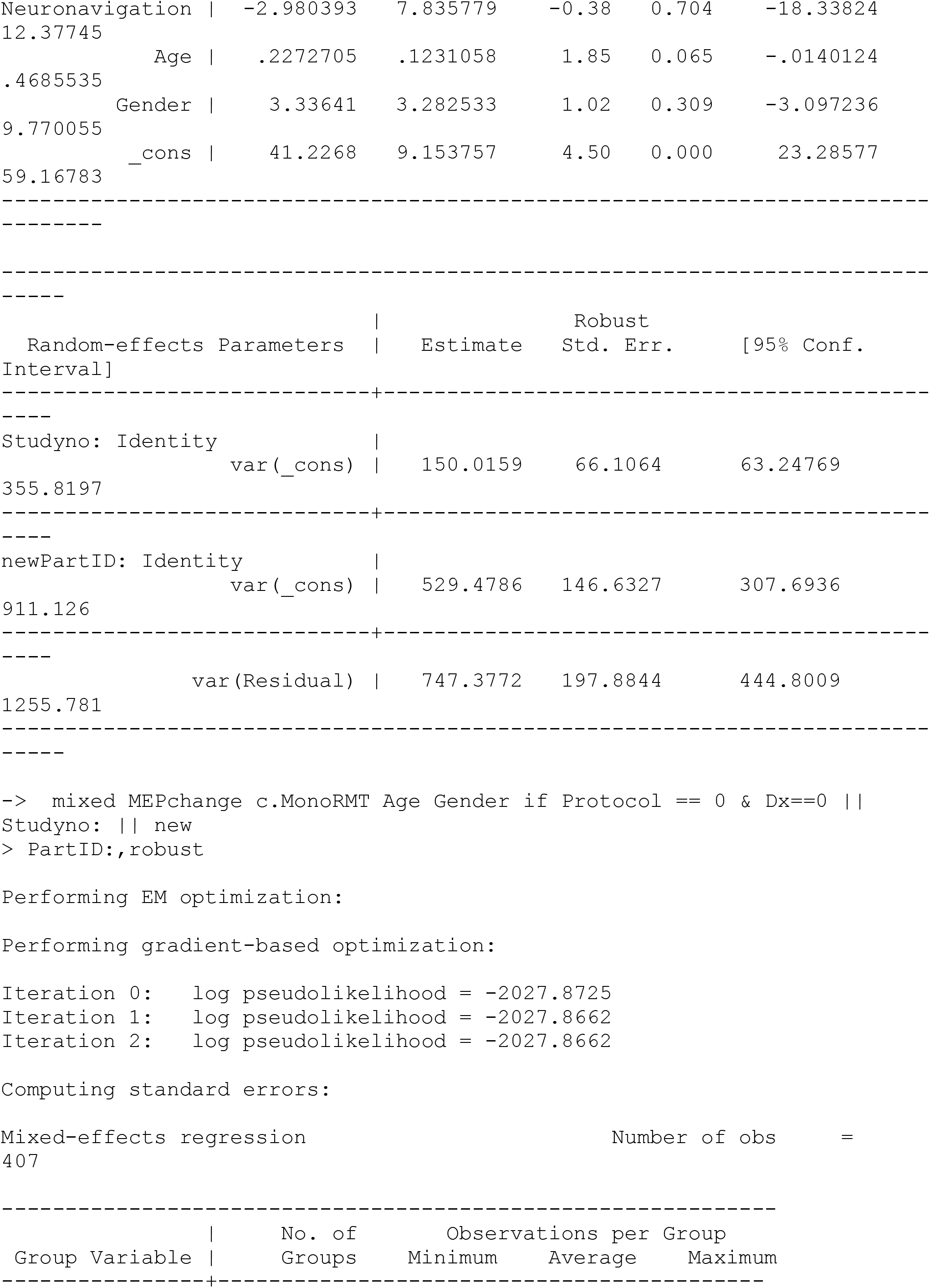
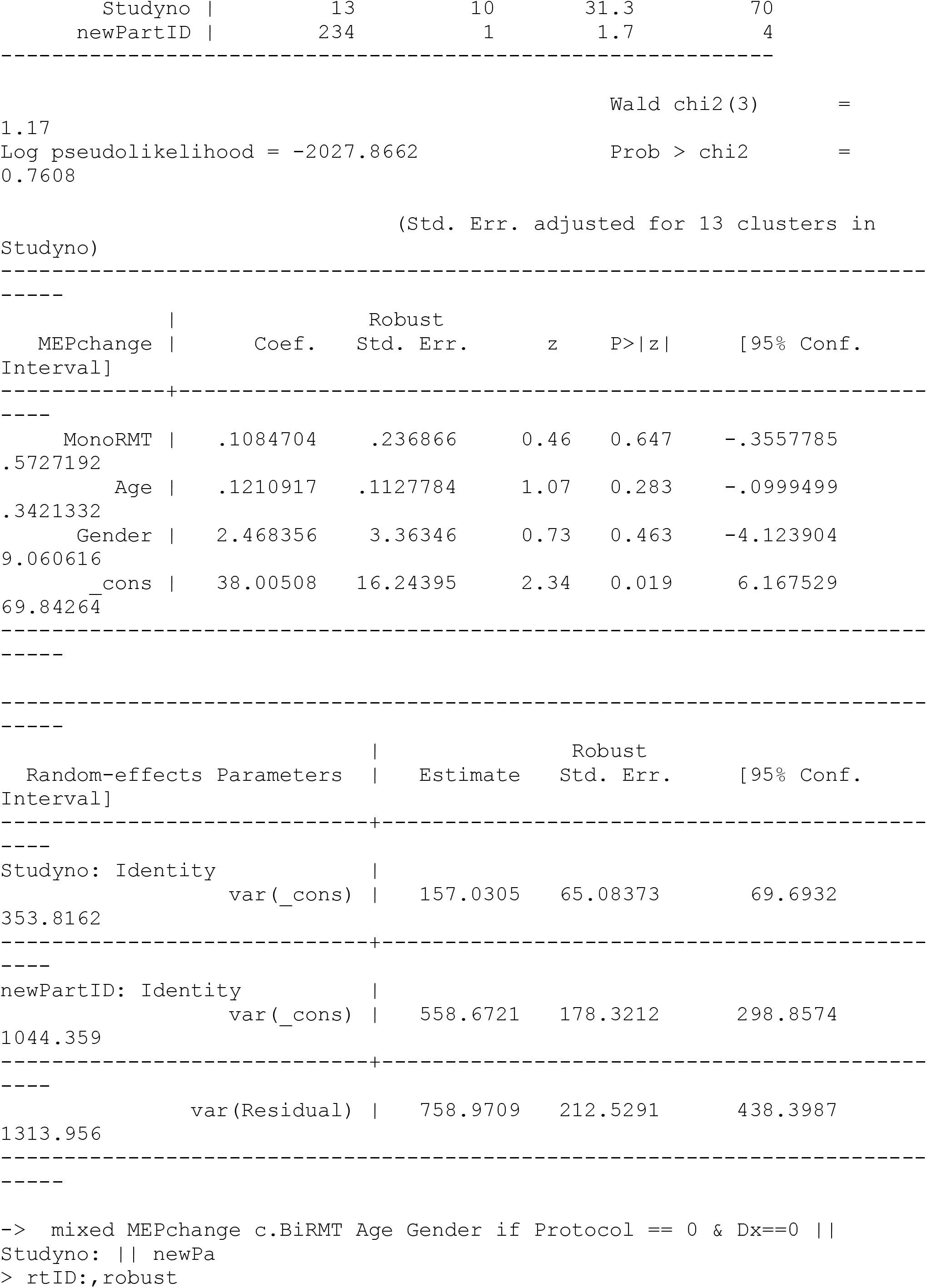
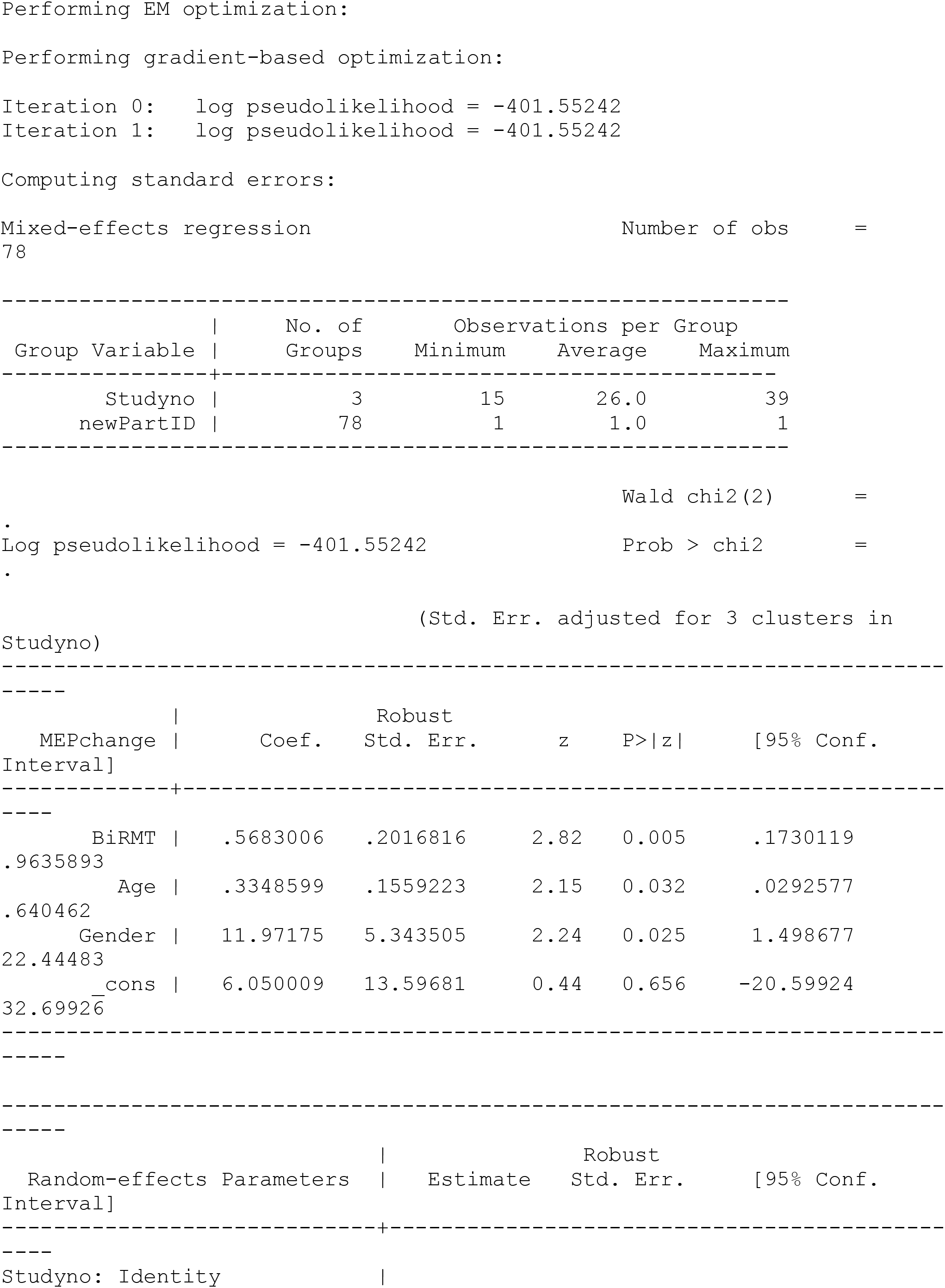
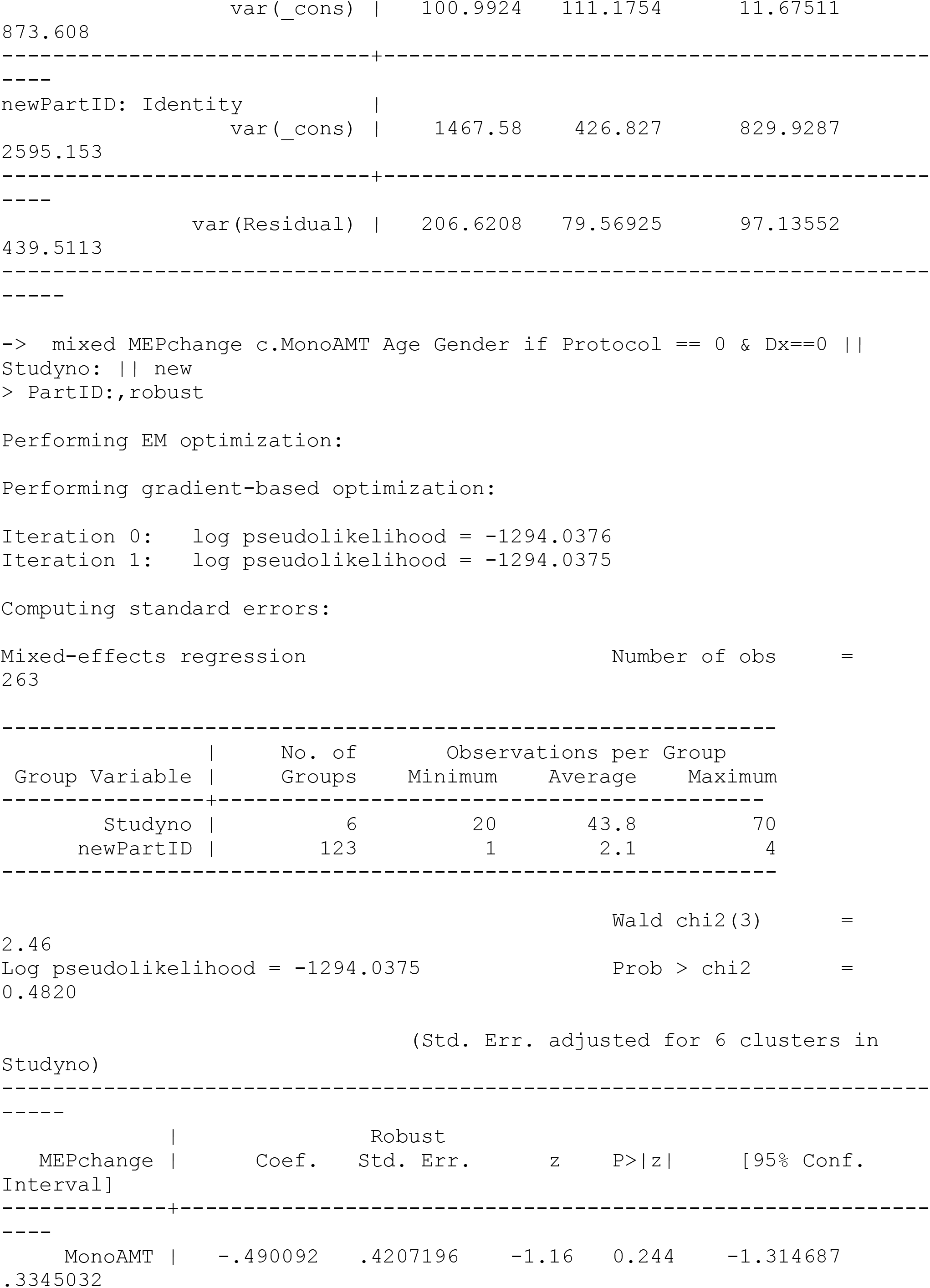
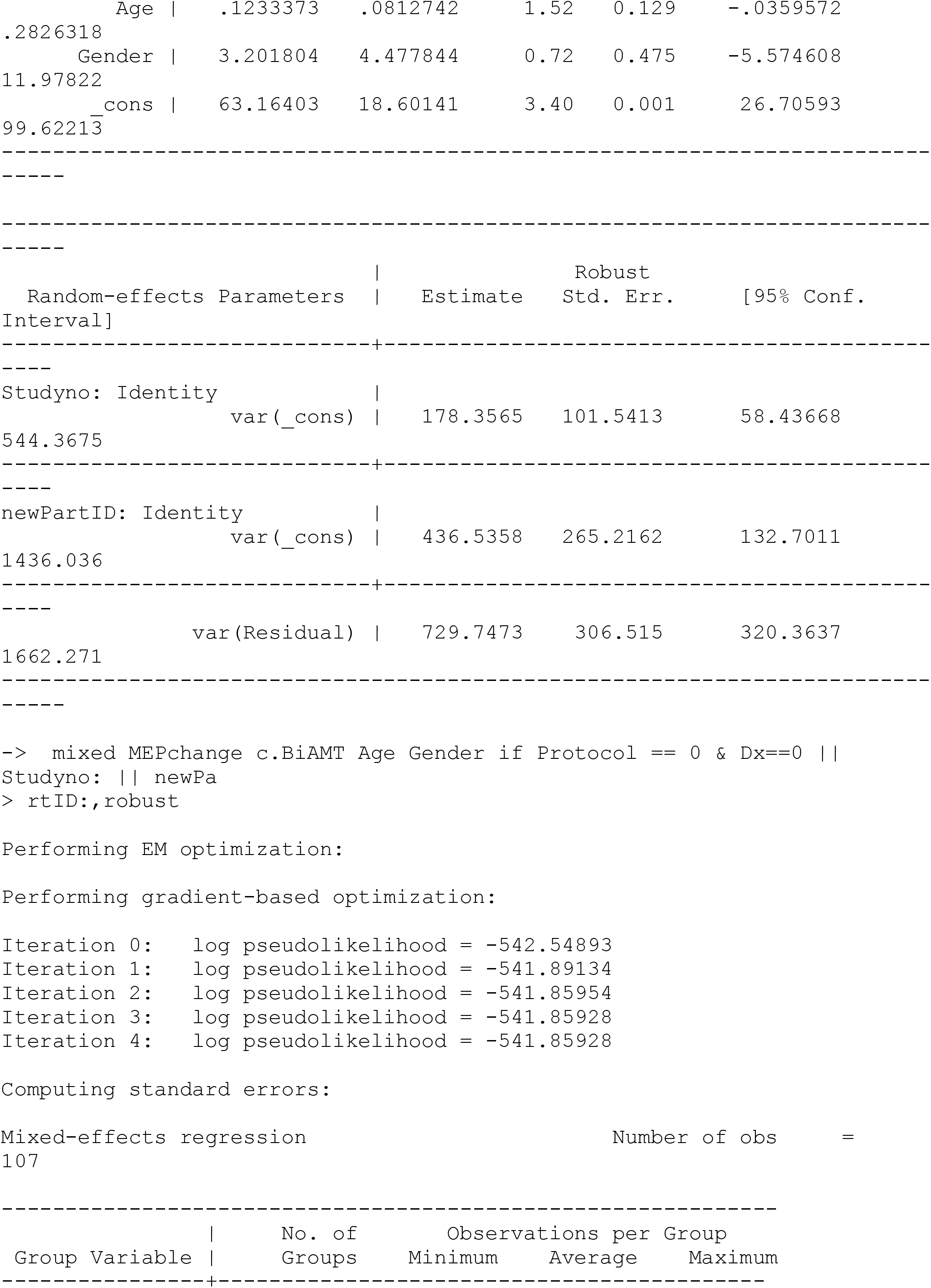
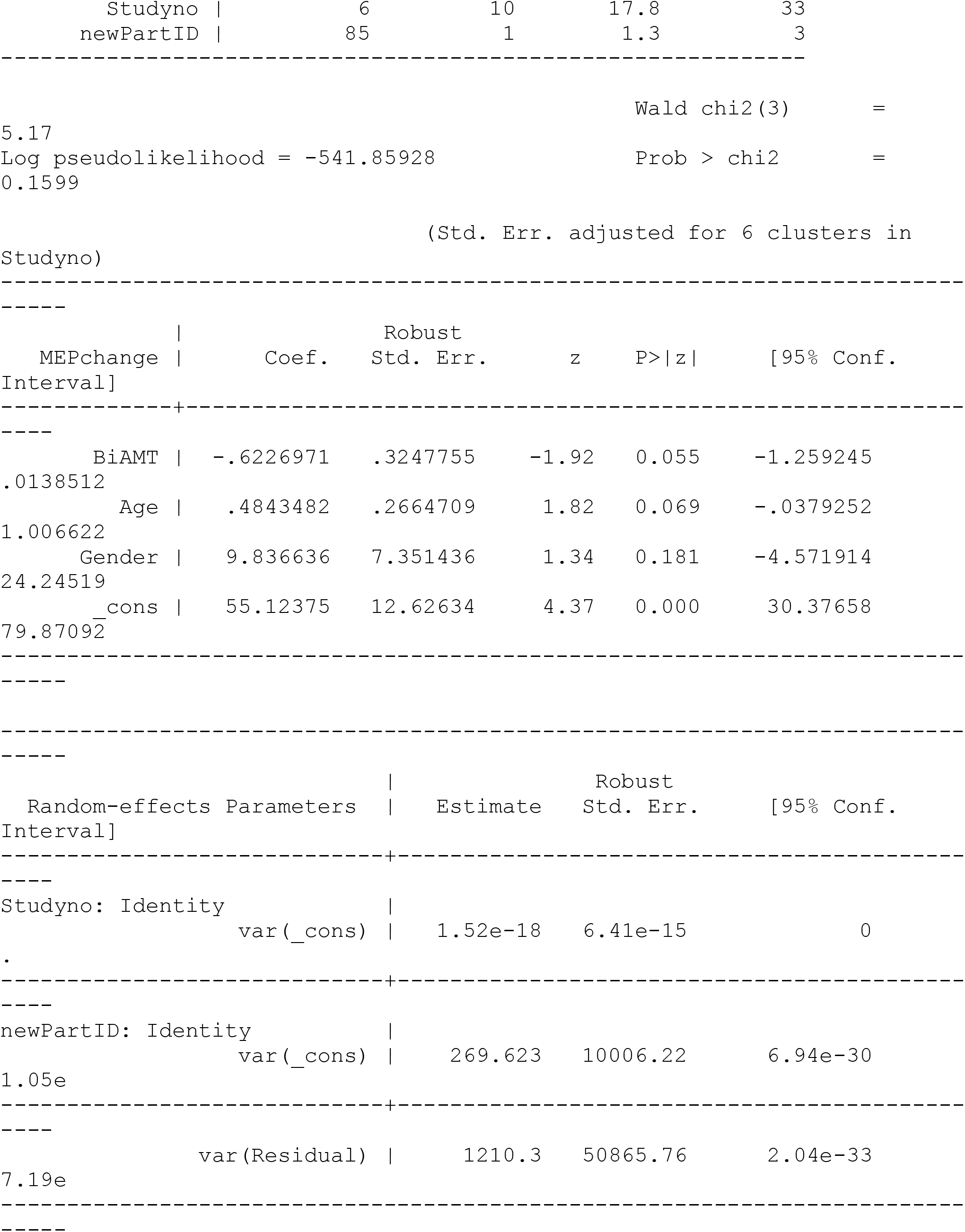
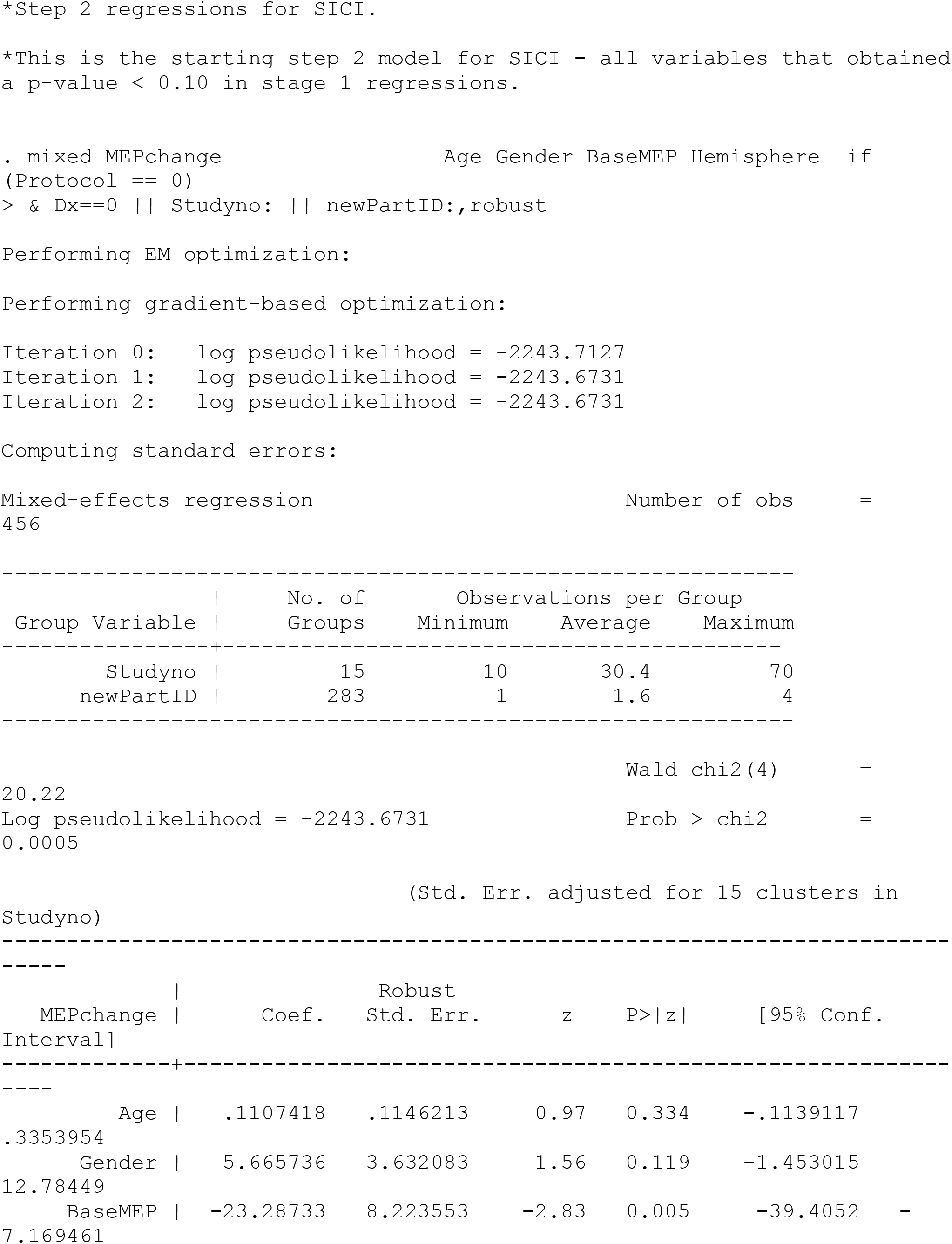
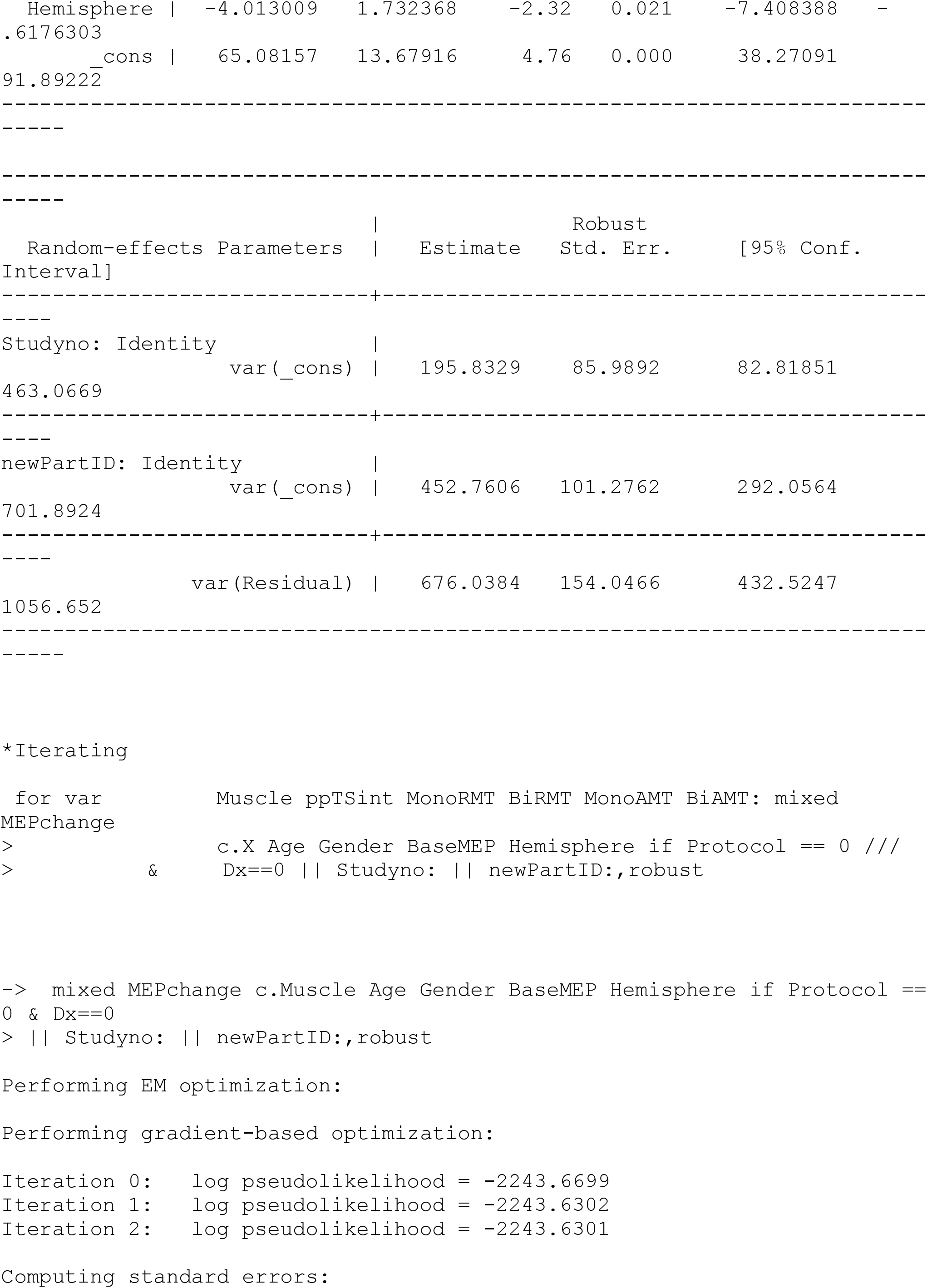
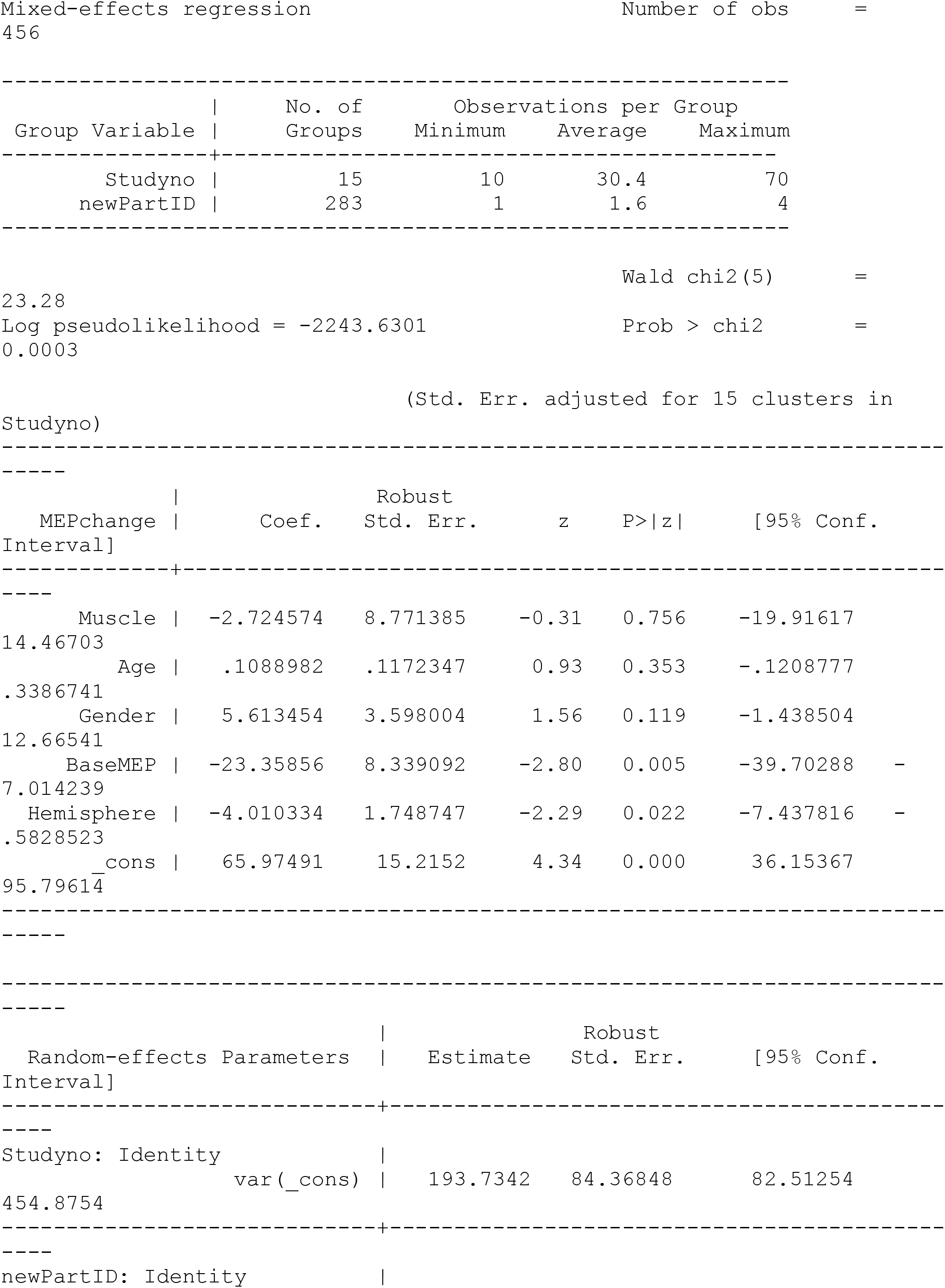
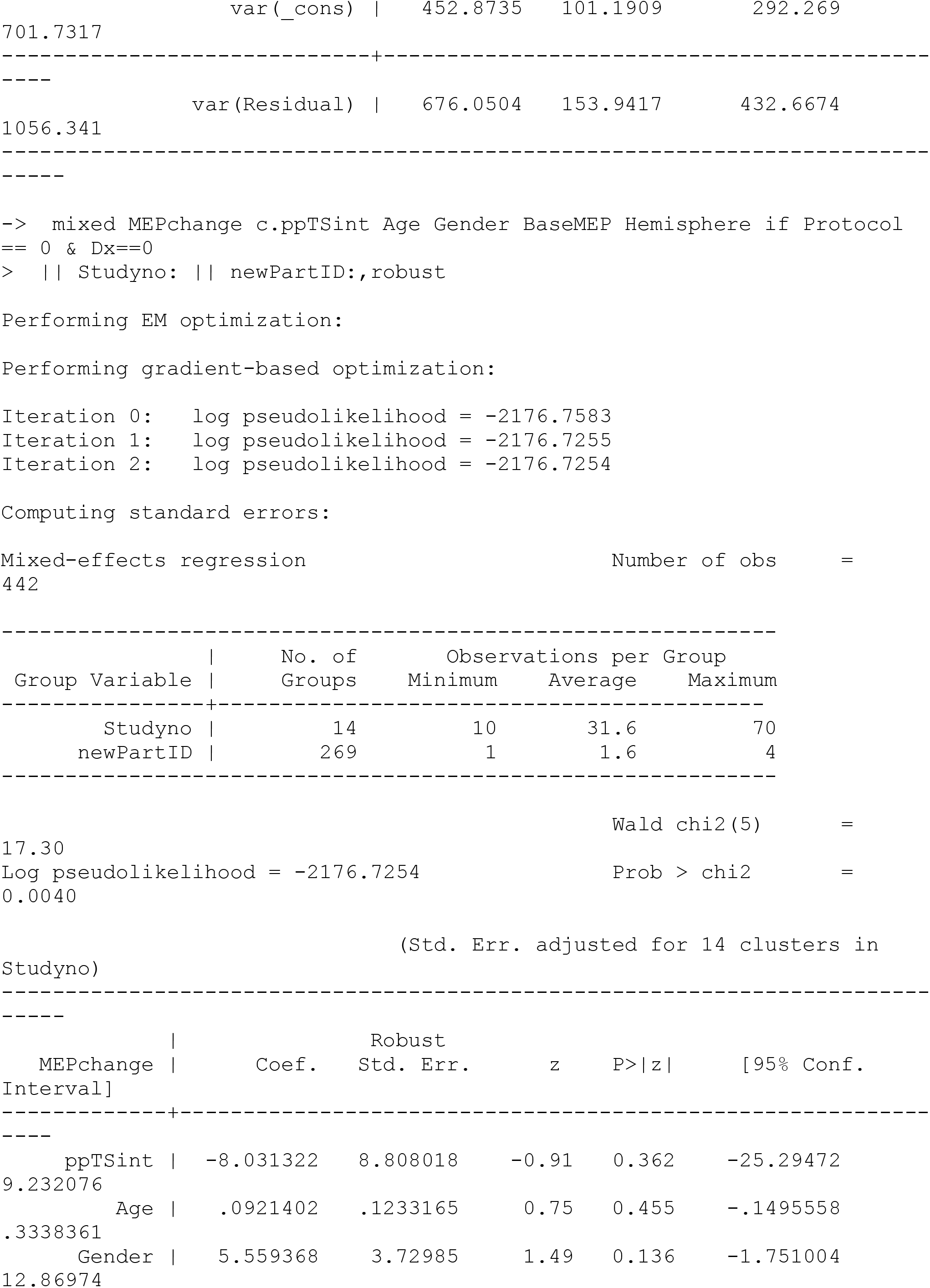
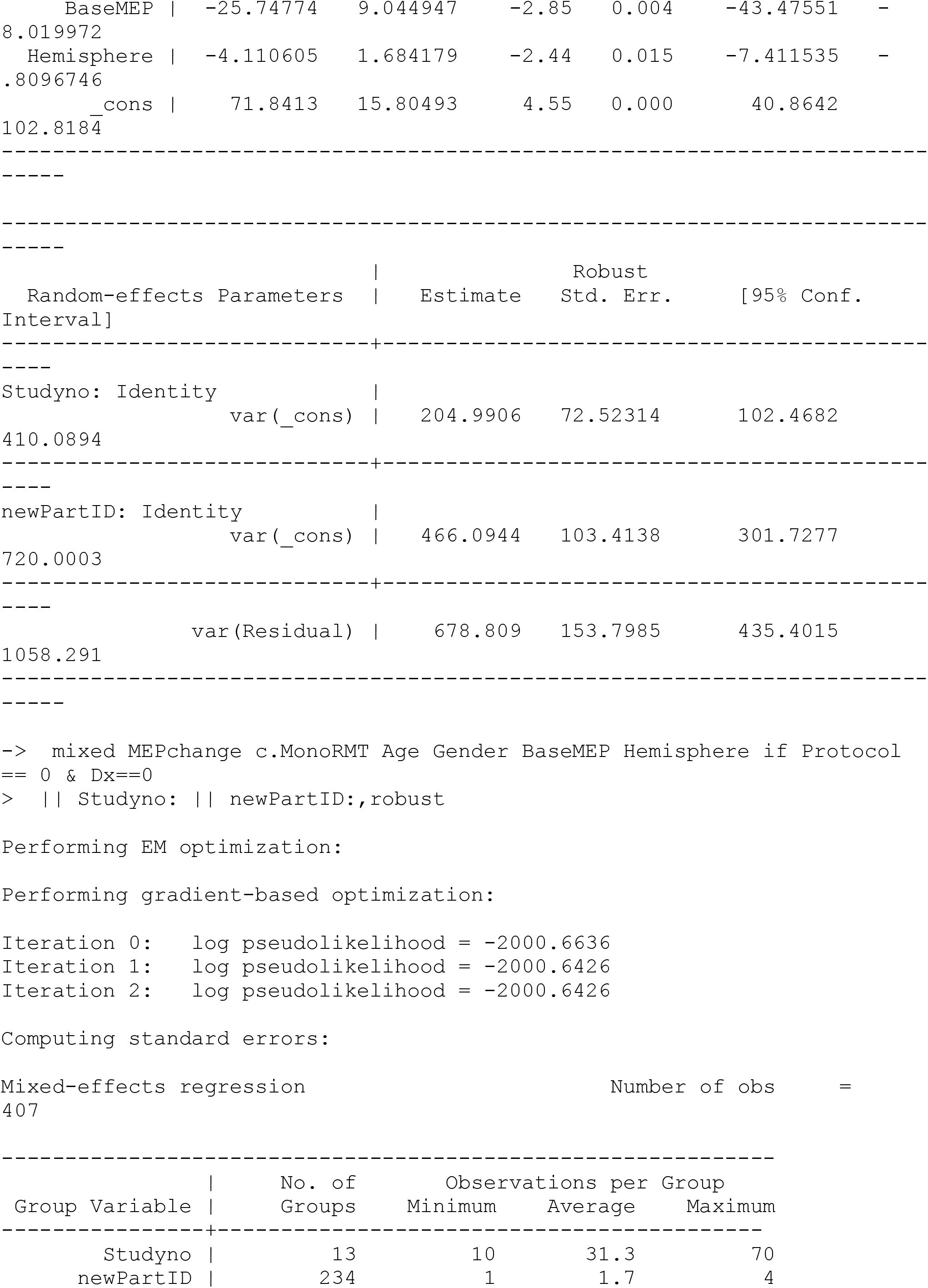
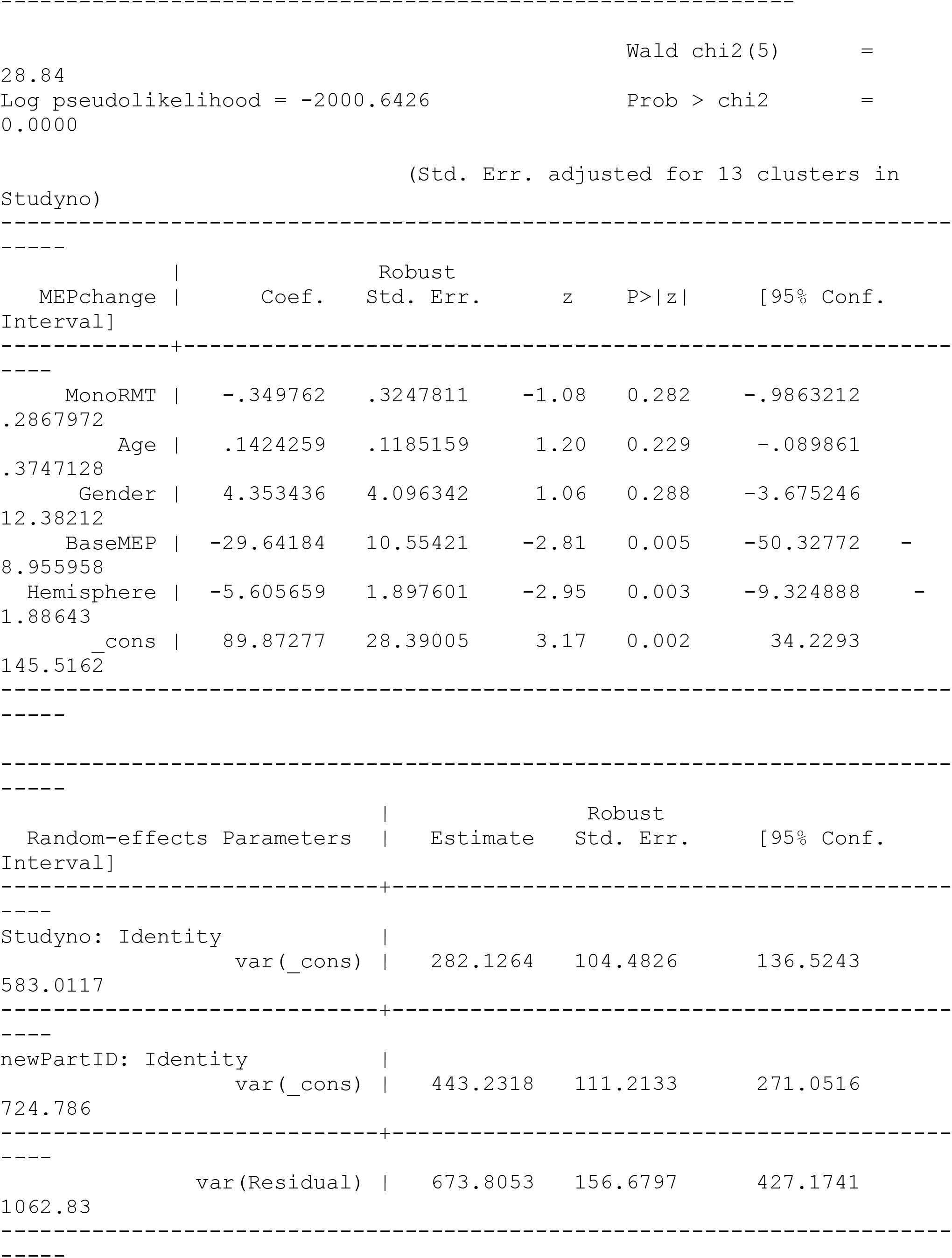
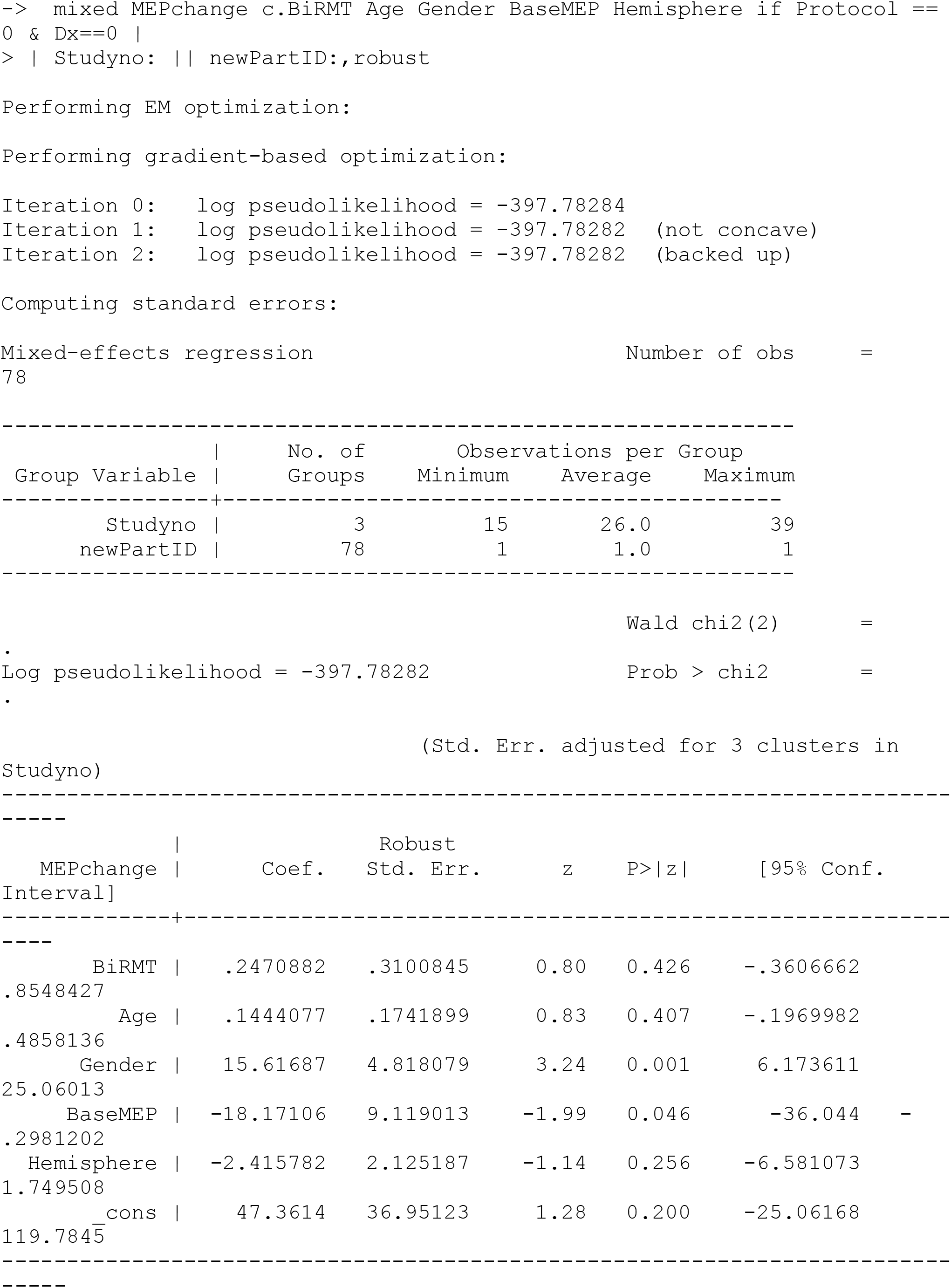
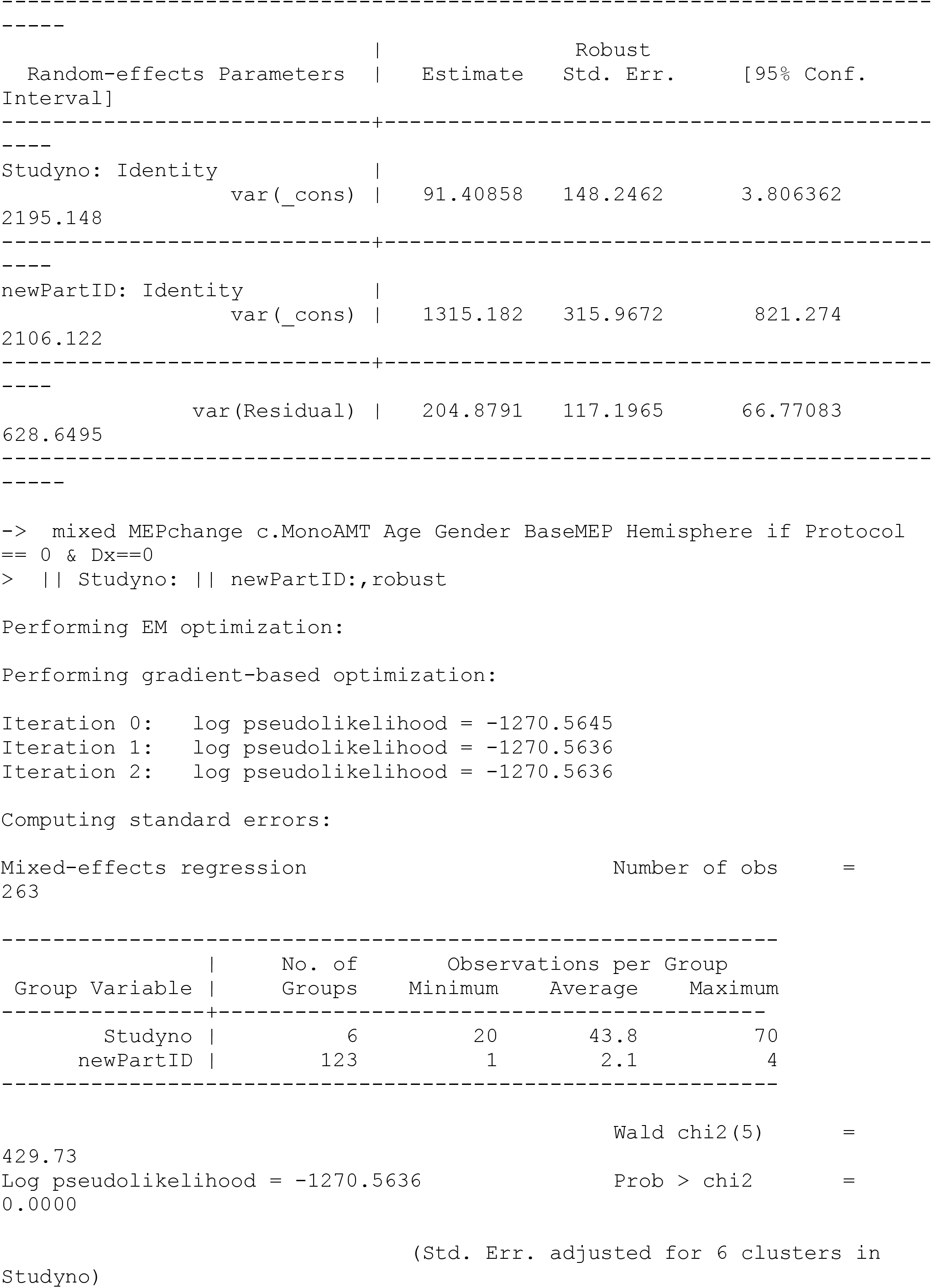
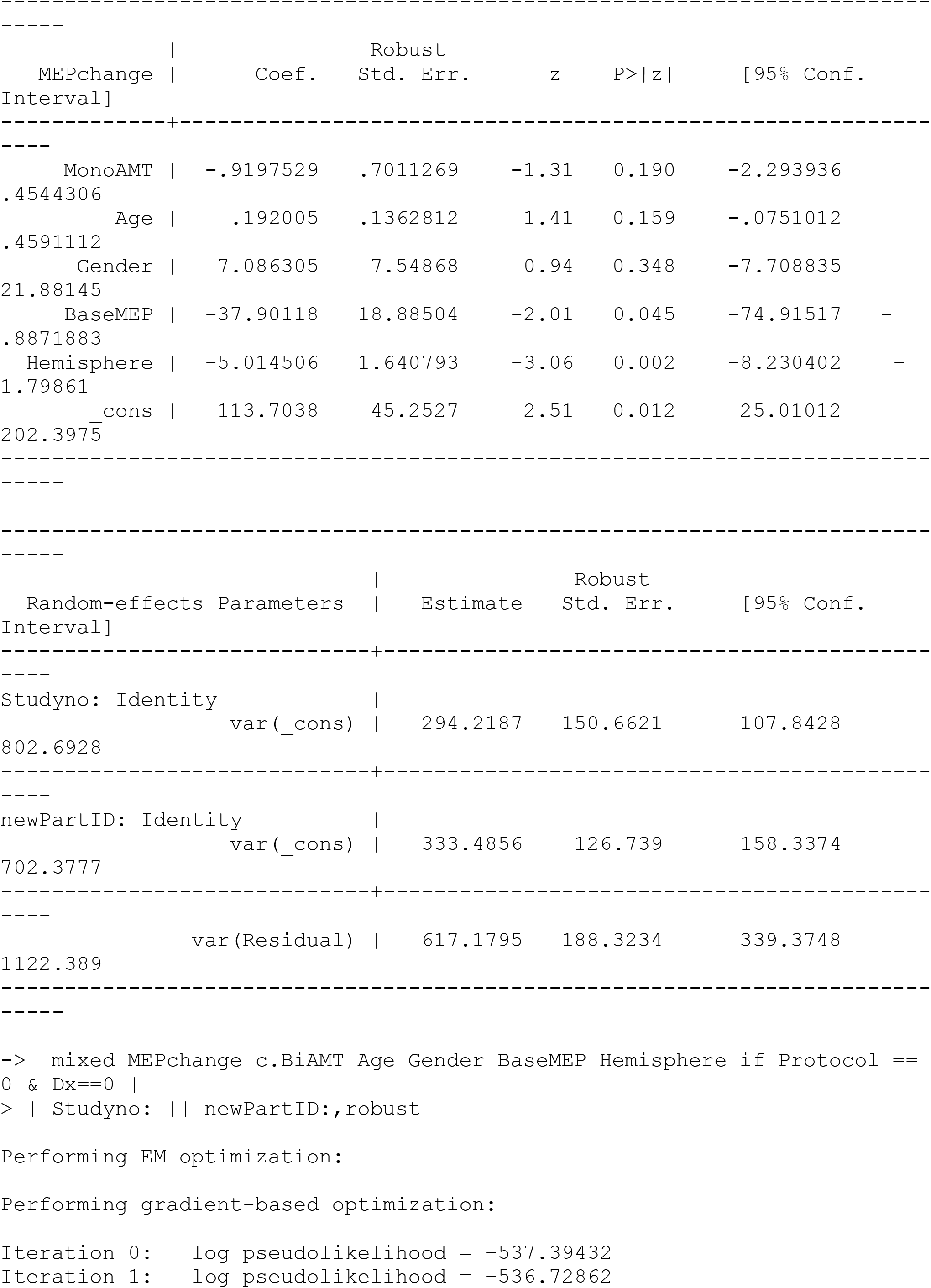
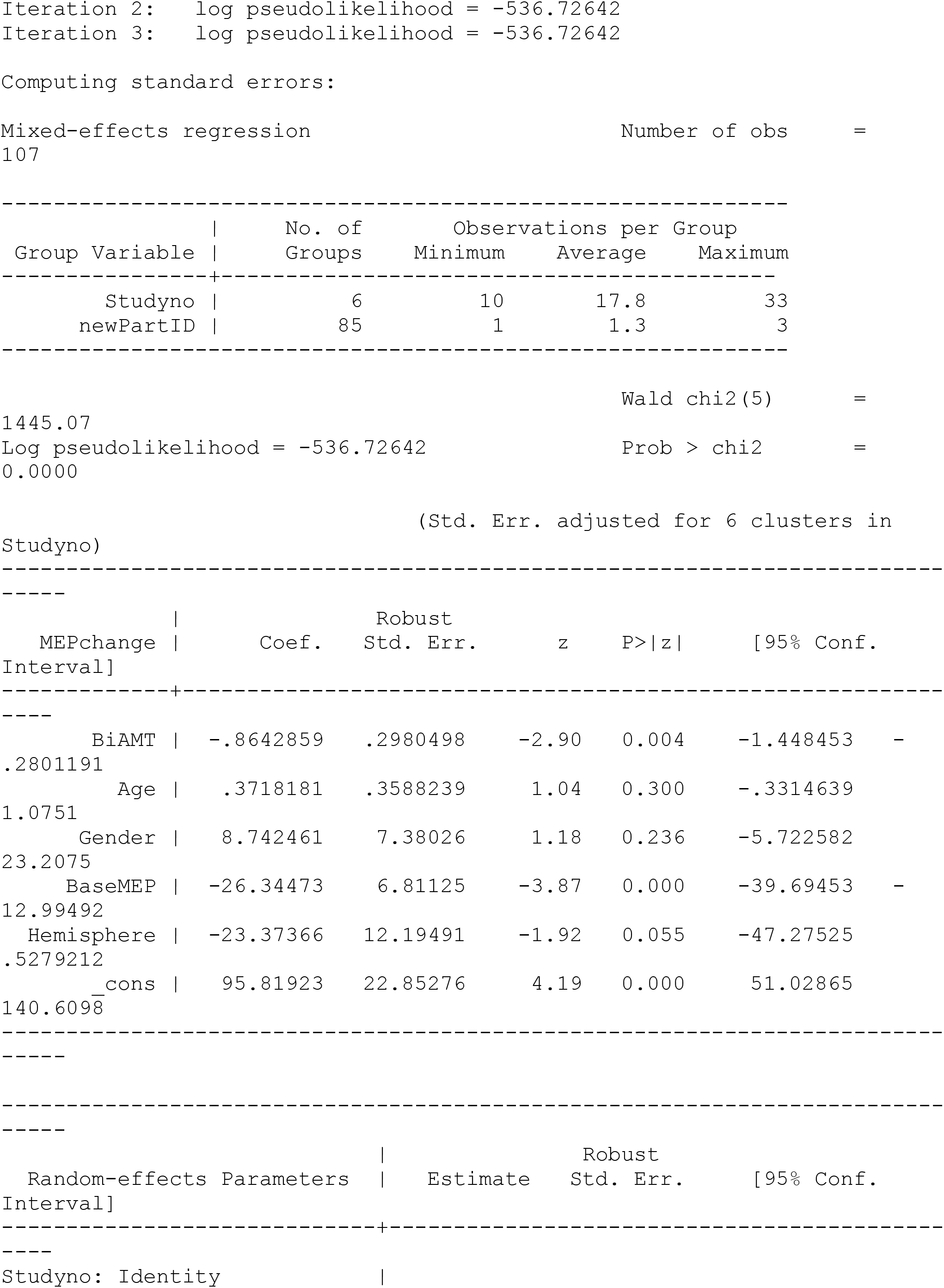
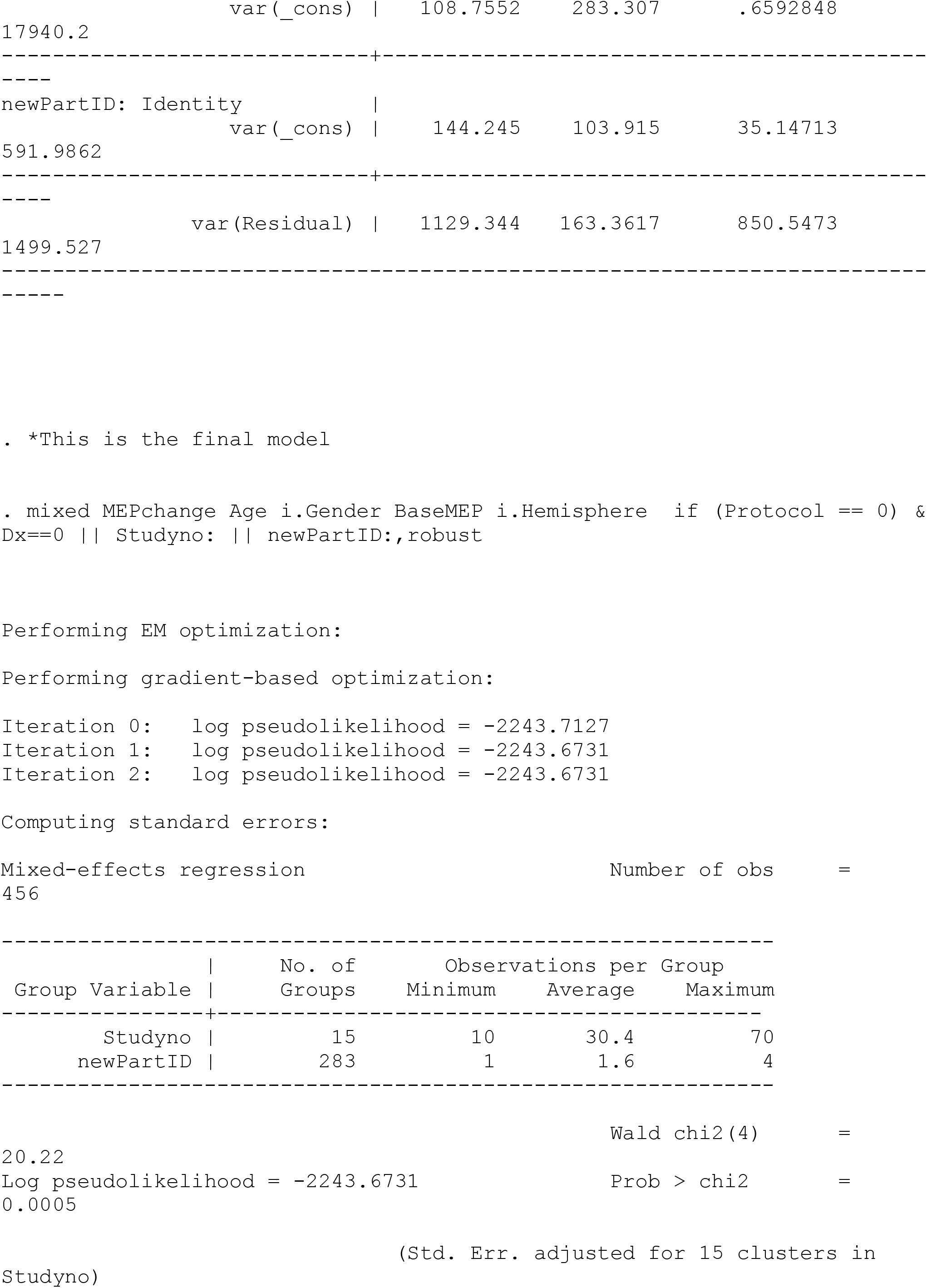
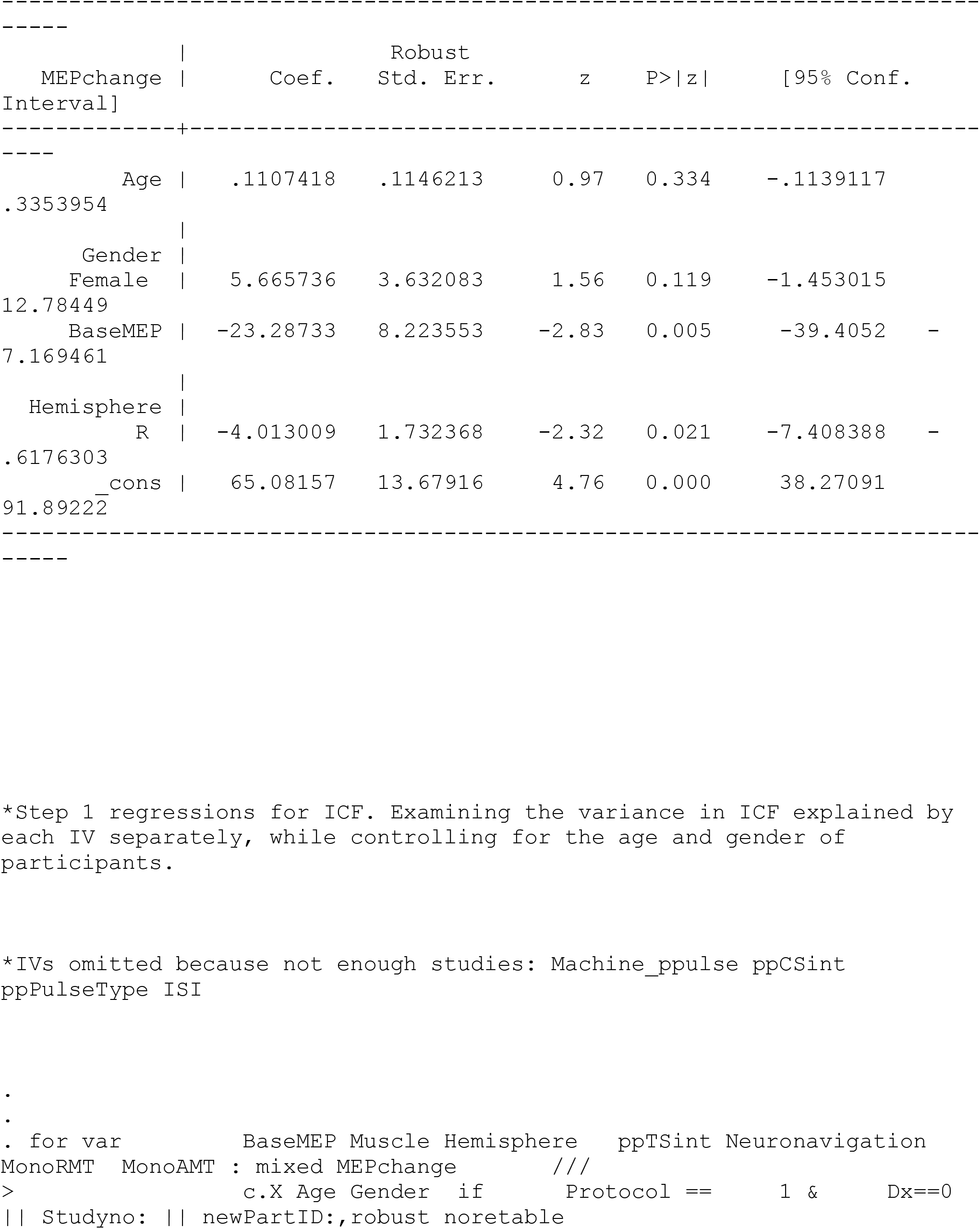
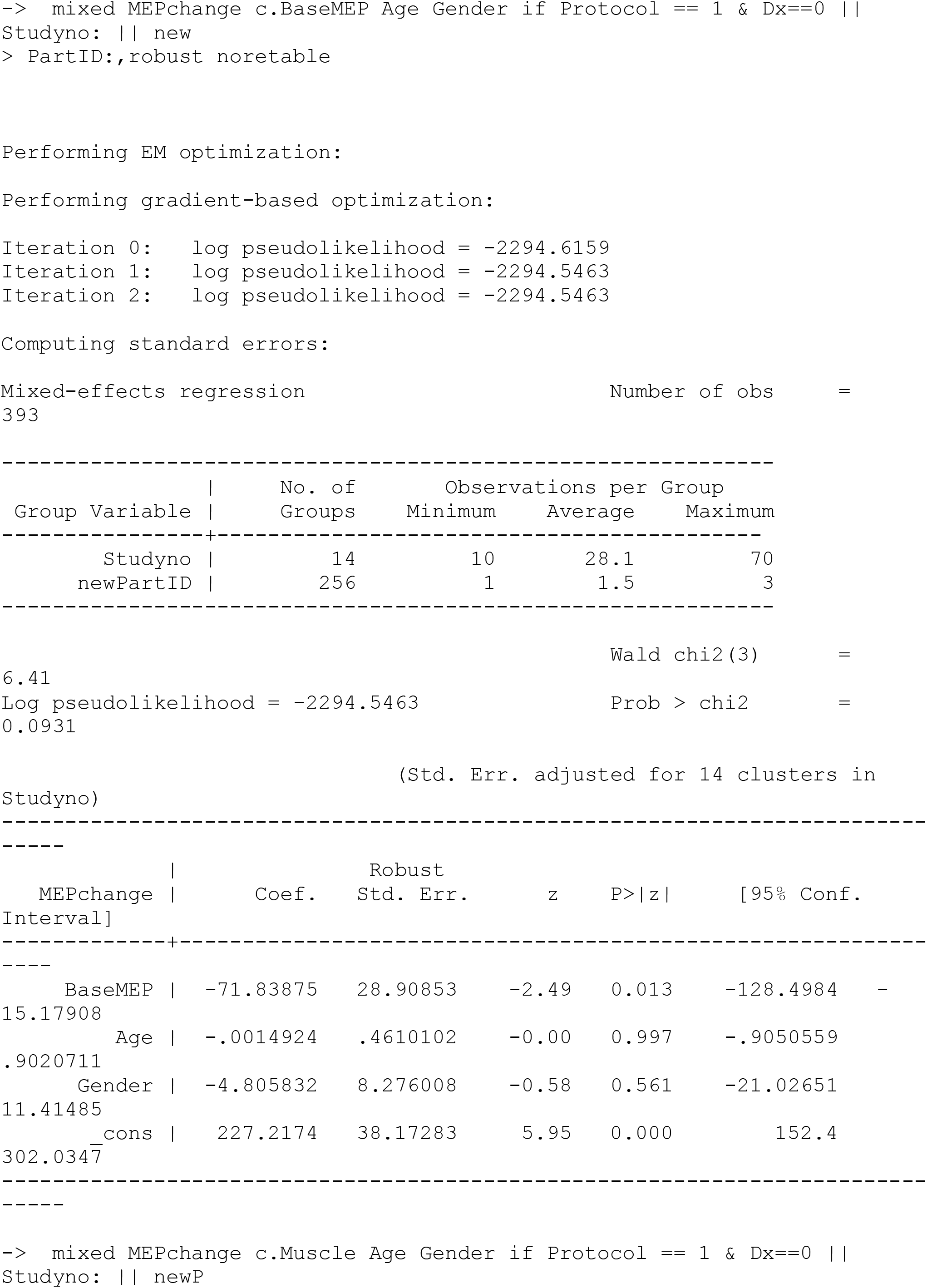
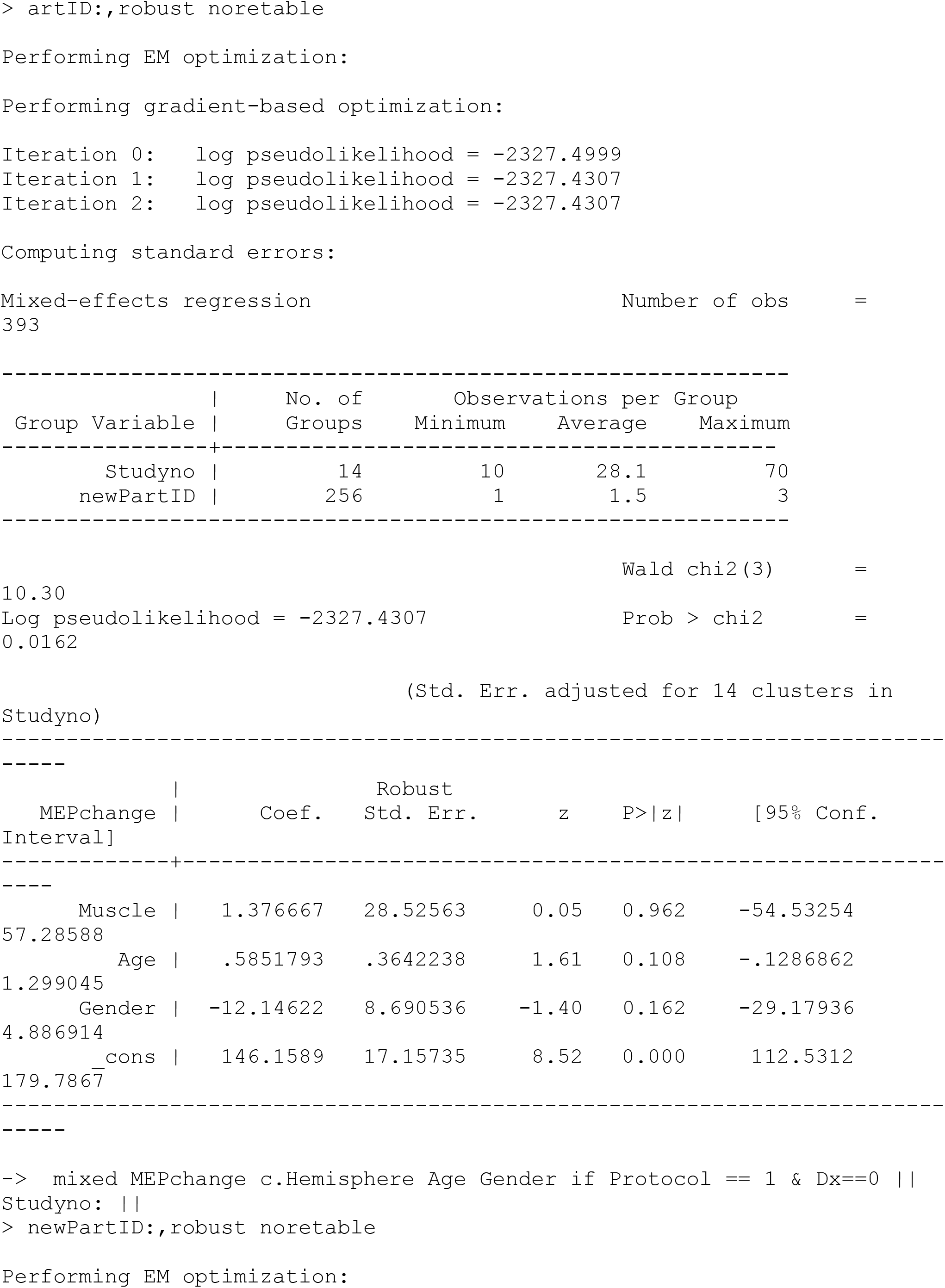
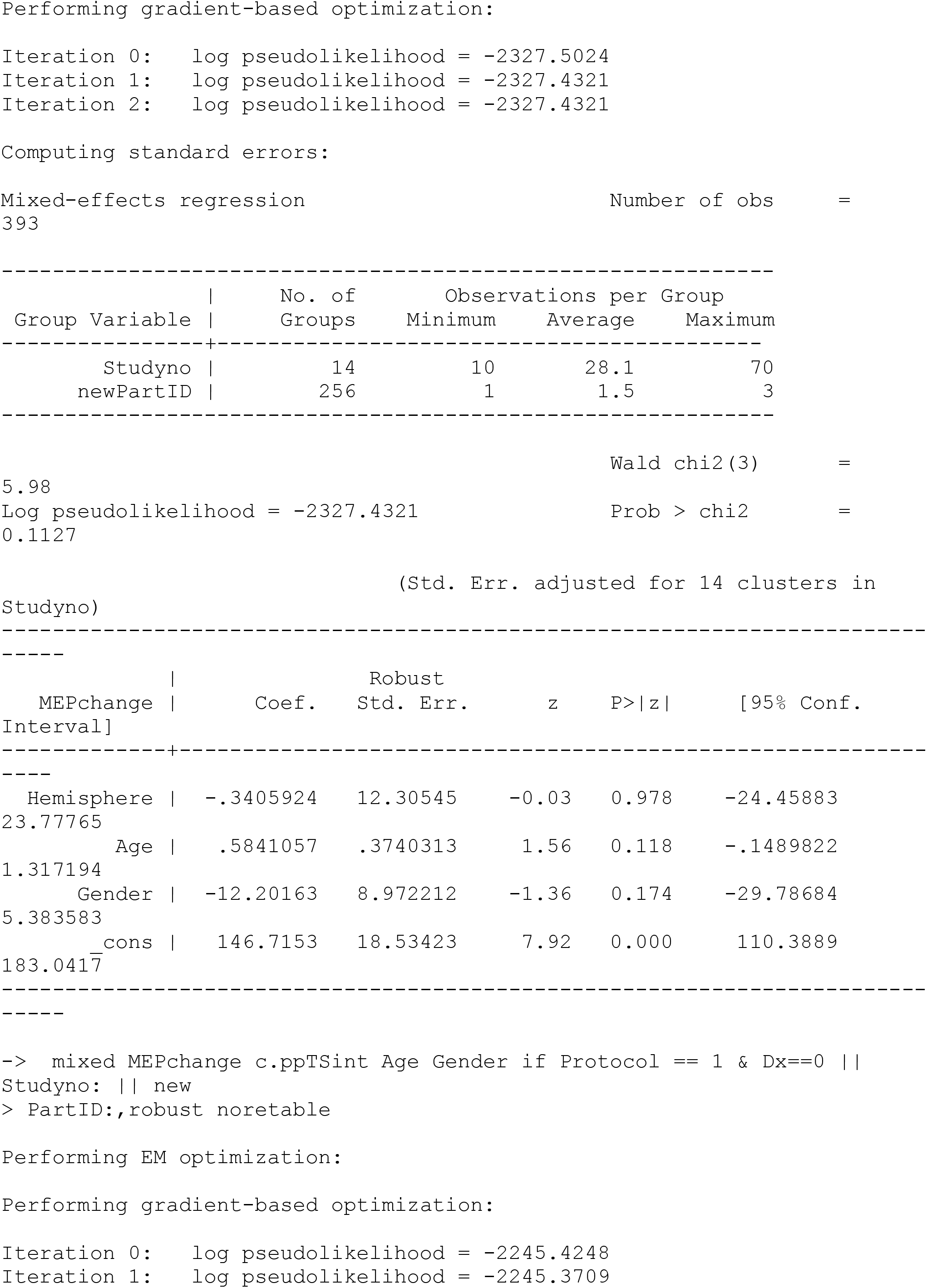
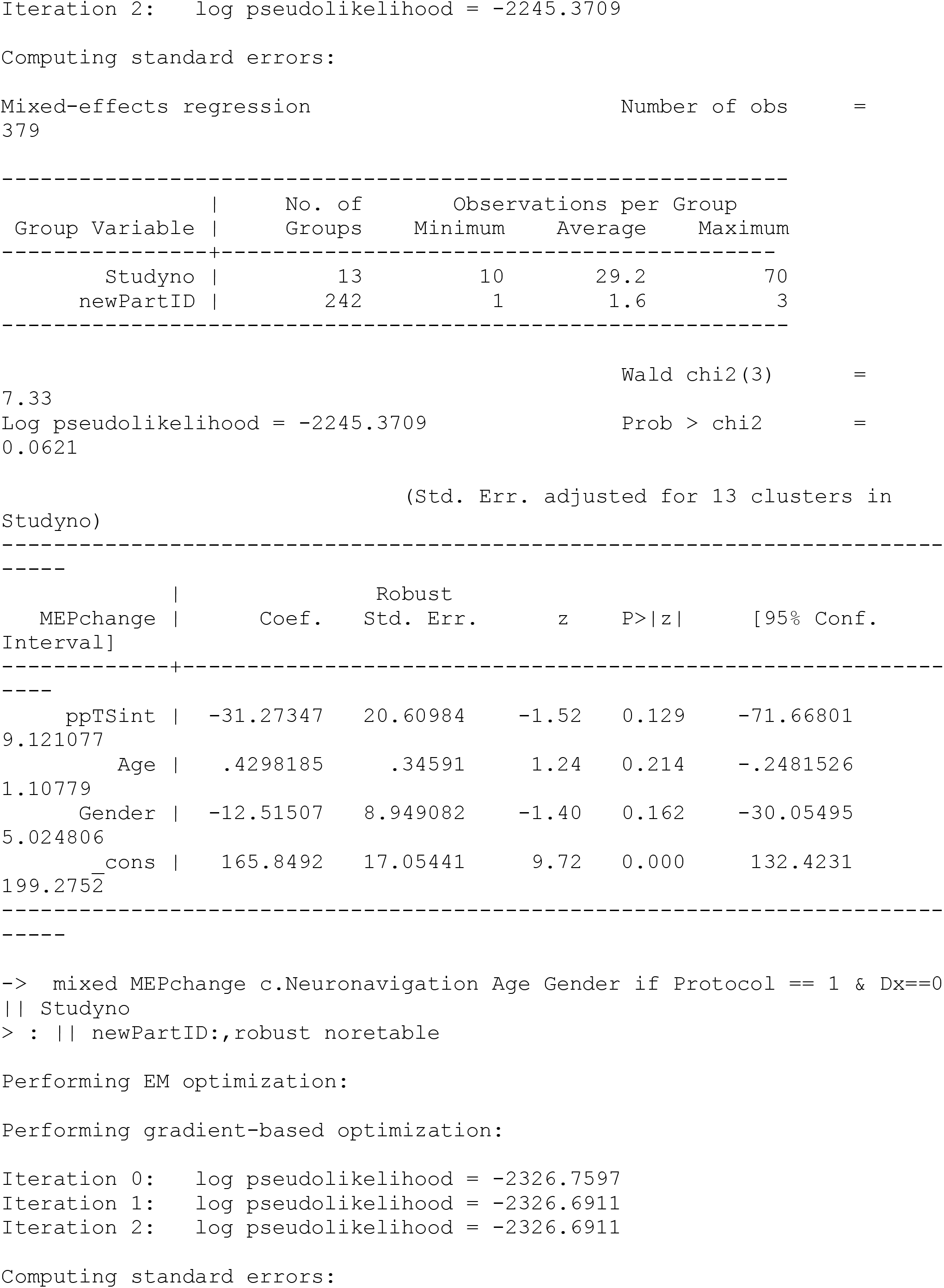
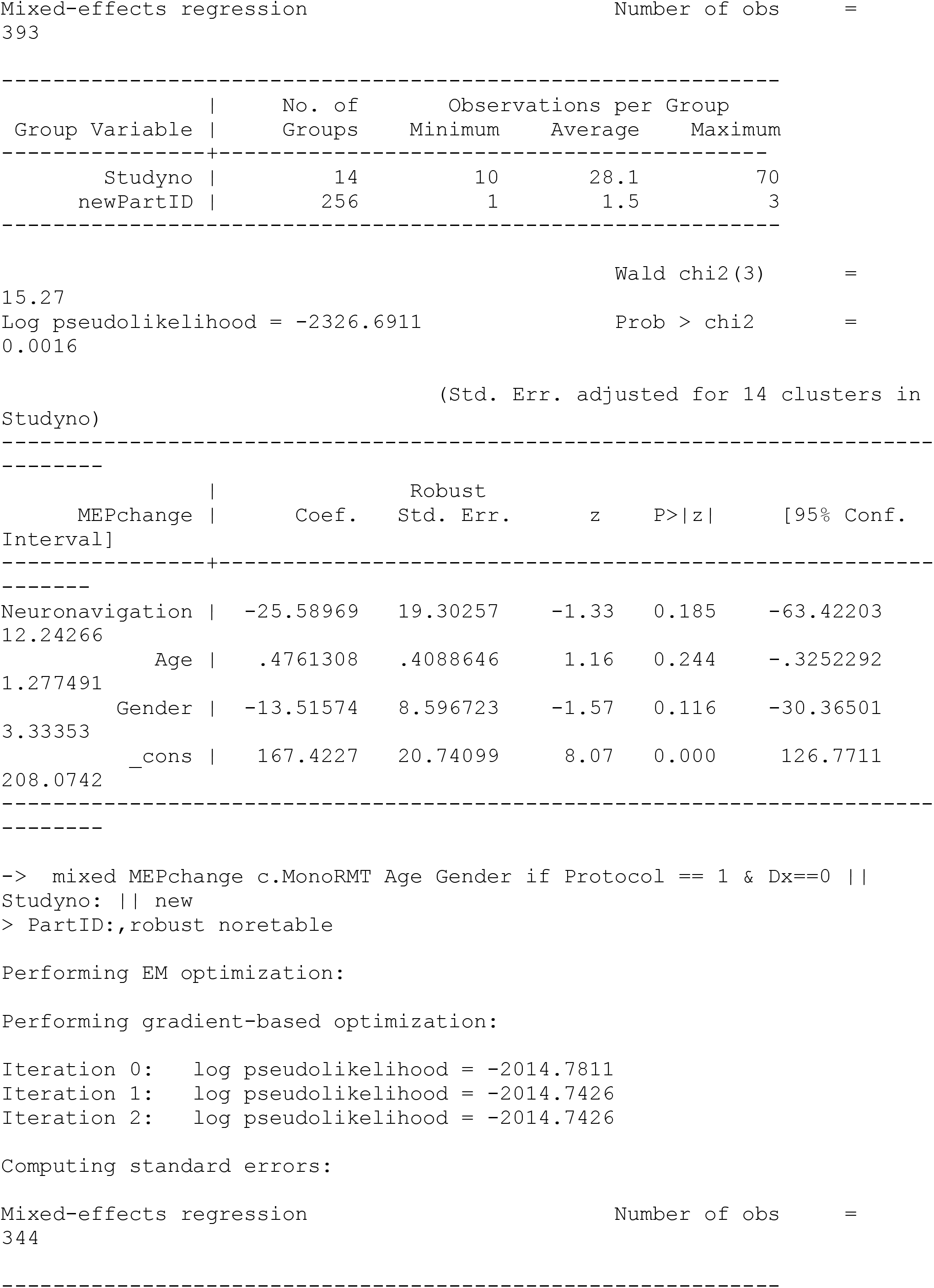
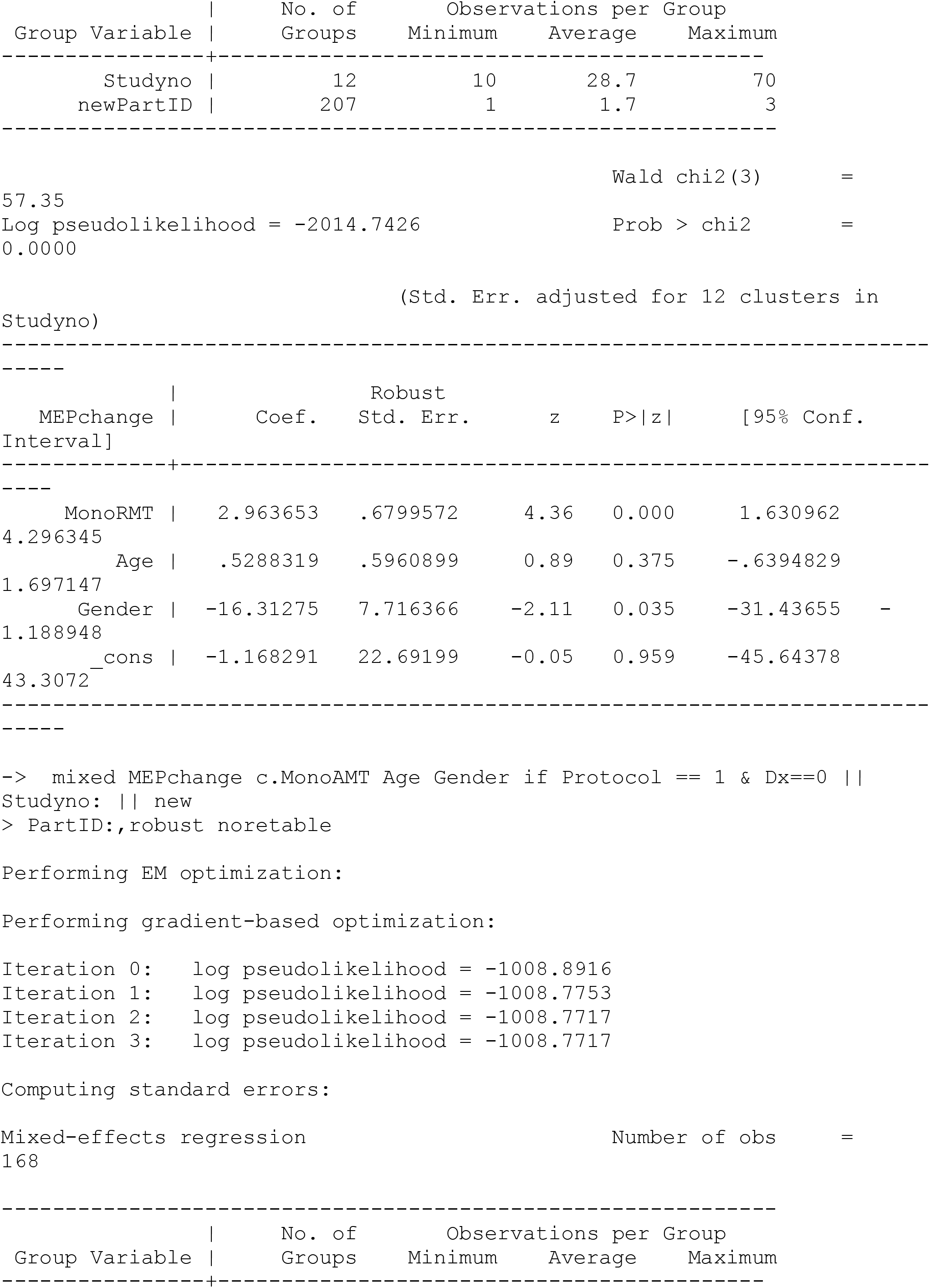
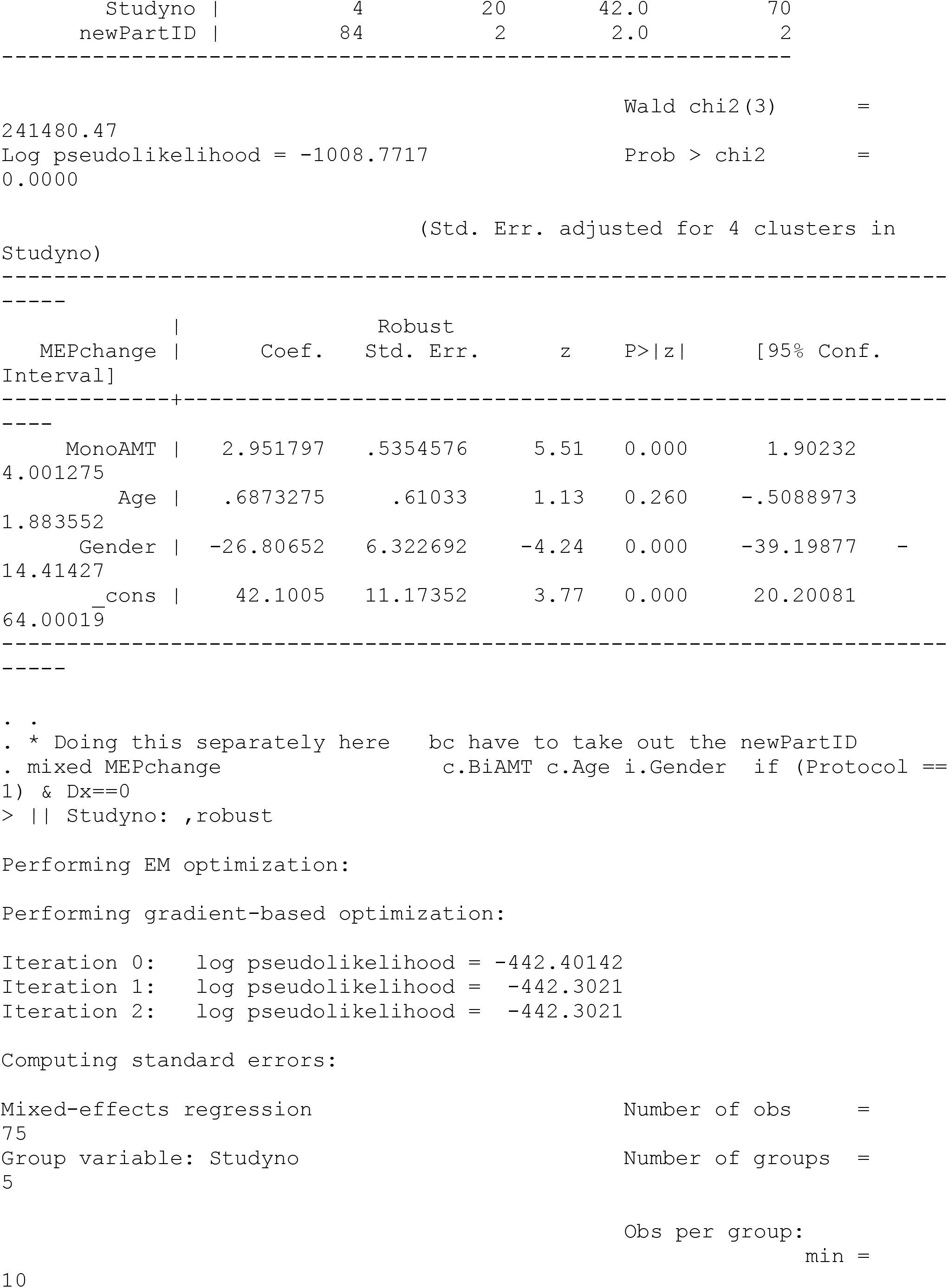
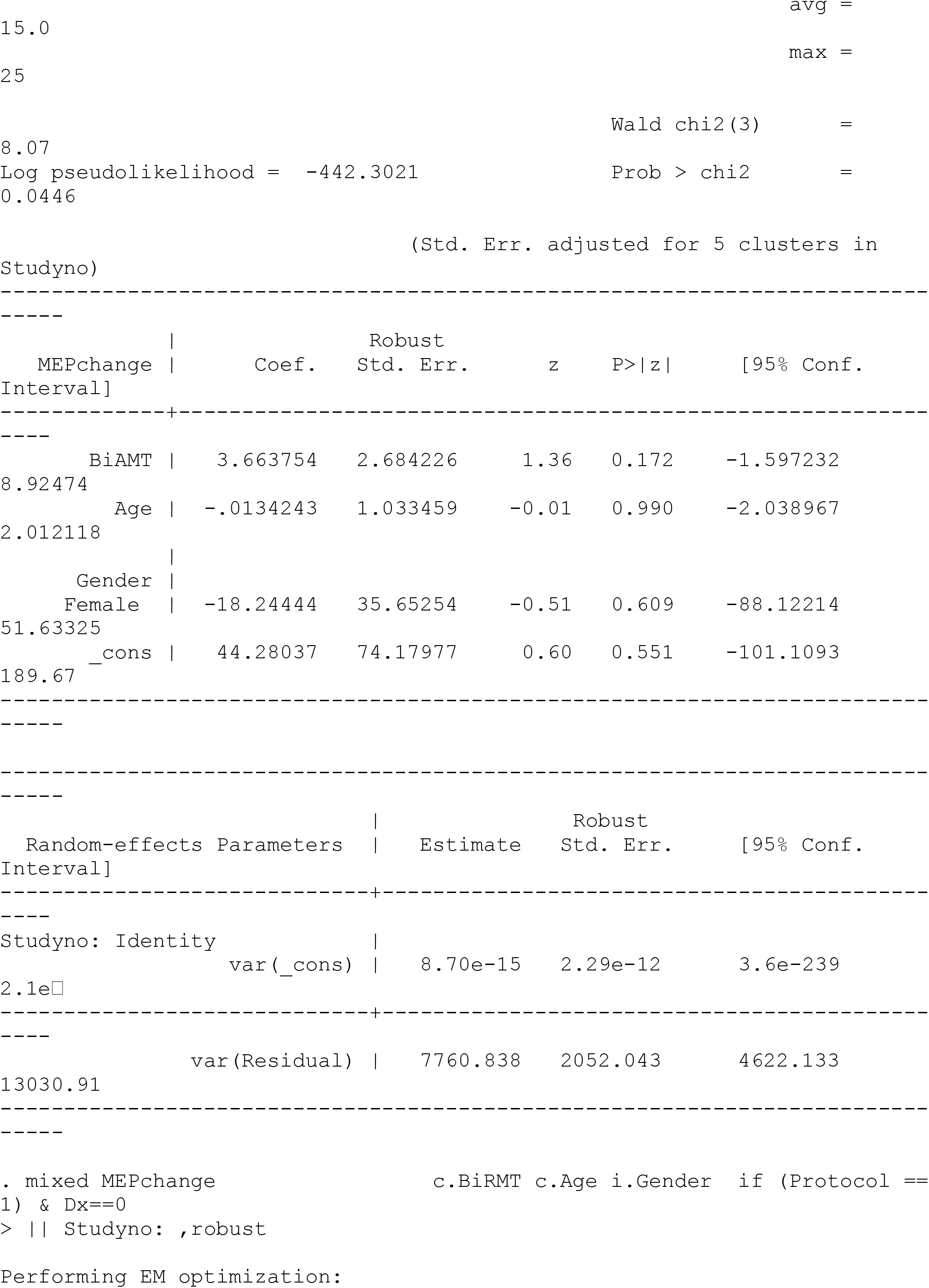
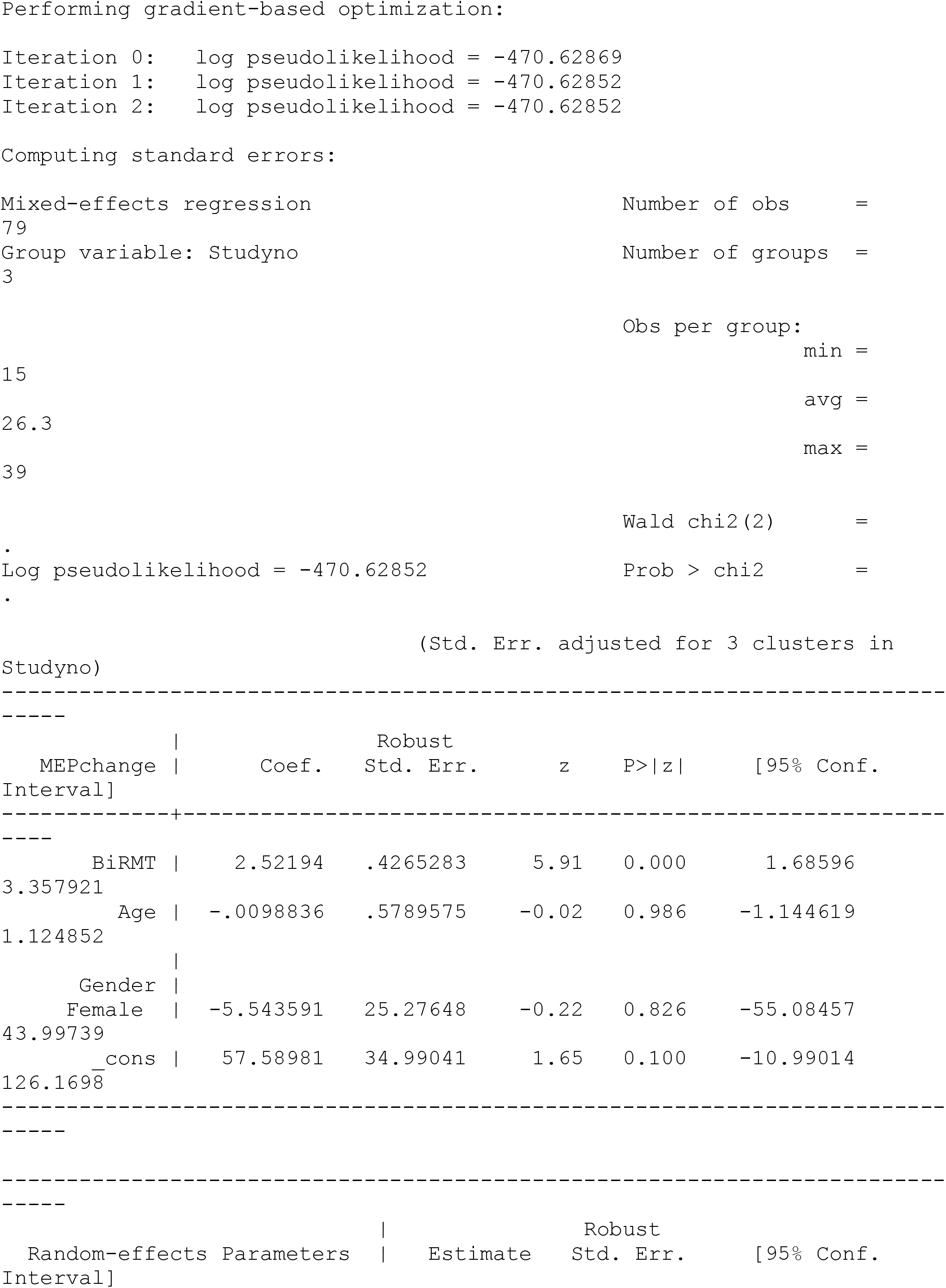
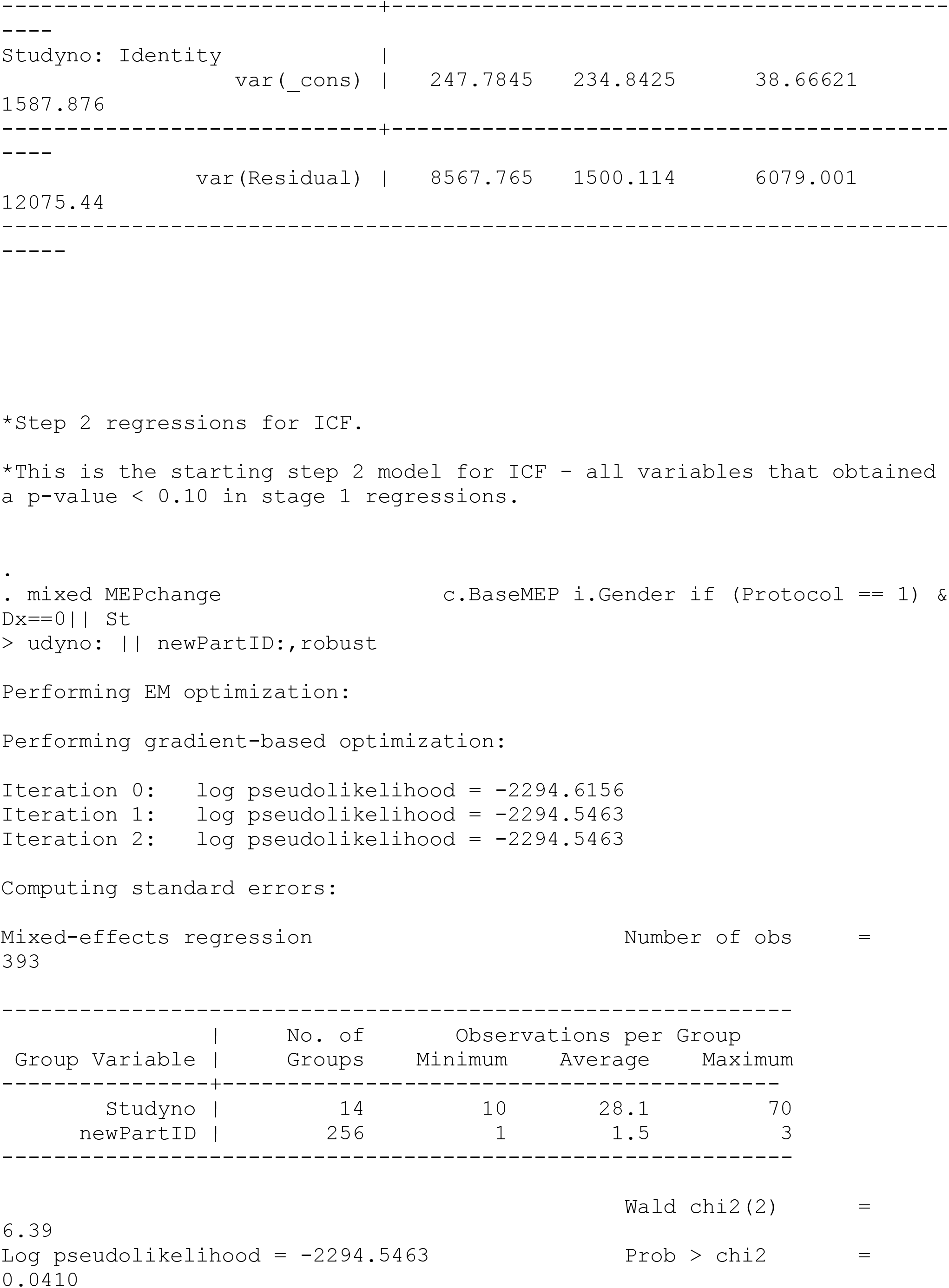
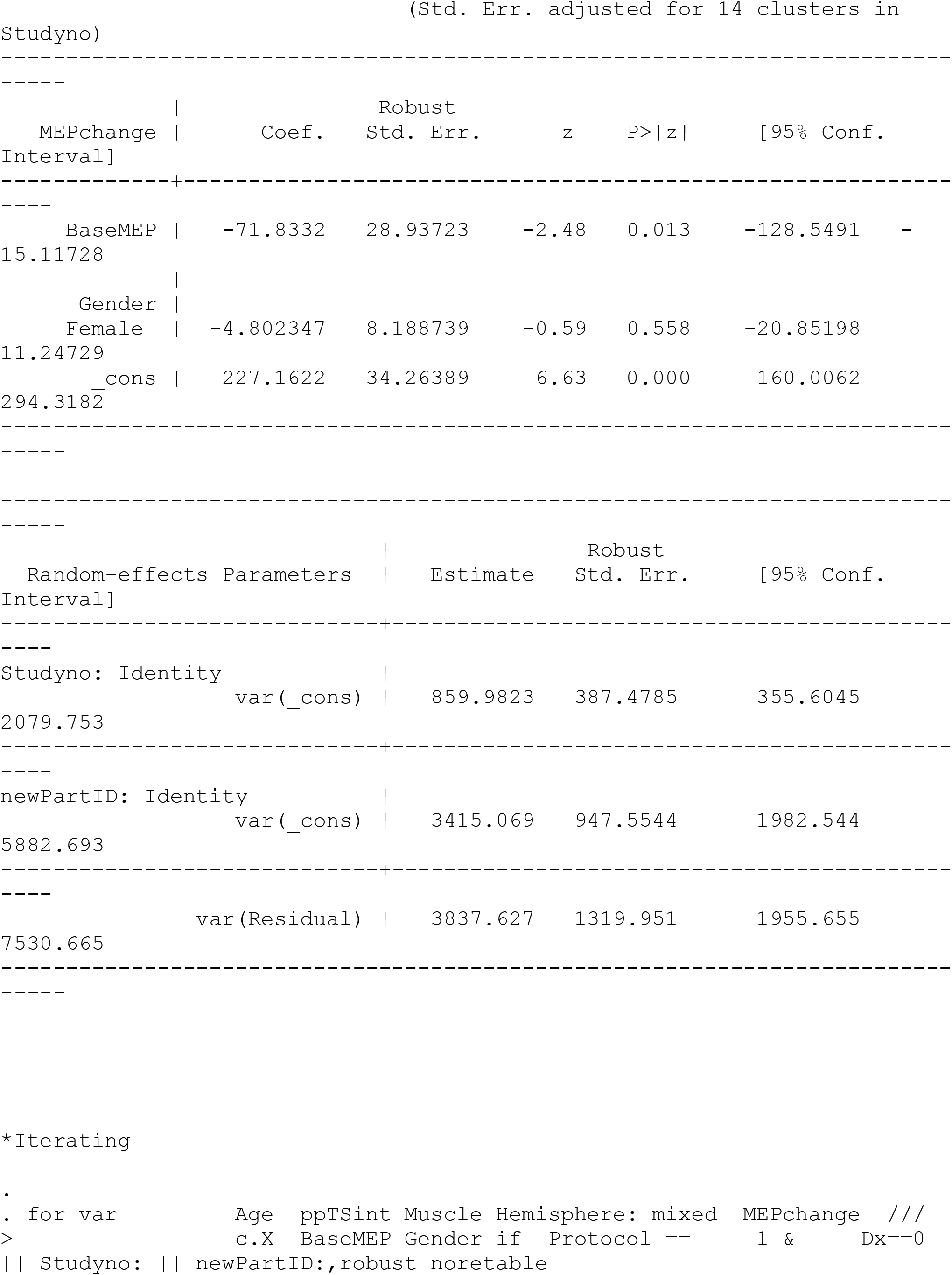
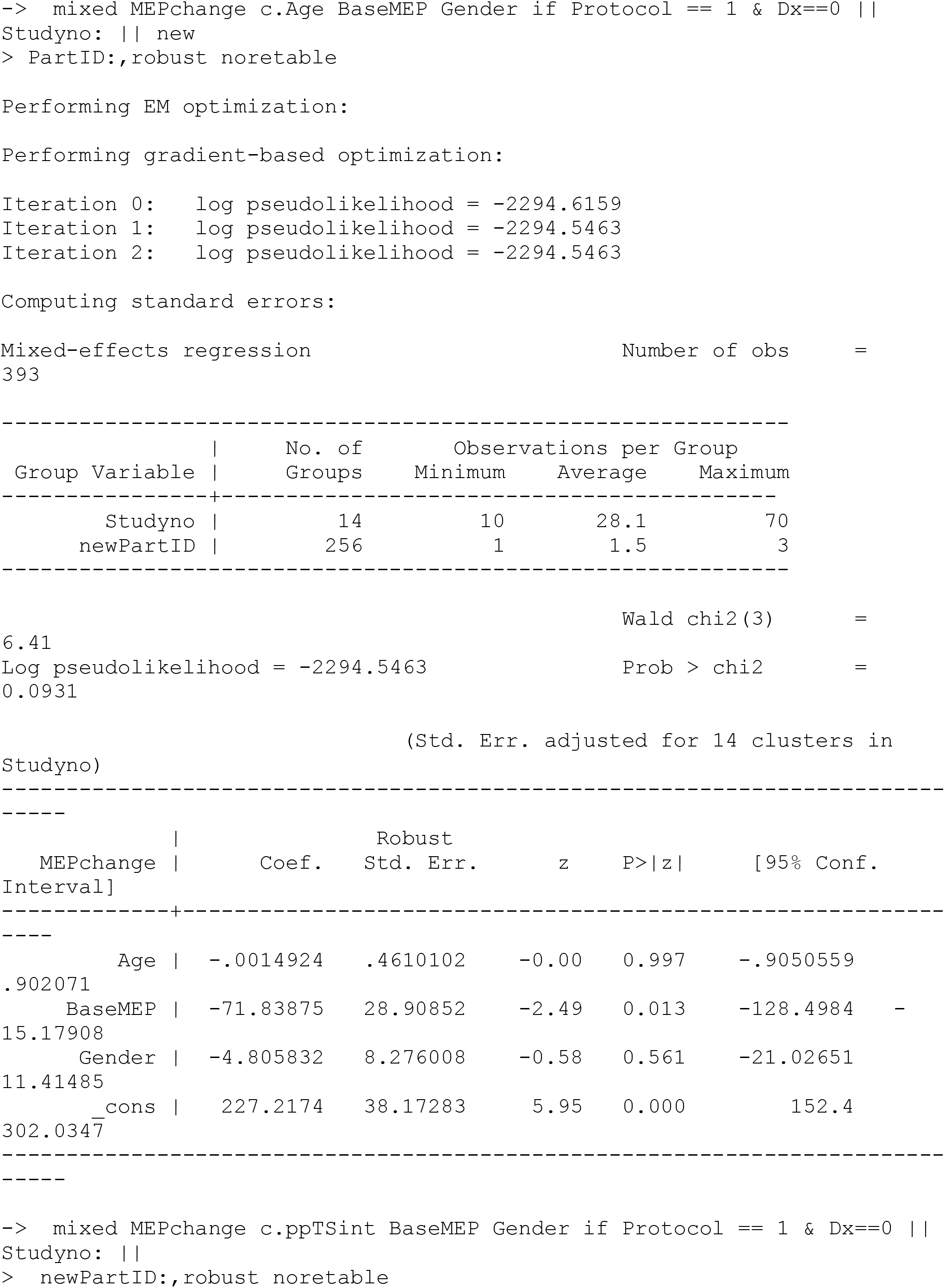
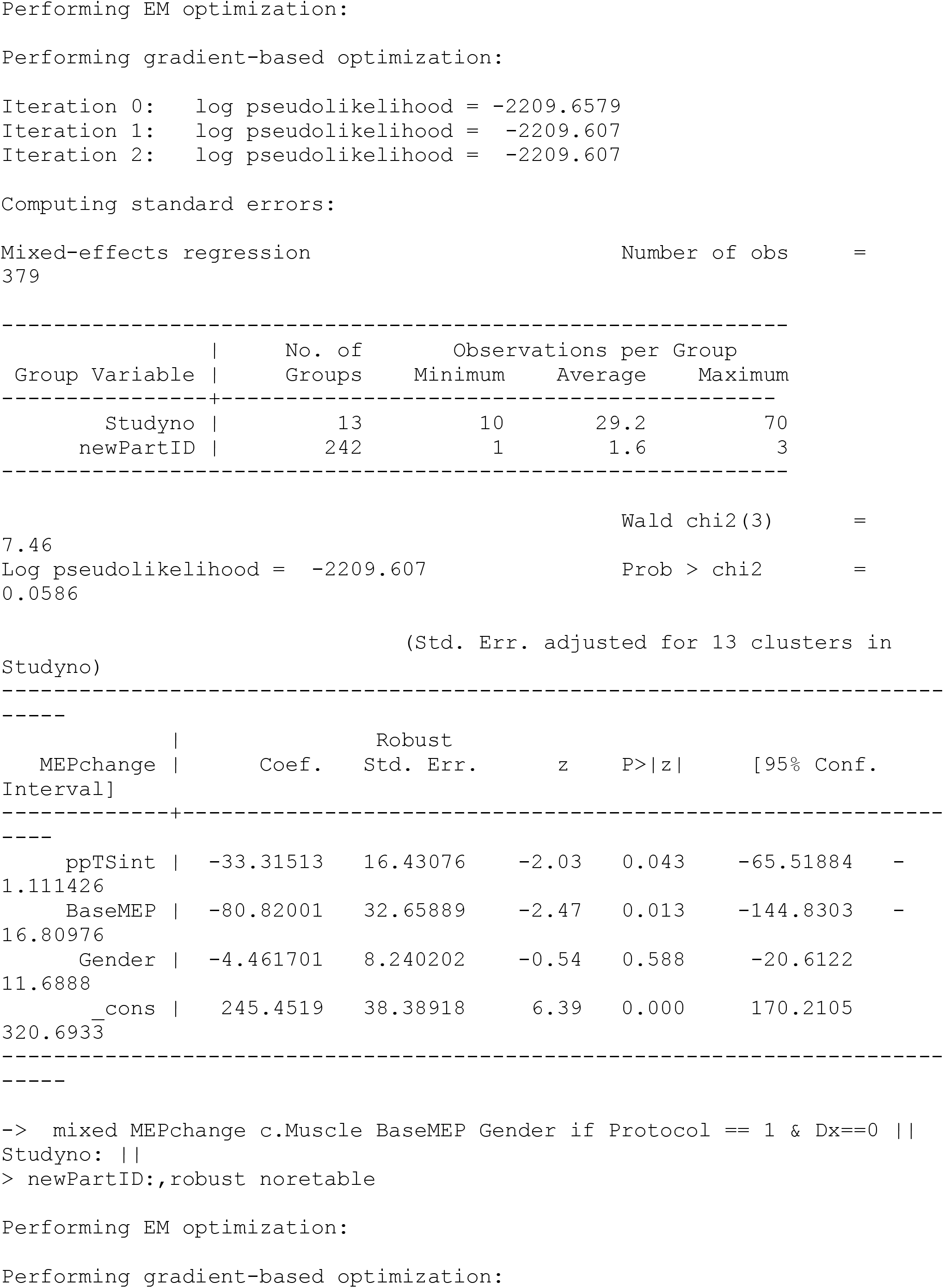
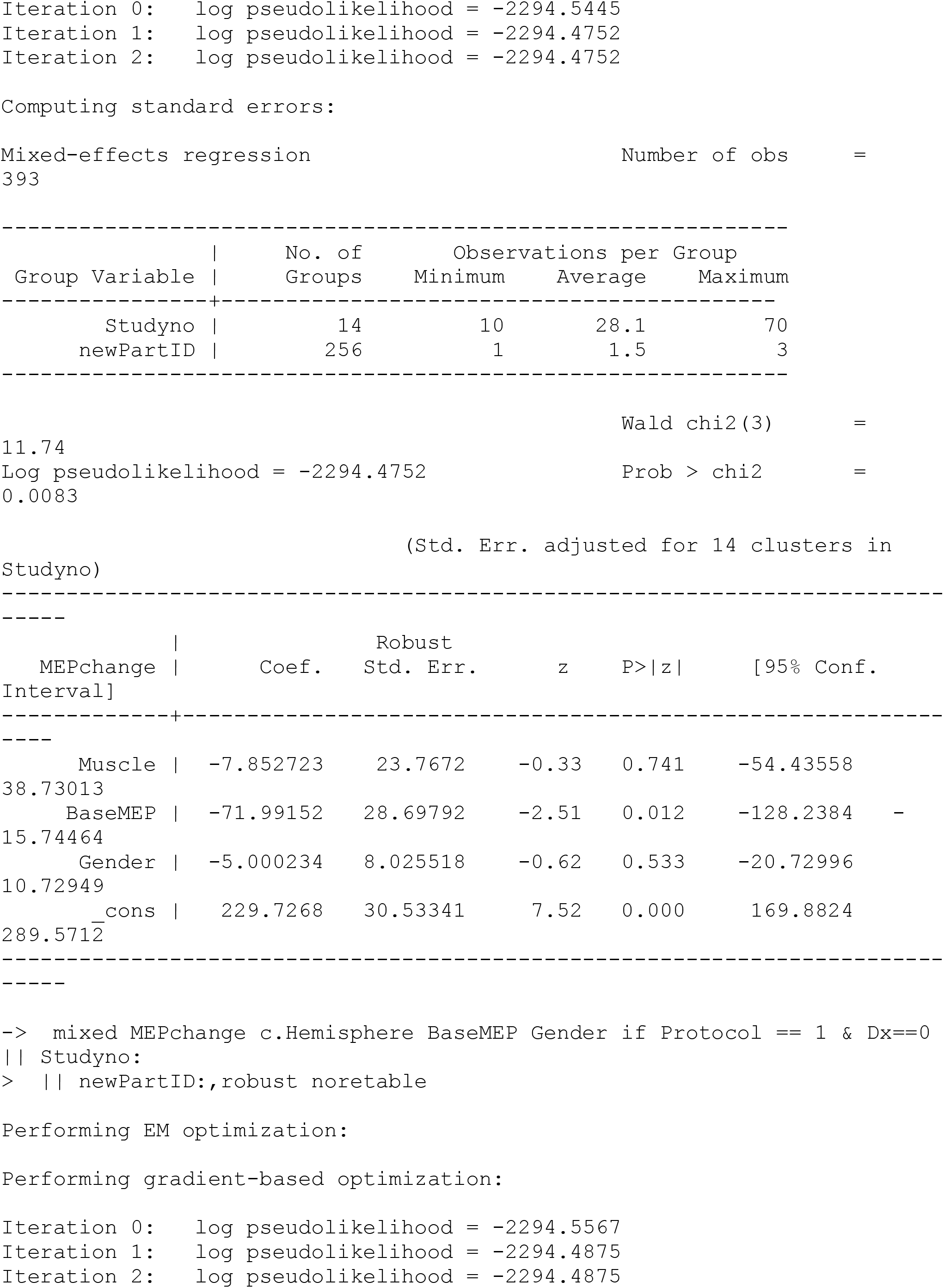
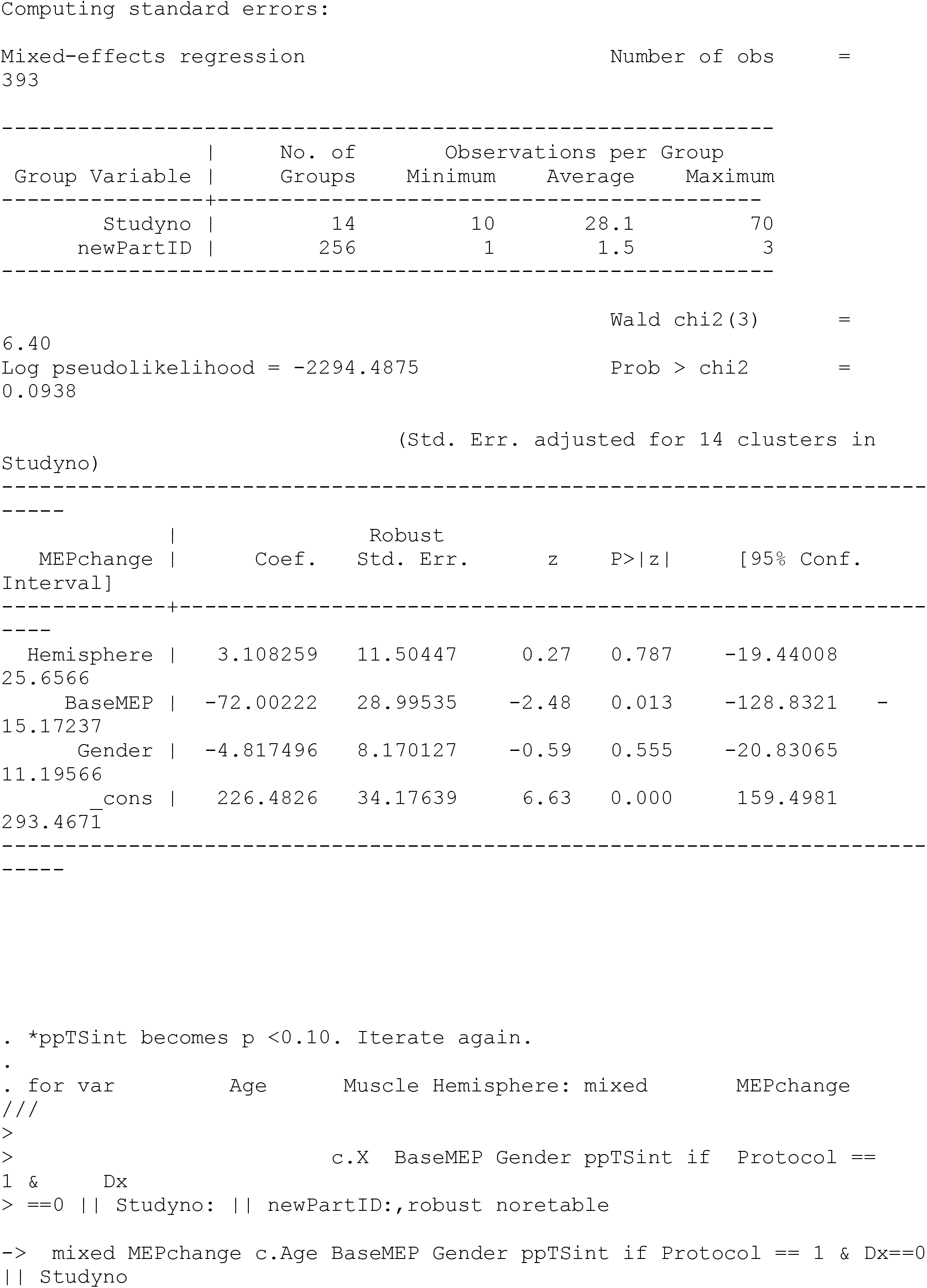
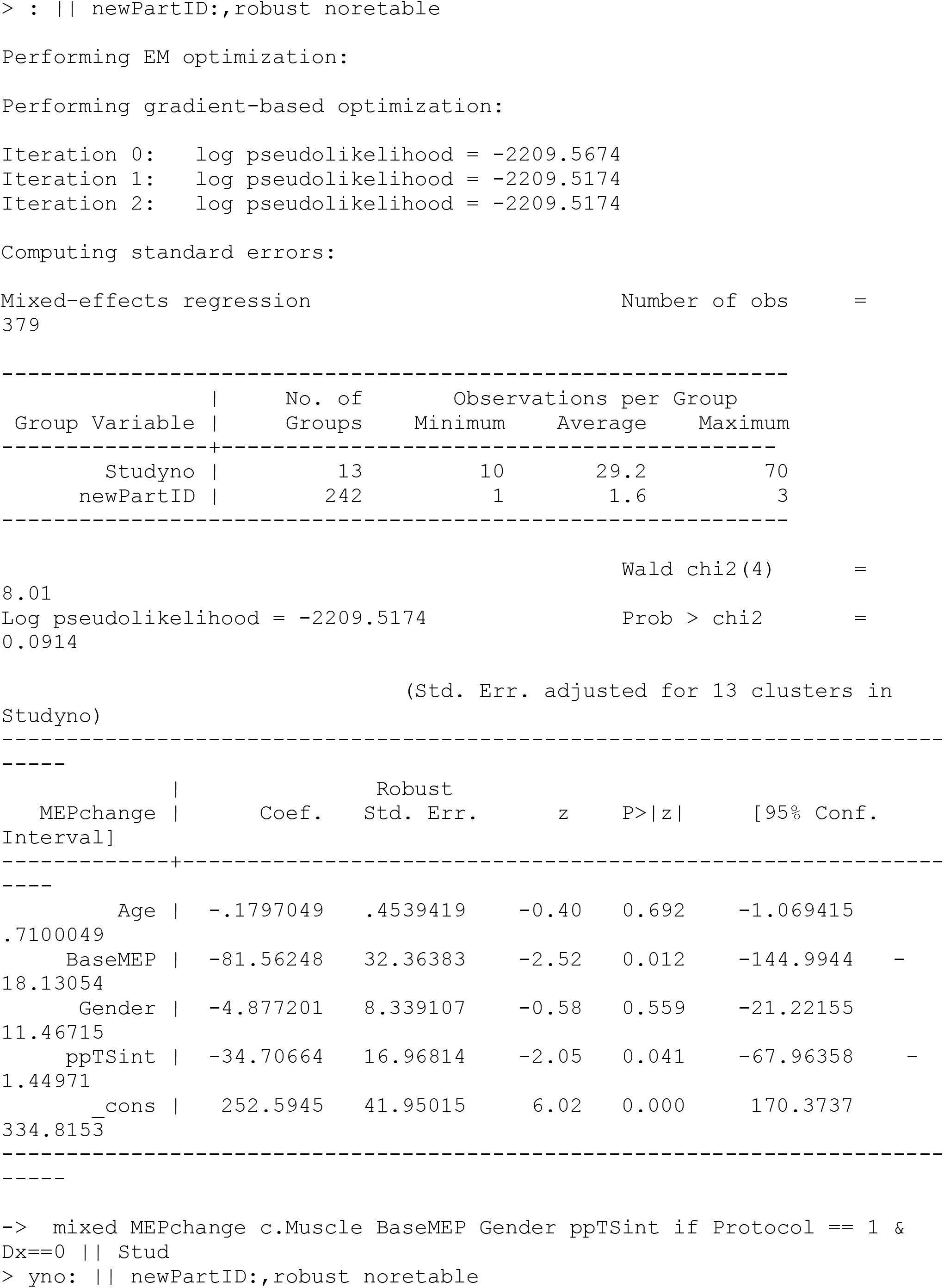
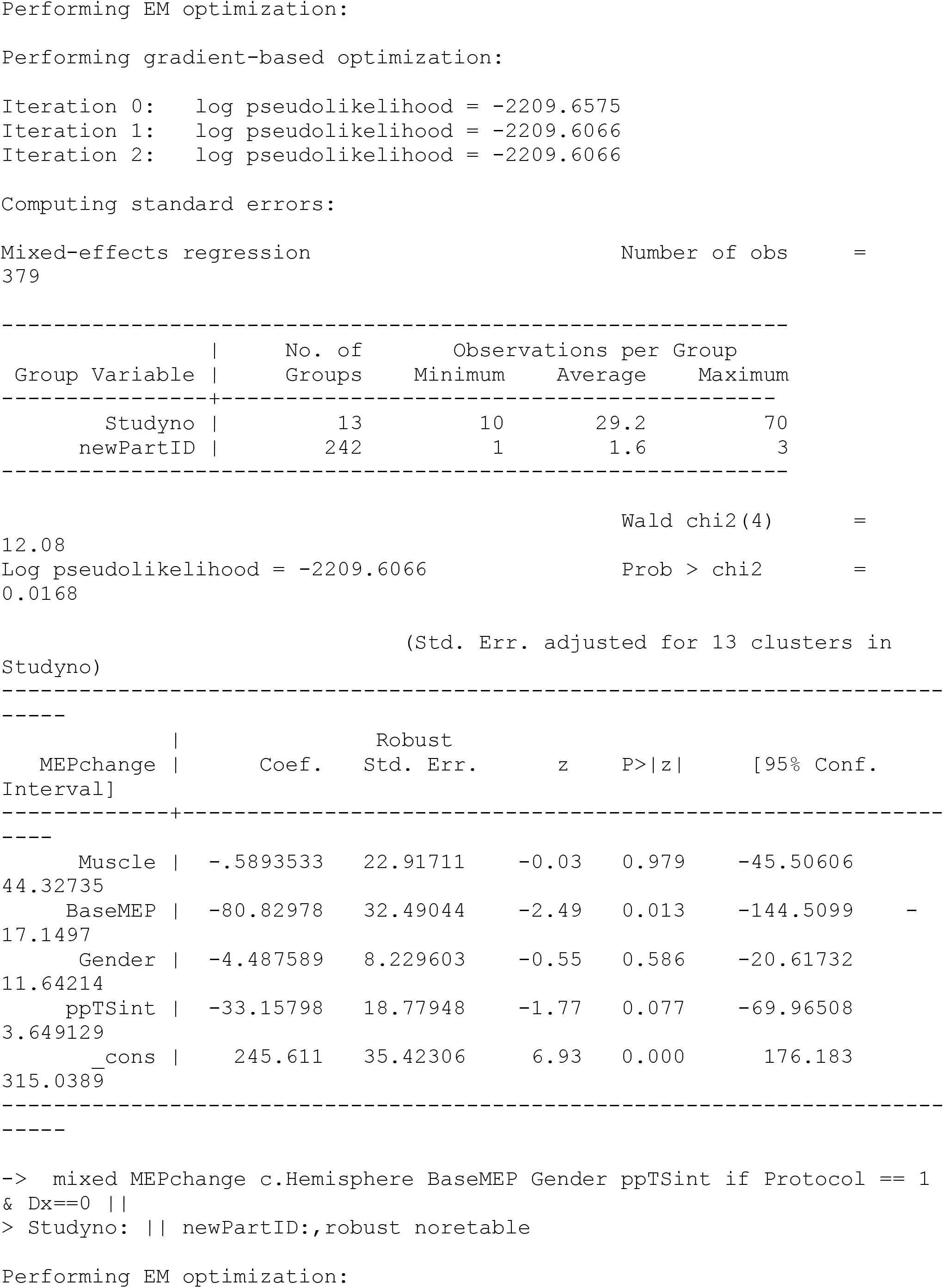
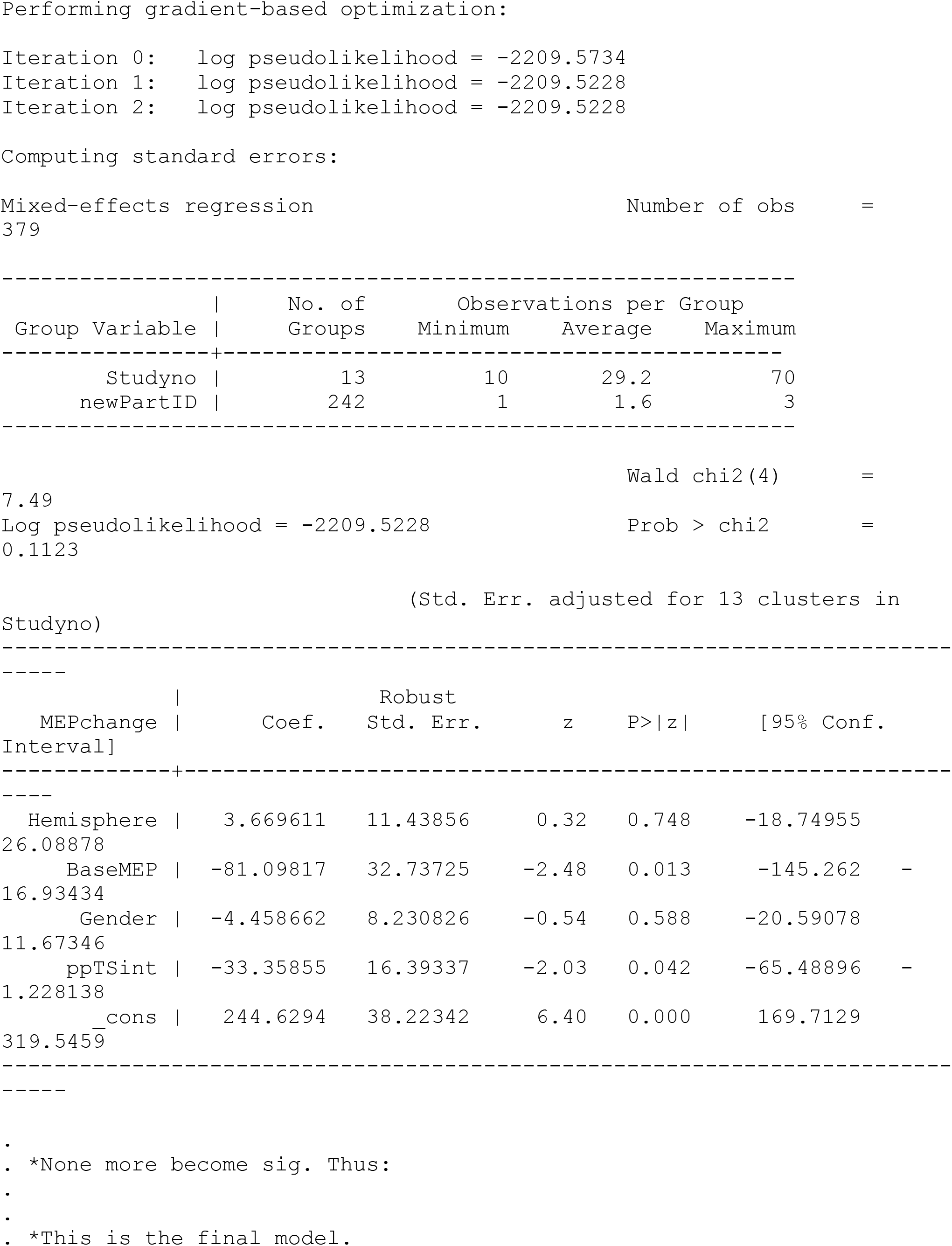
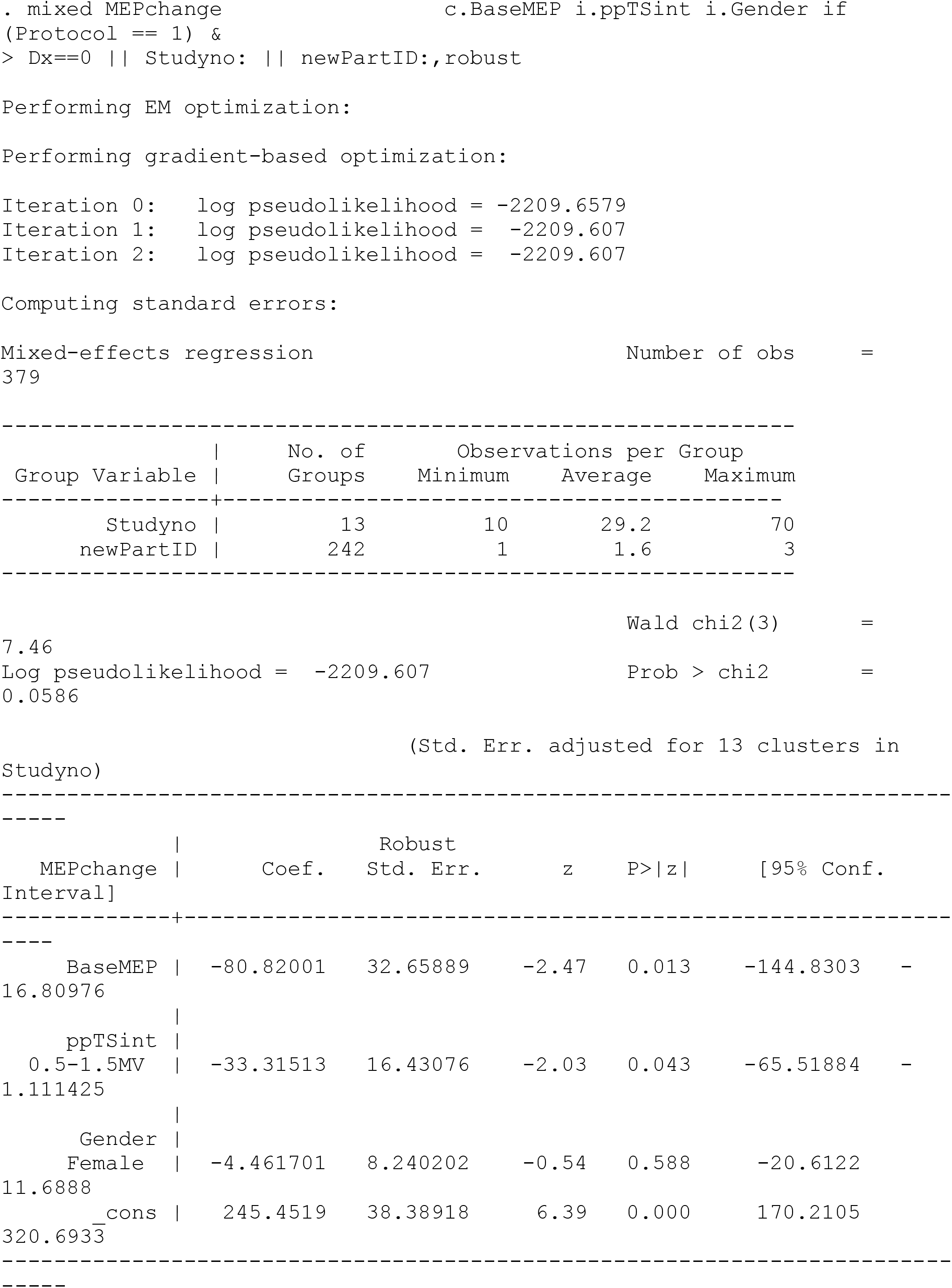

**Supplementary file 9. Non-linear relationships for ICF.** Post-hoc analyses demonstrated significant non-linear relationships between ICF normalised MEP and monophasic AMT and biphasic RMT.

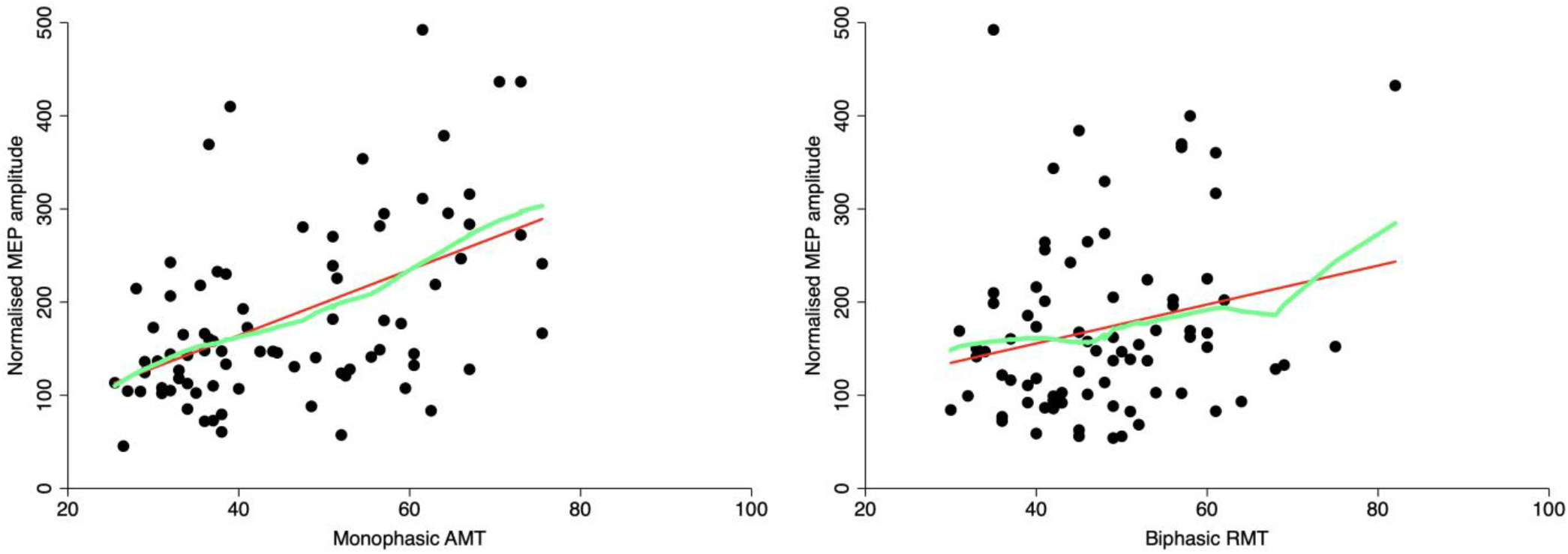

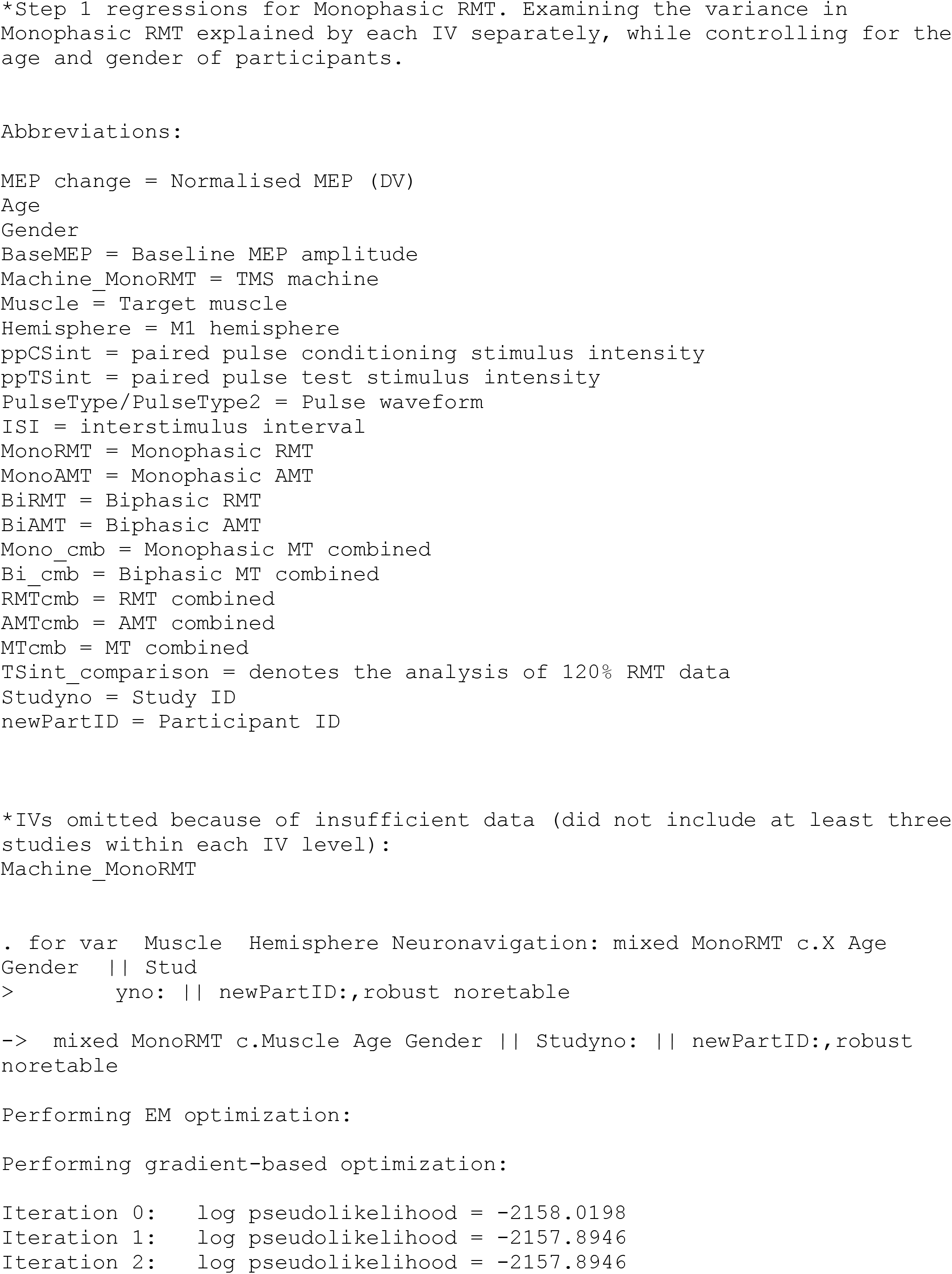
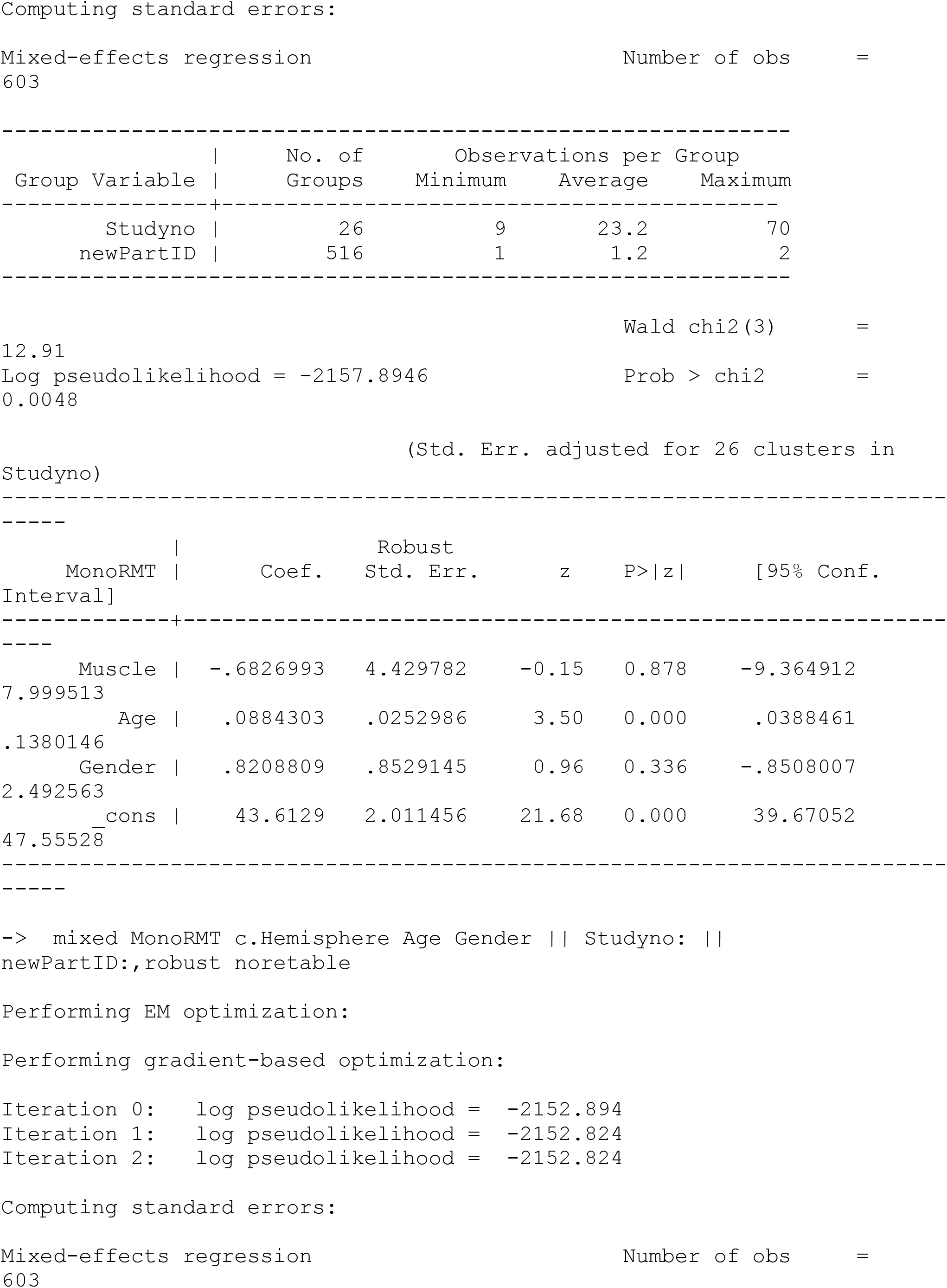
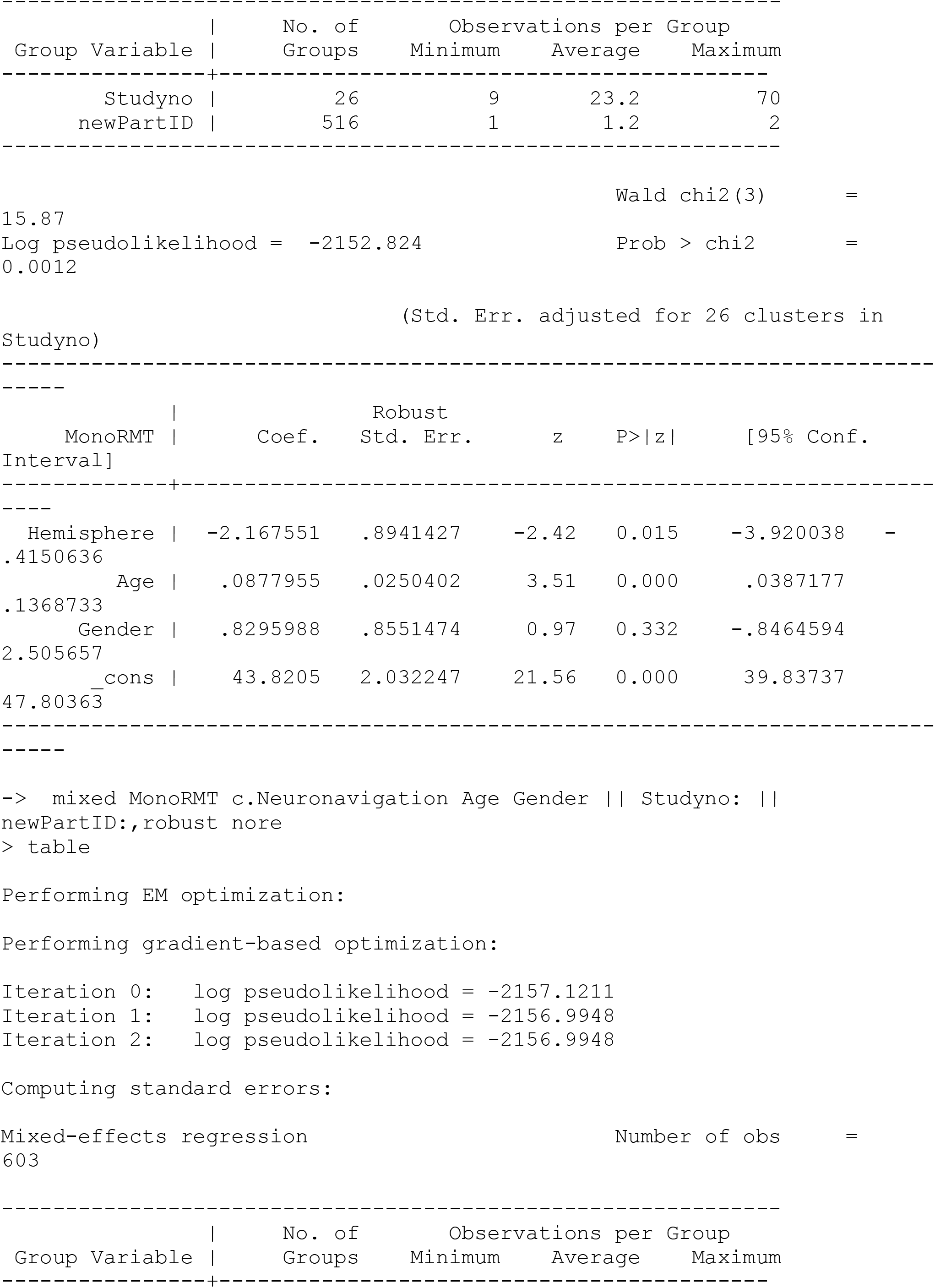
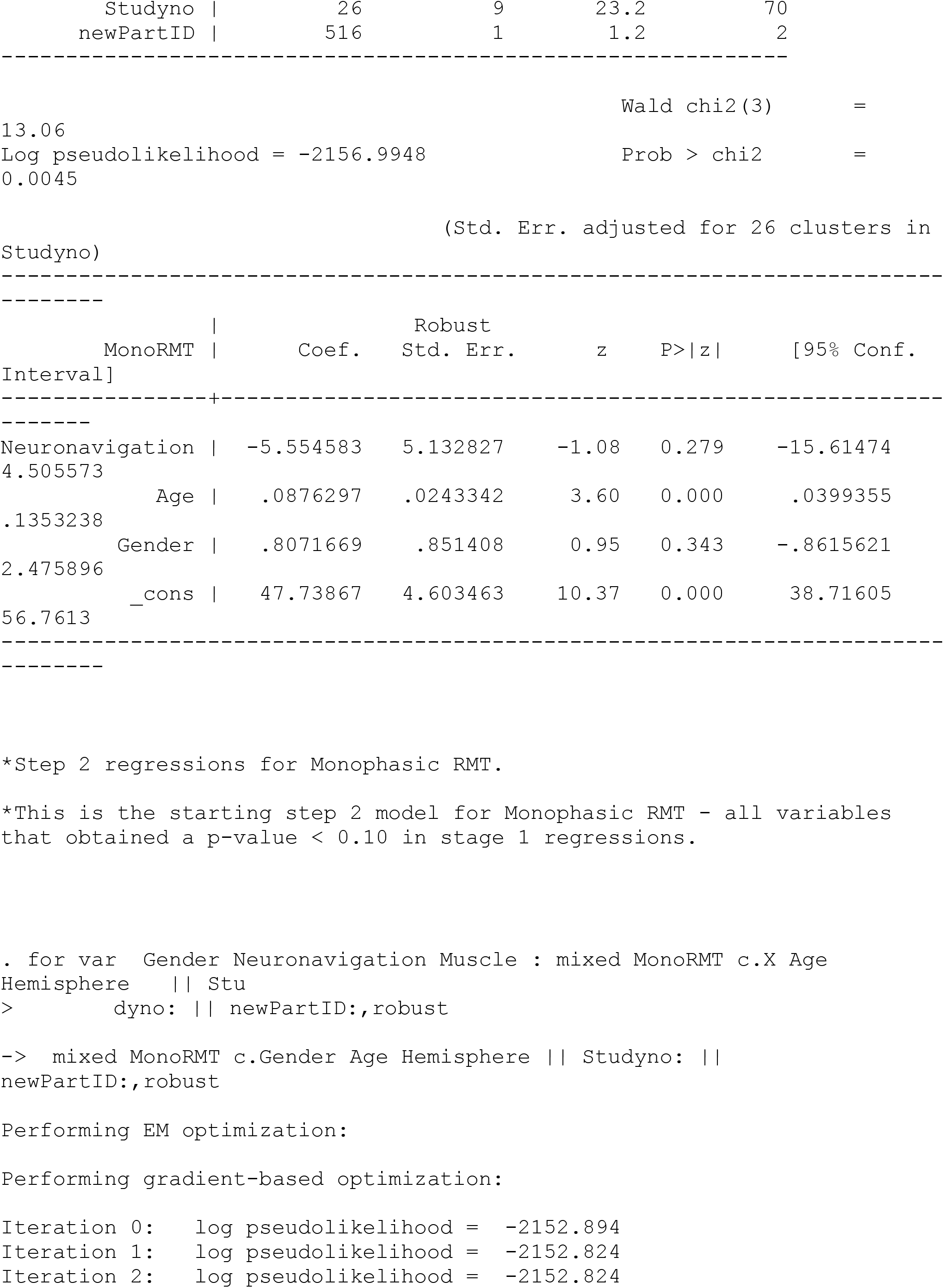
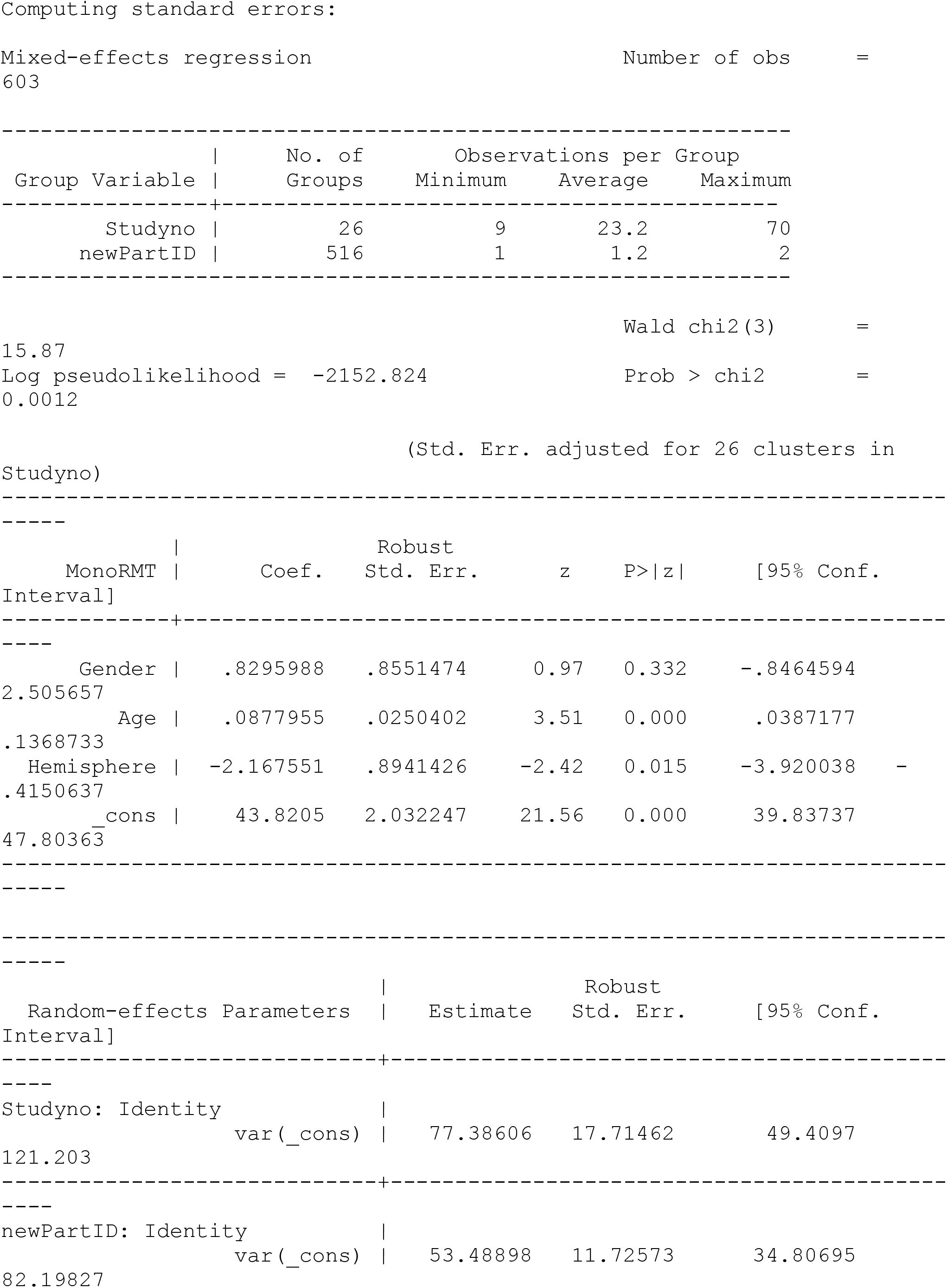
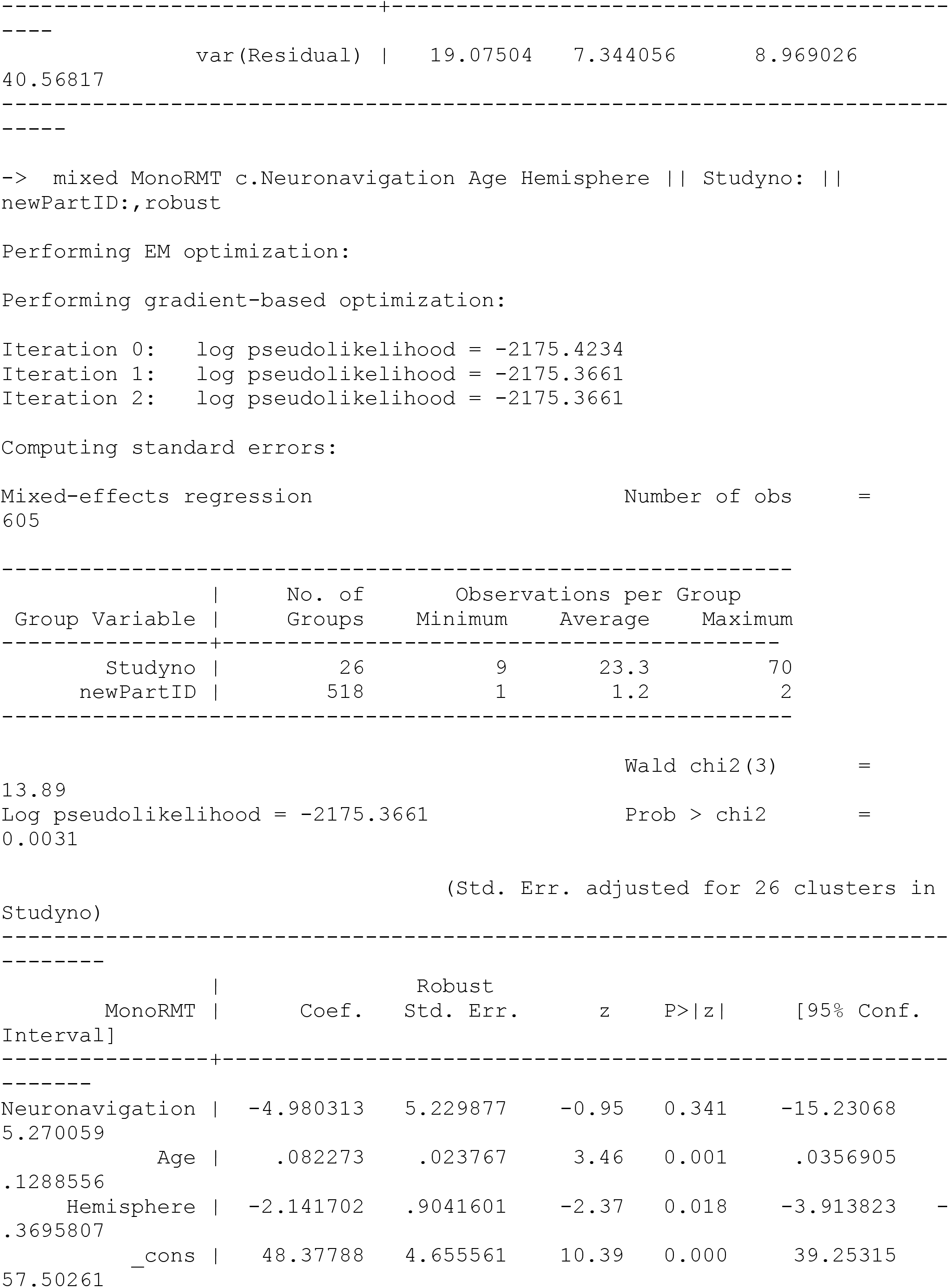
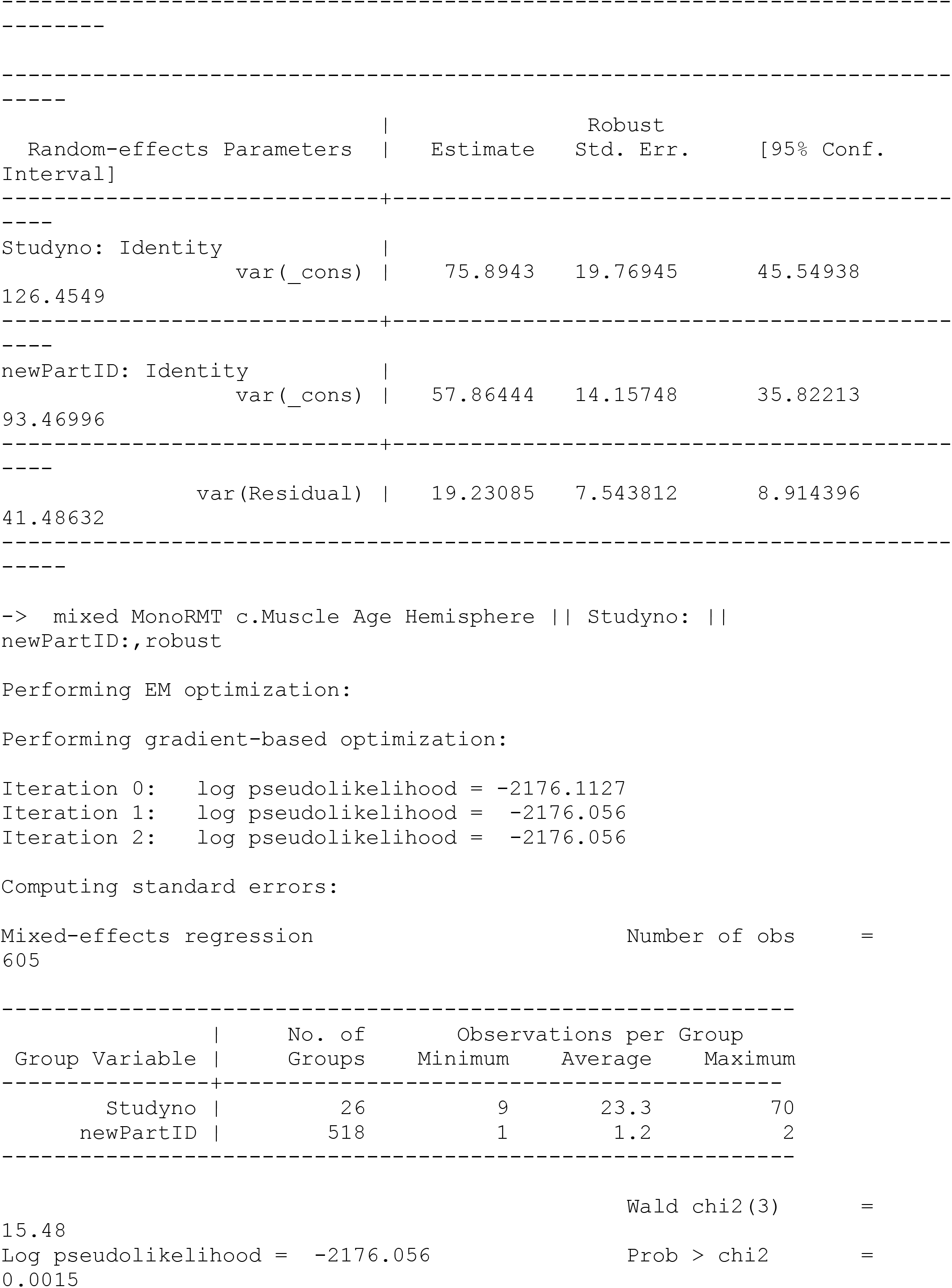
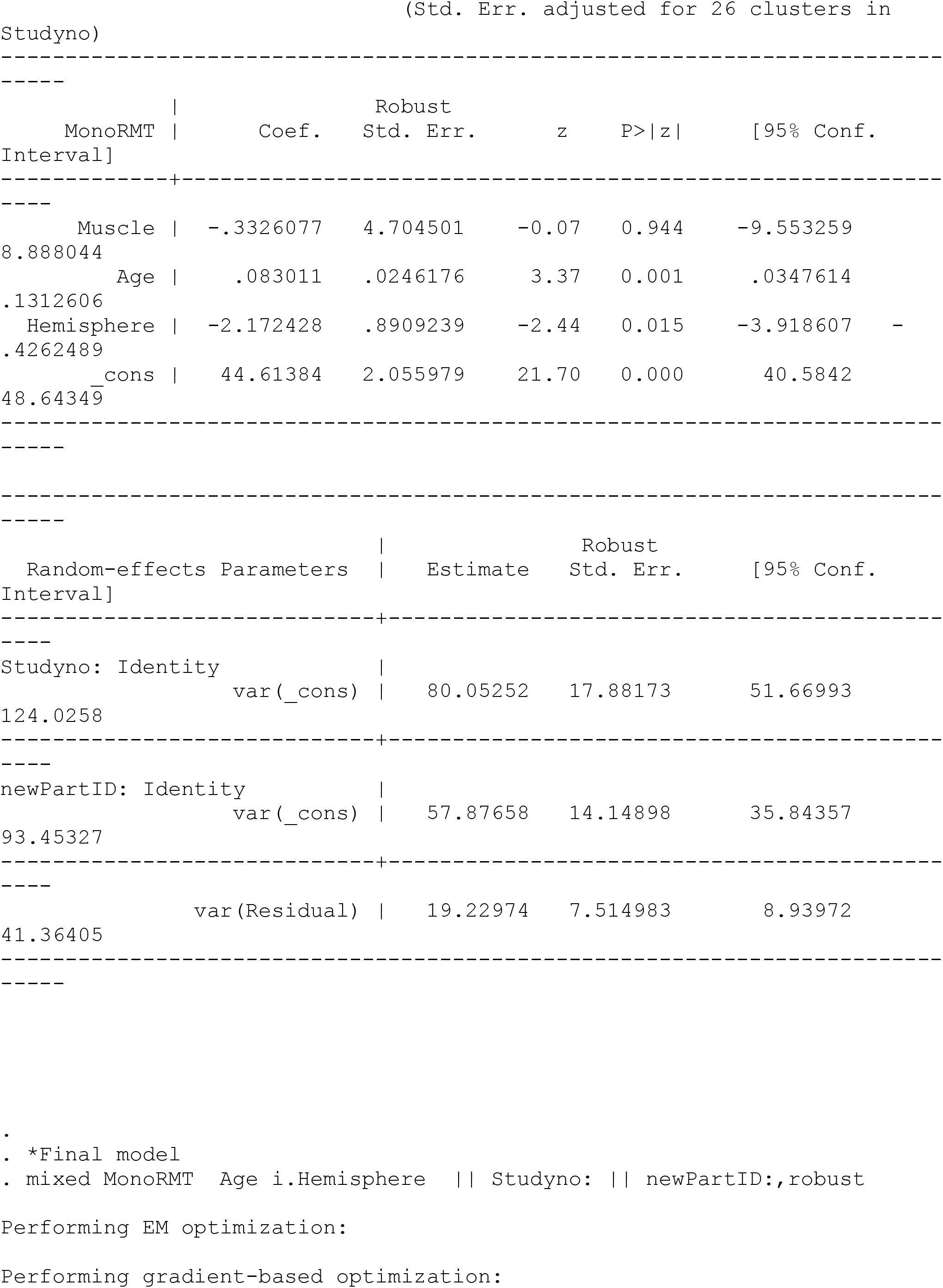
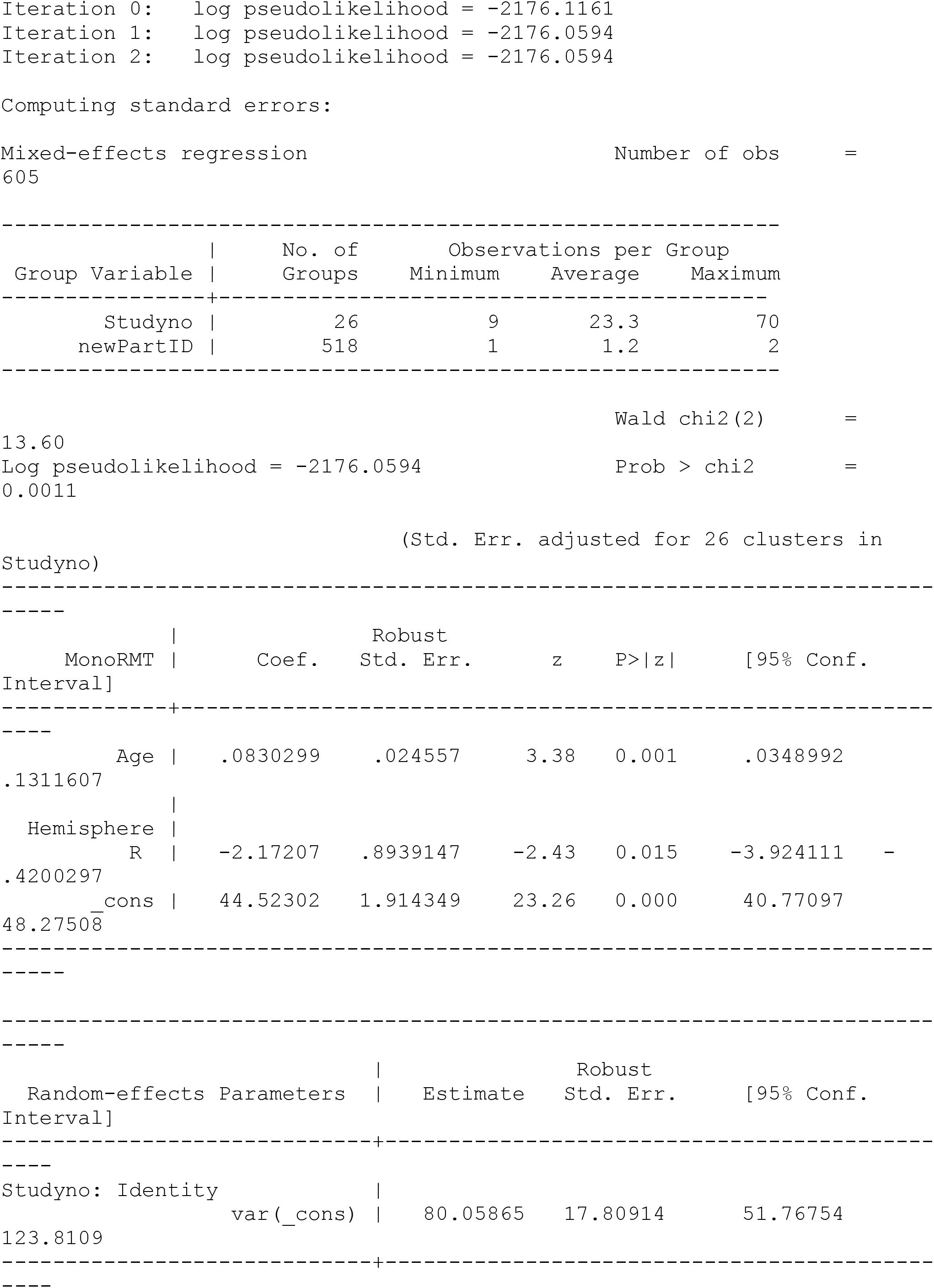
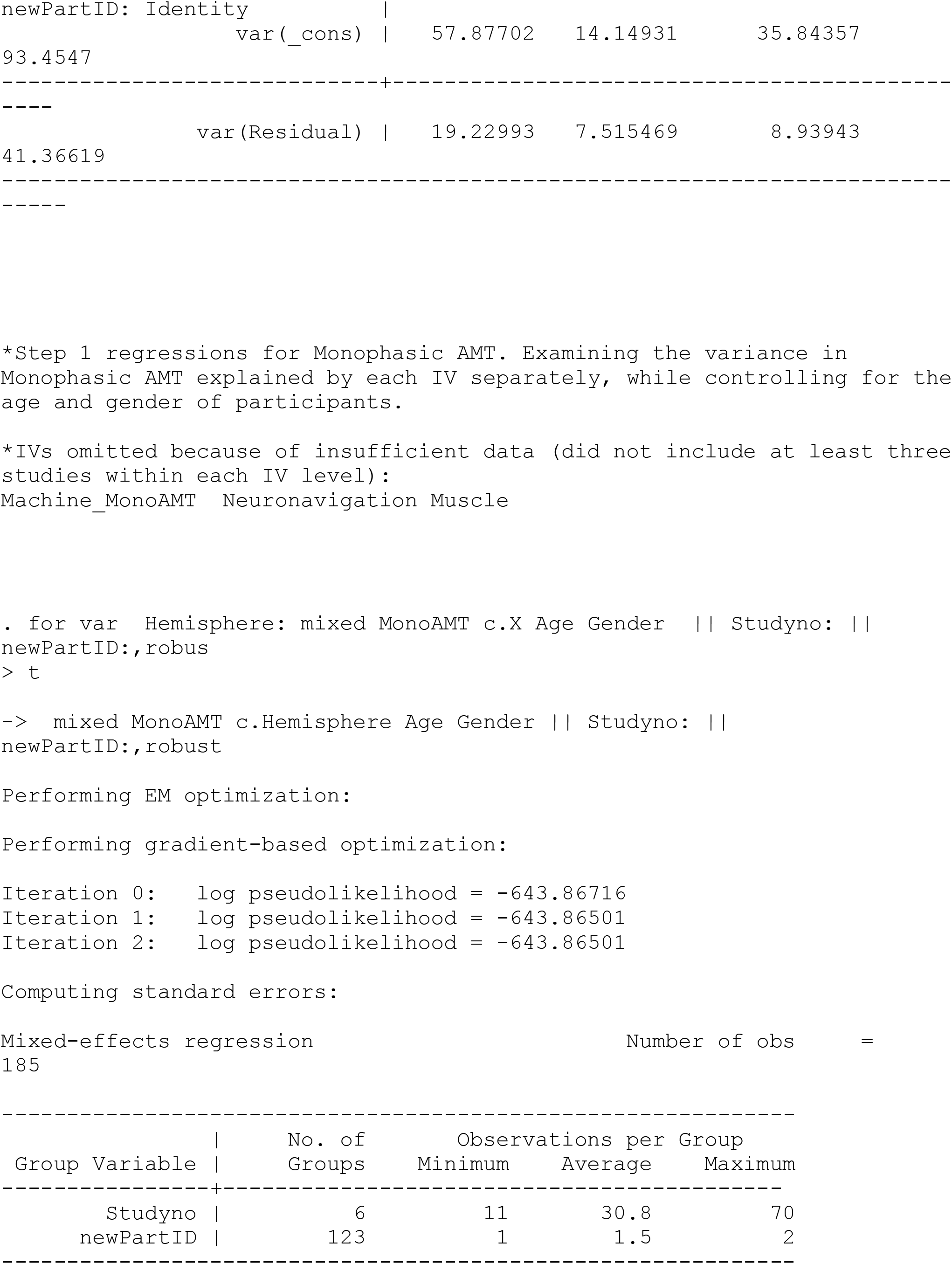
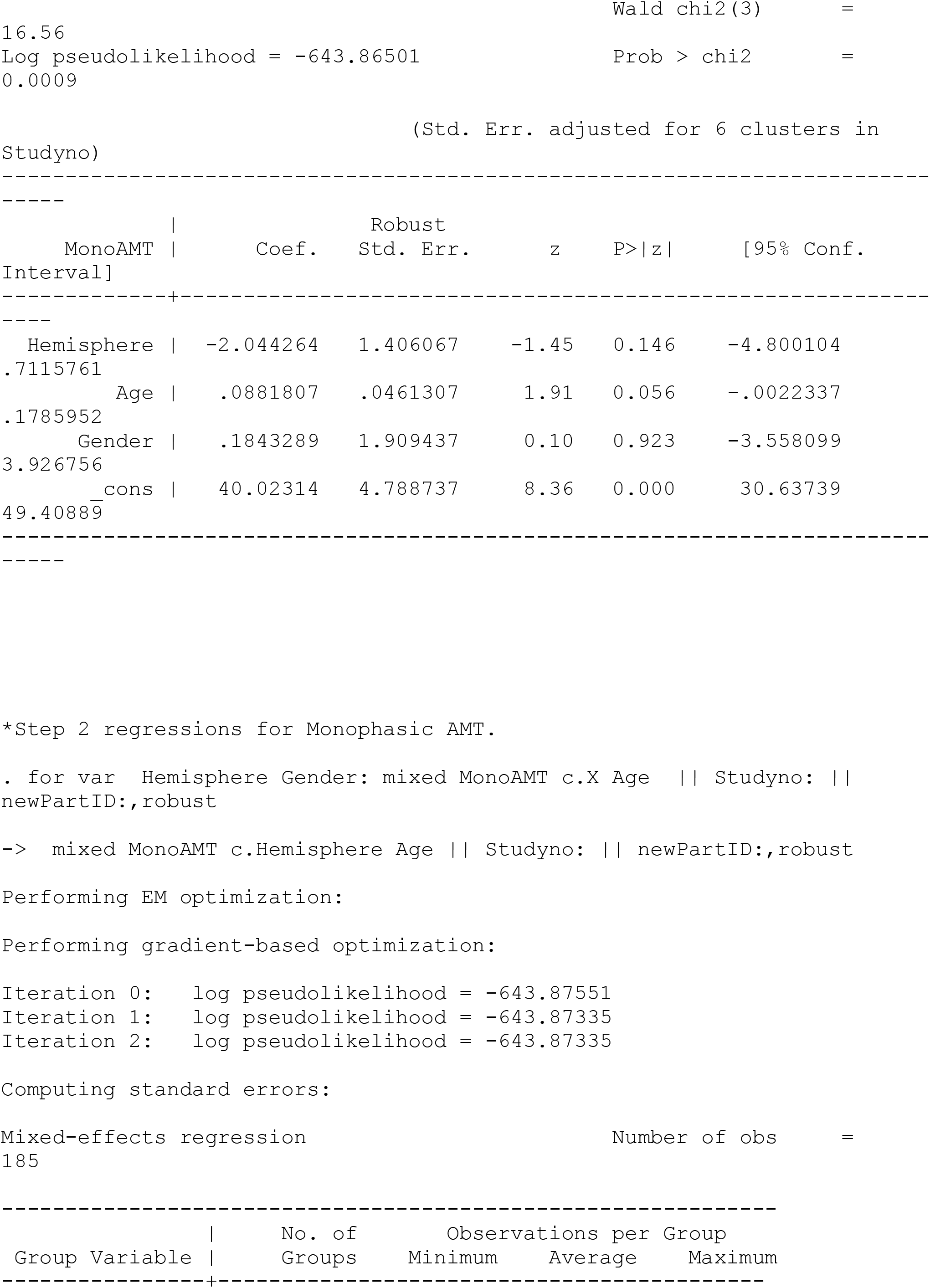
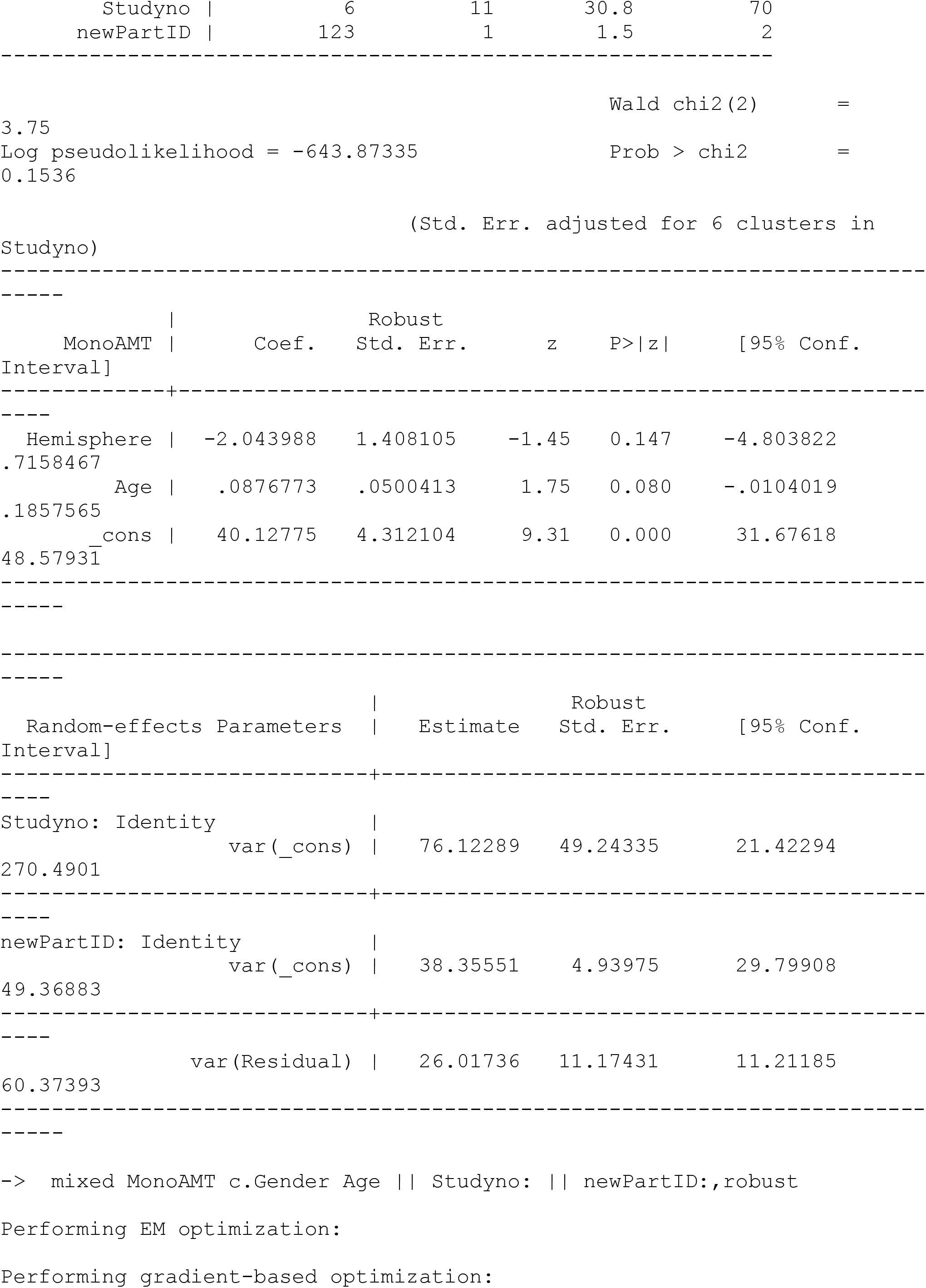
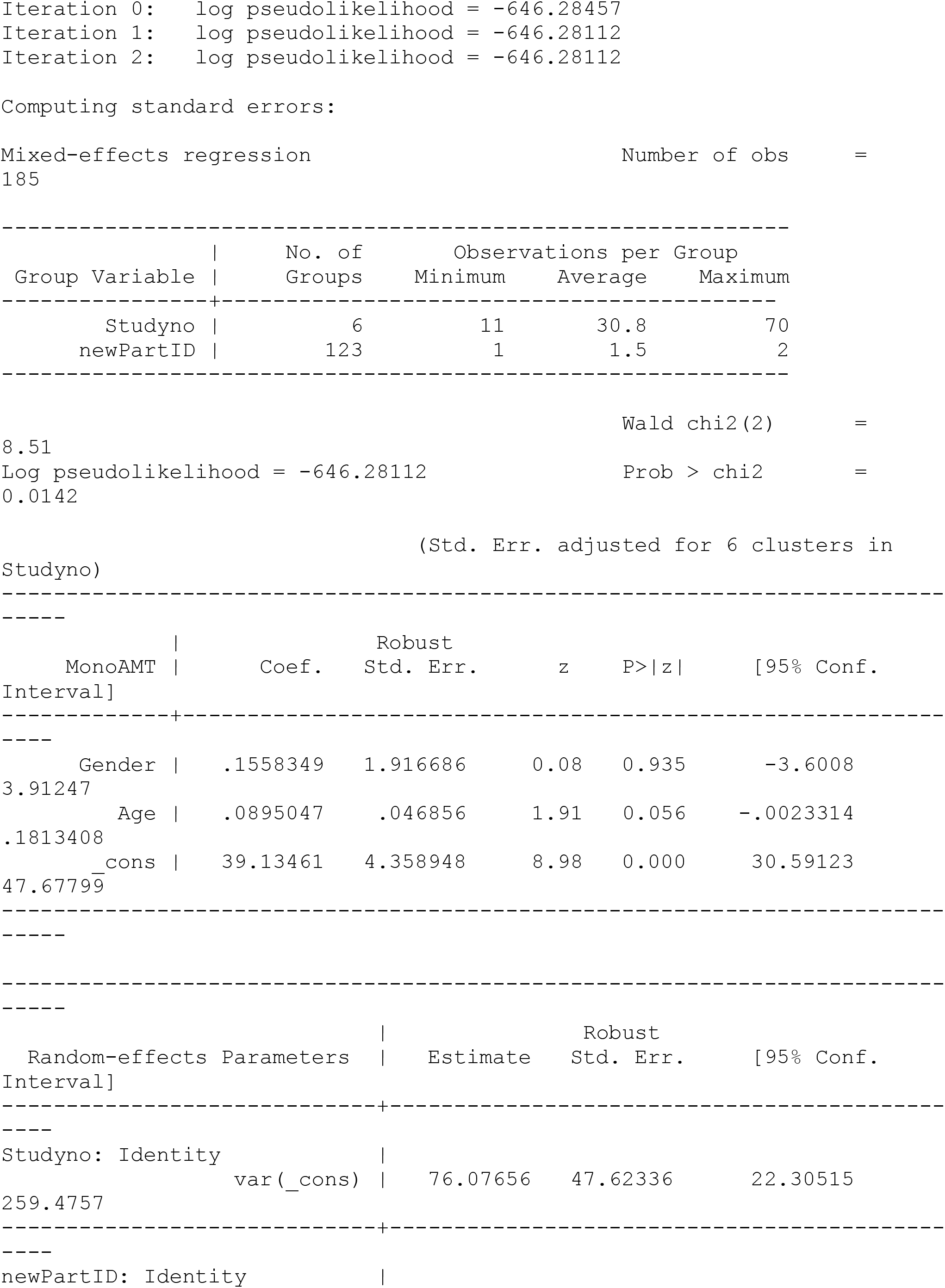
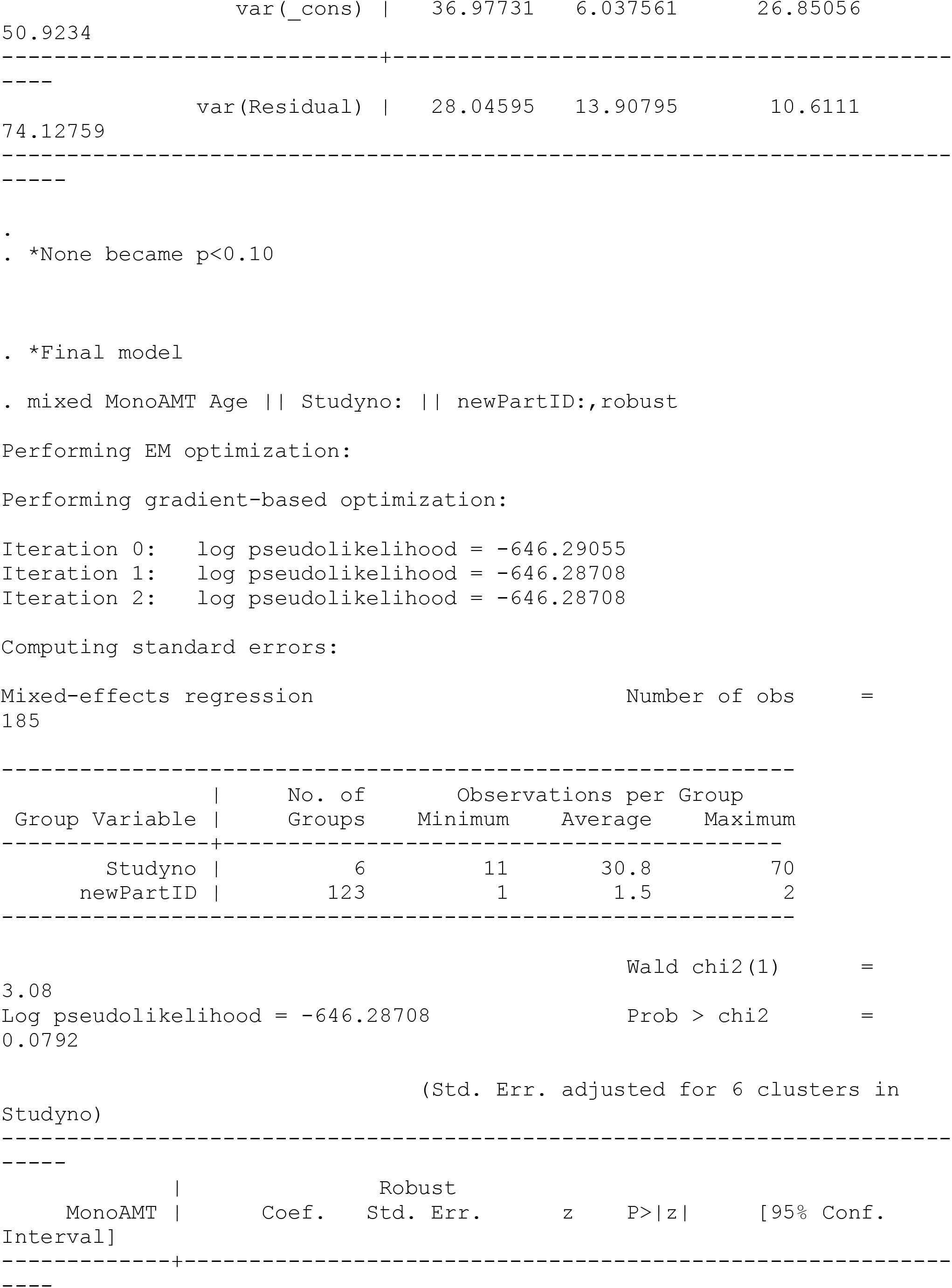
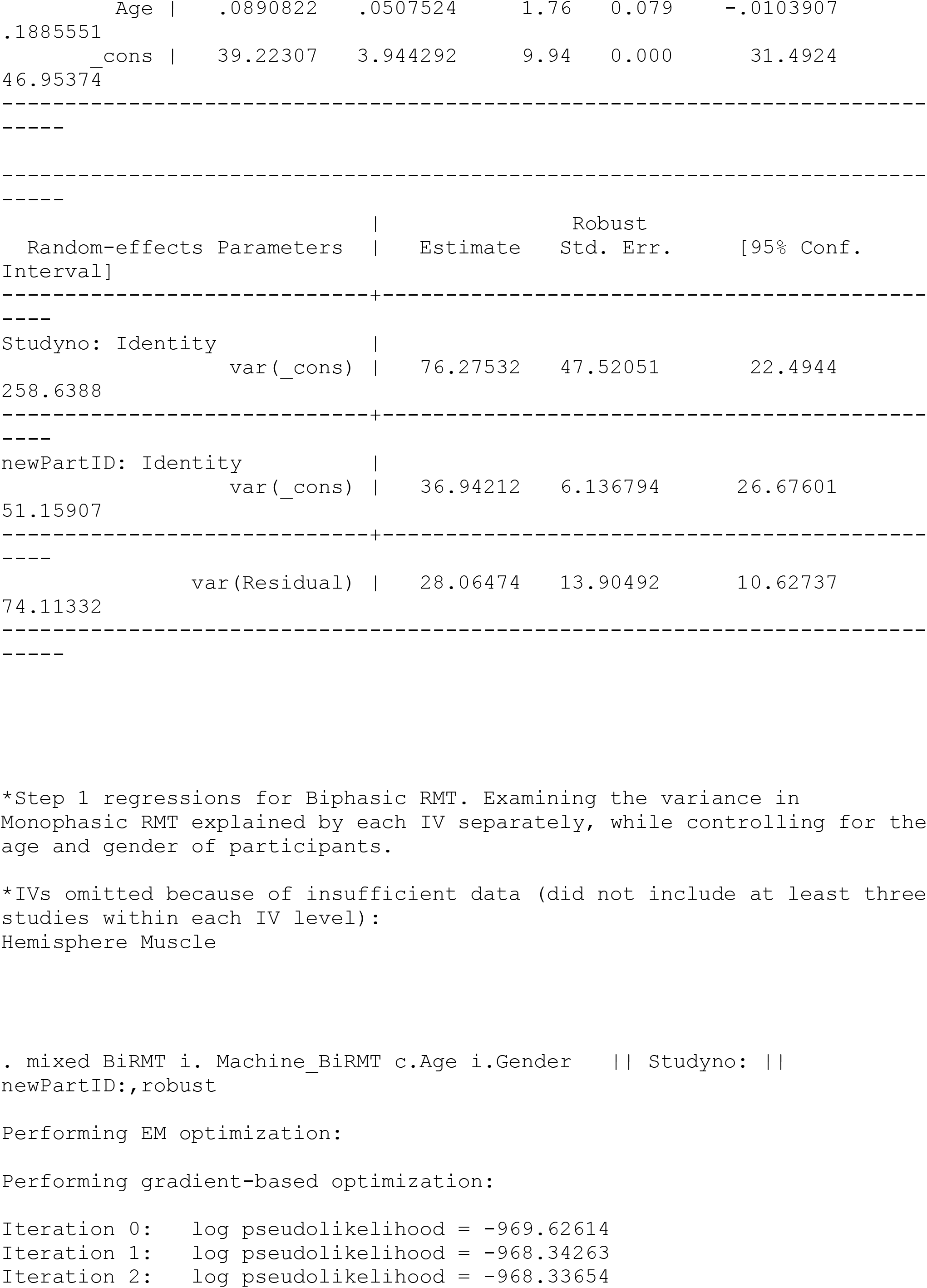
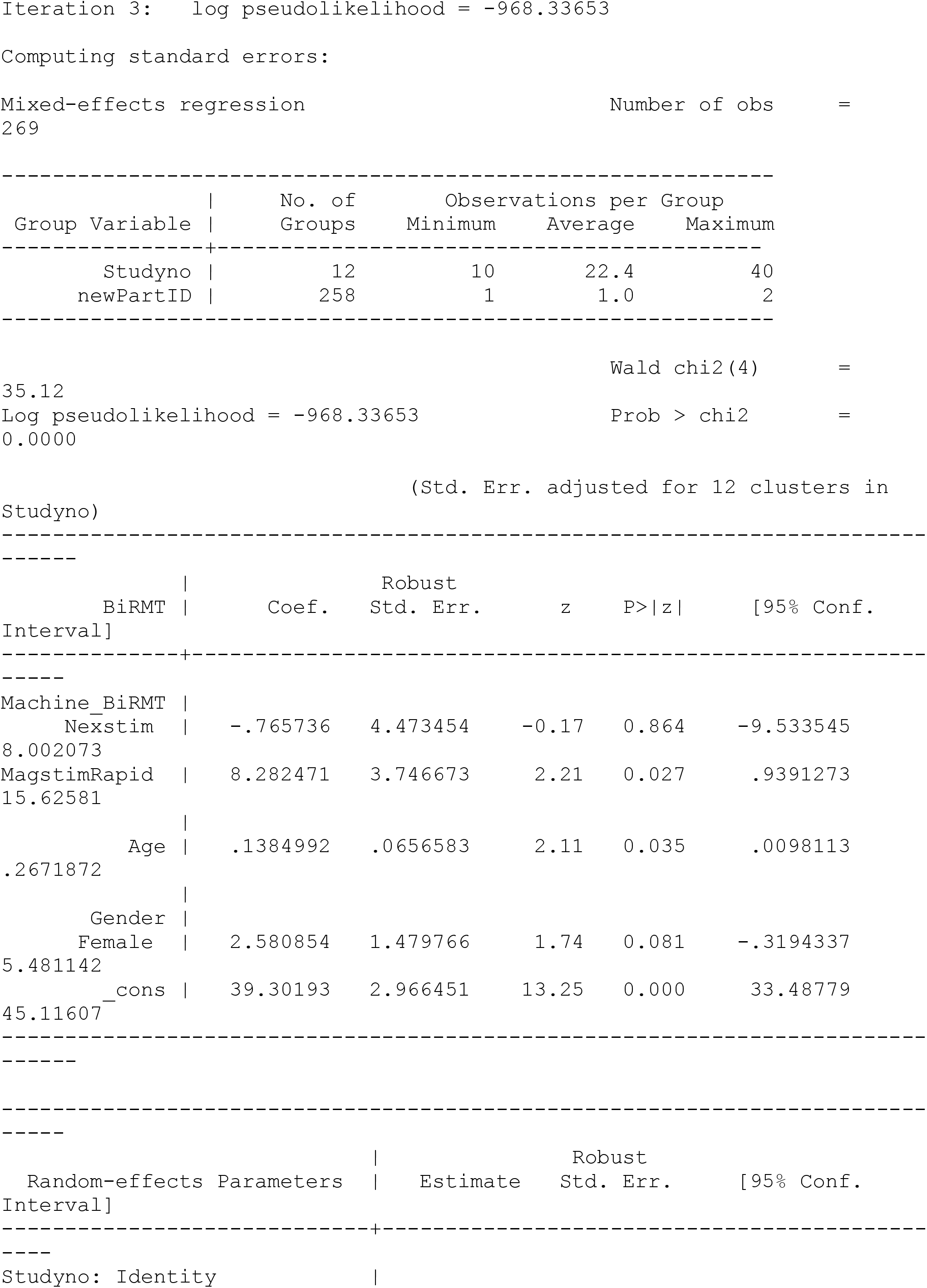
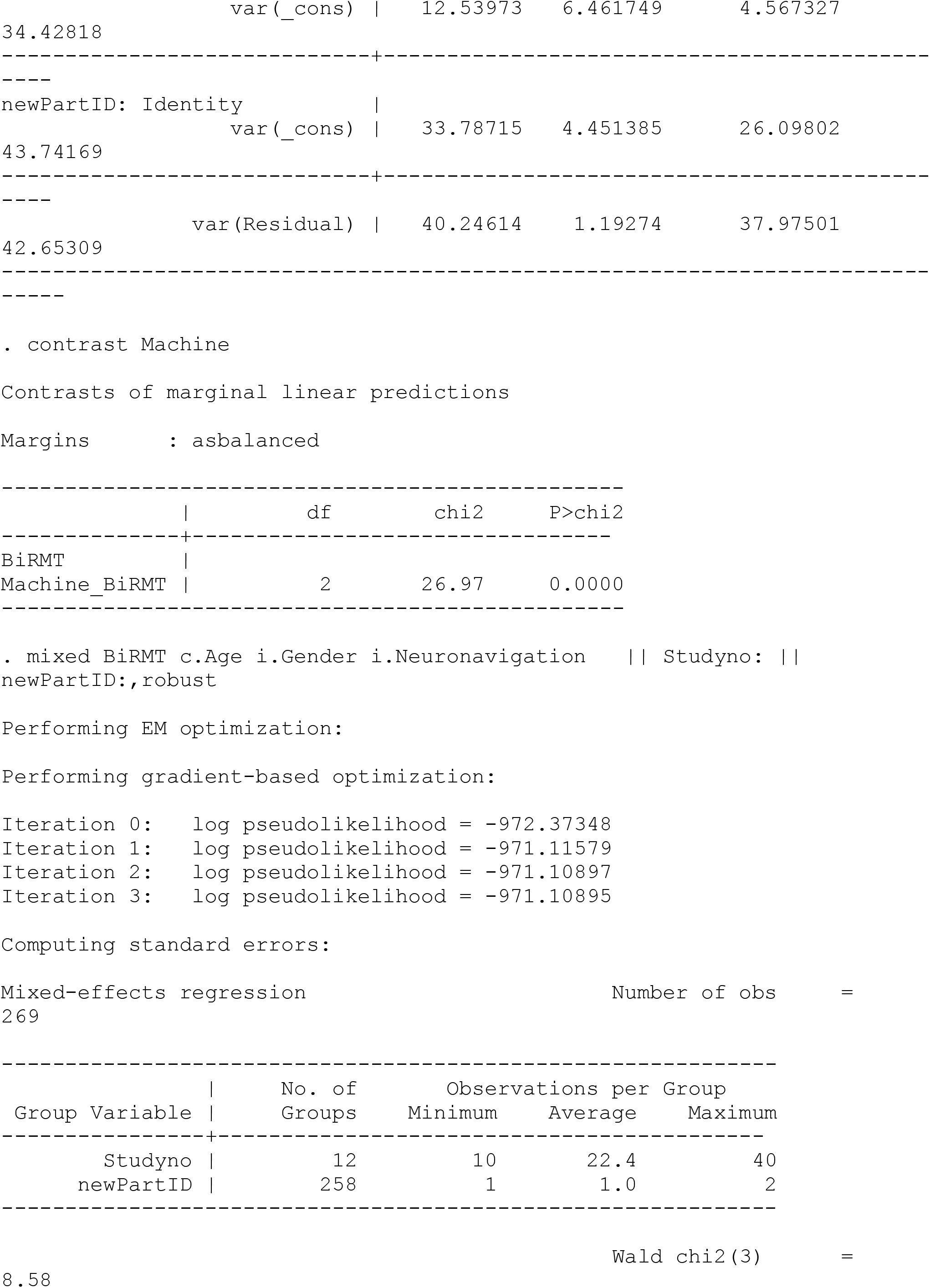
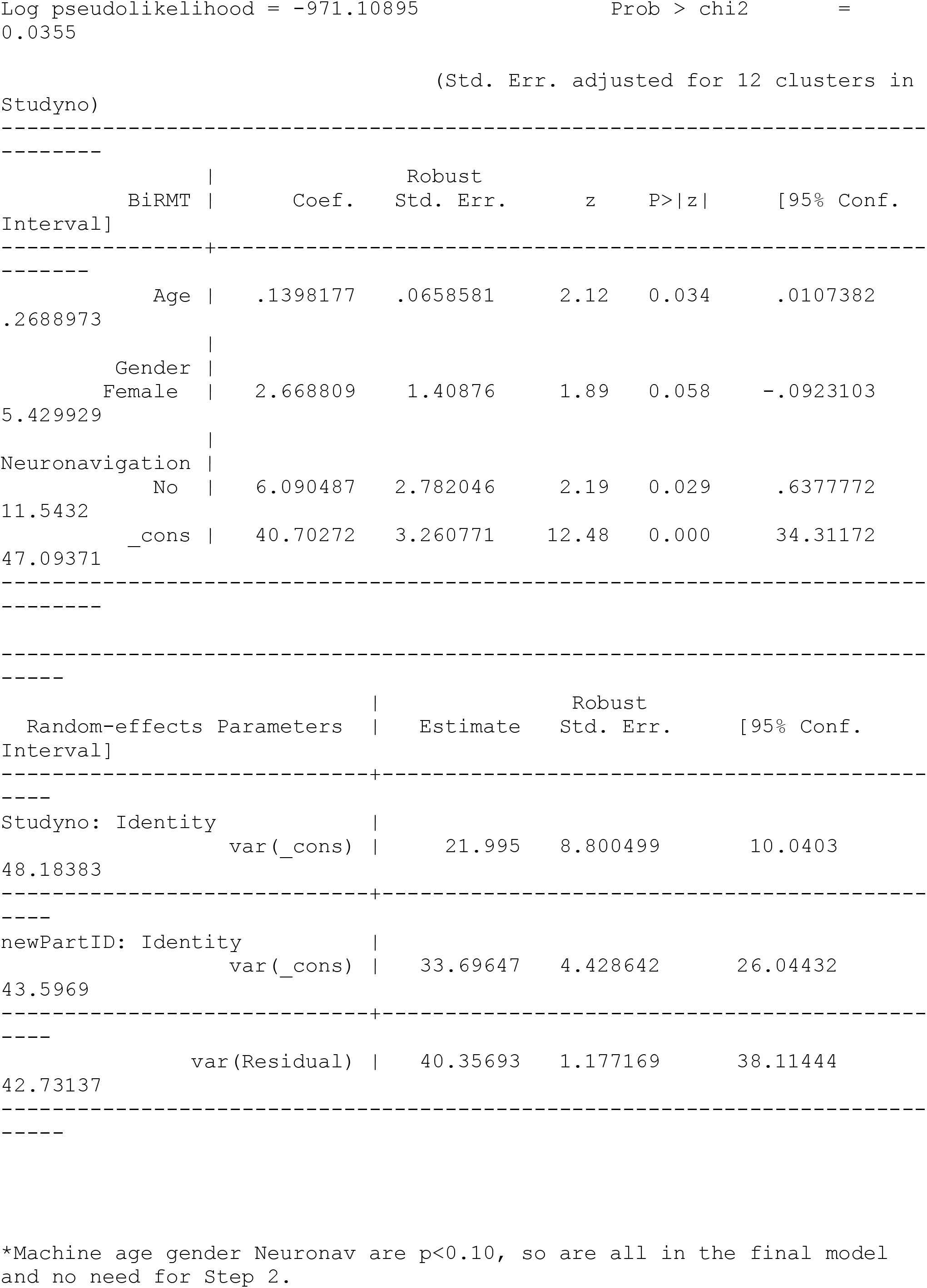
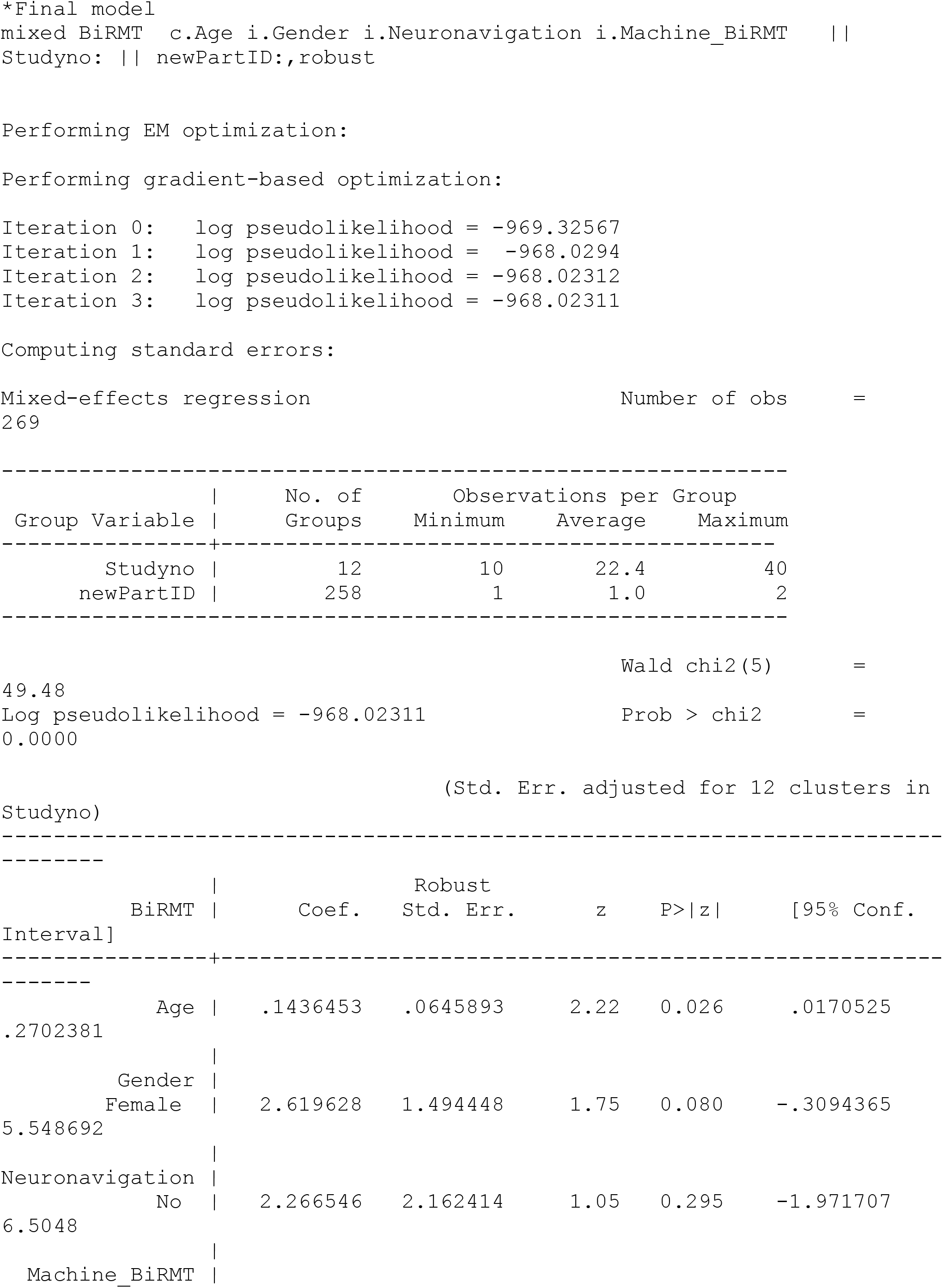
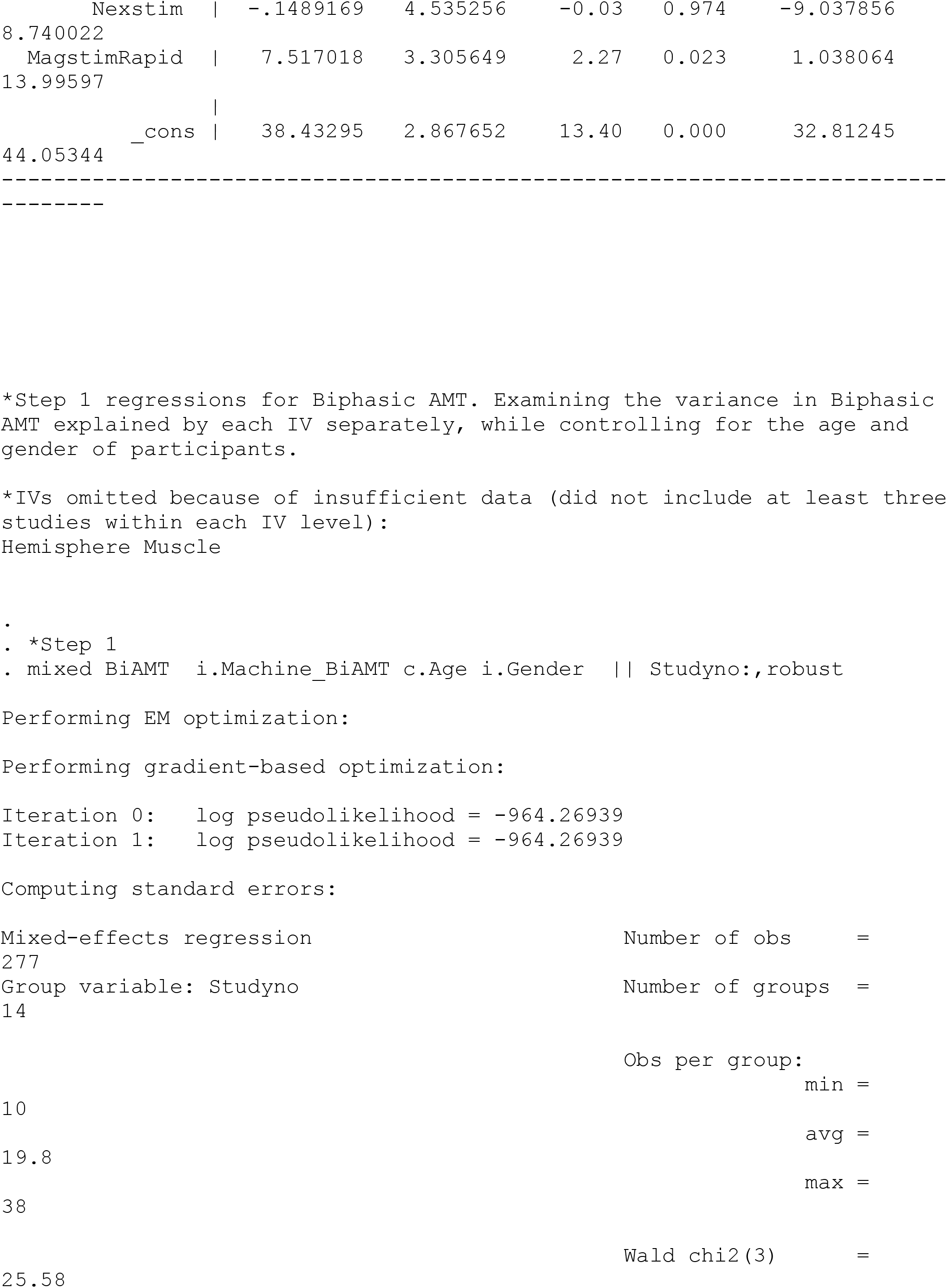
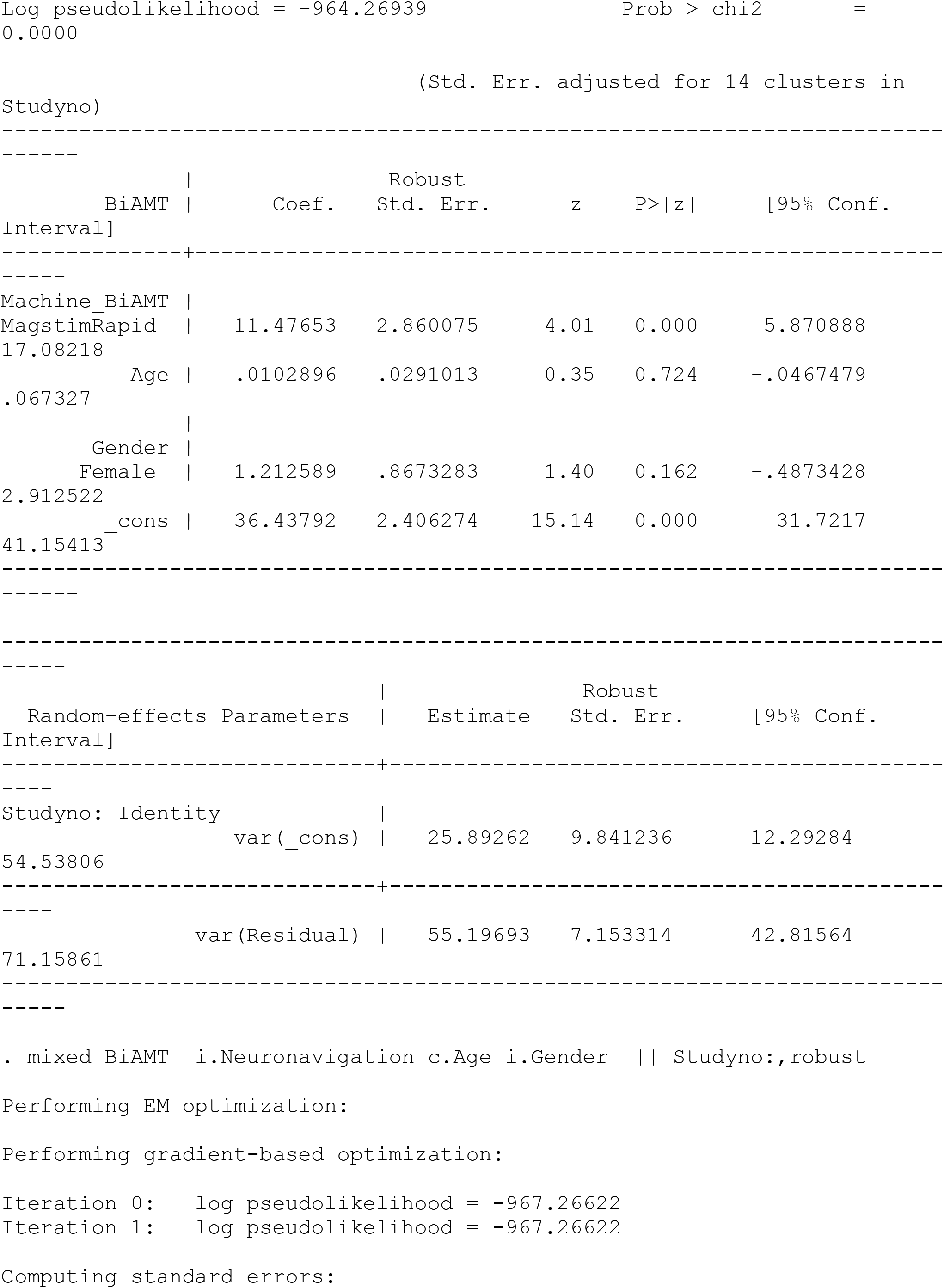
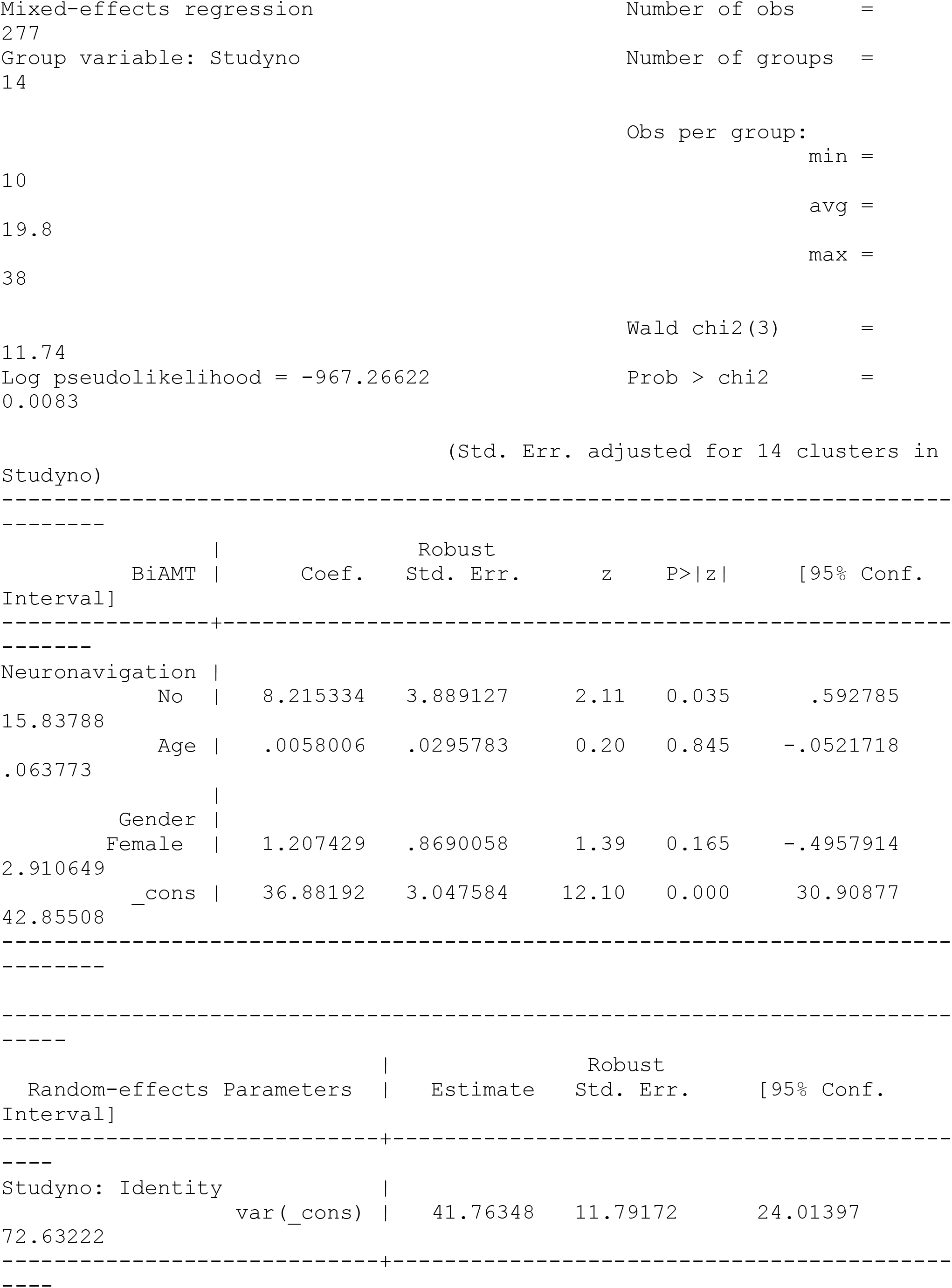
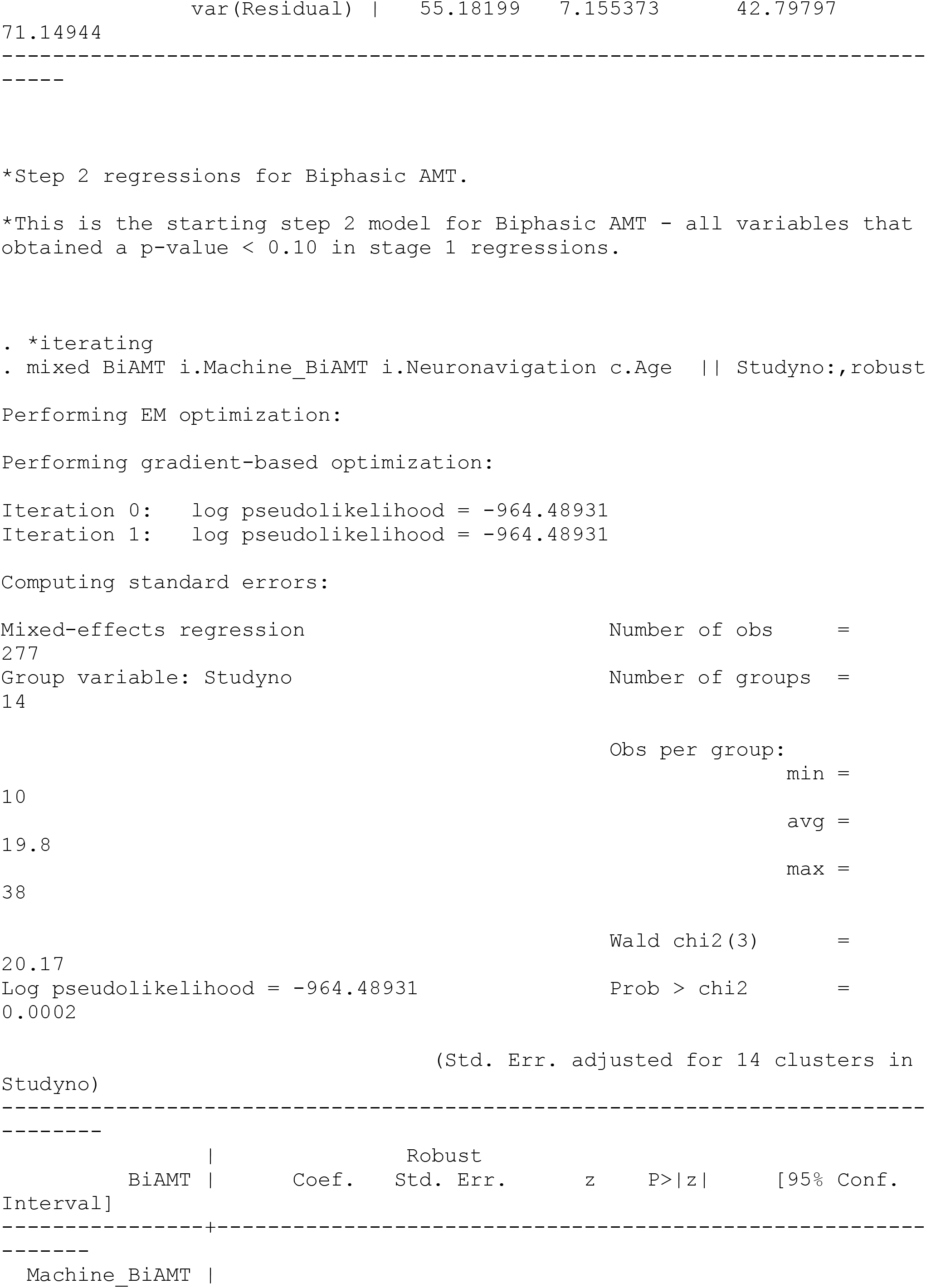
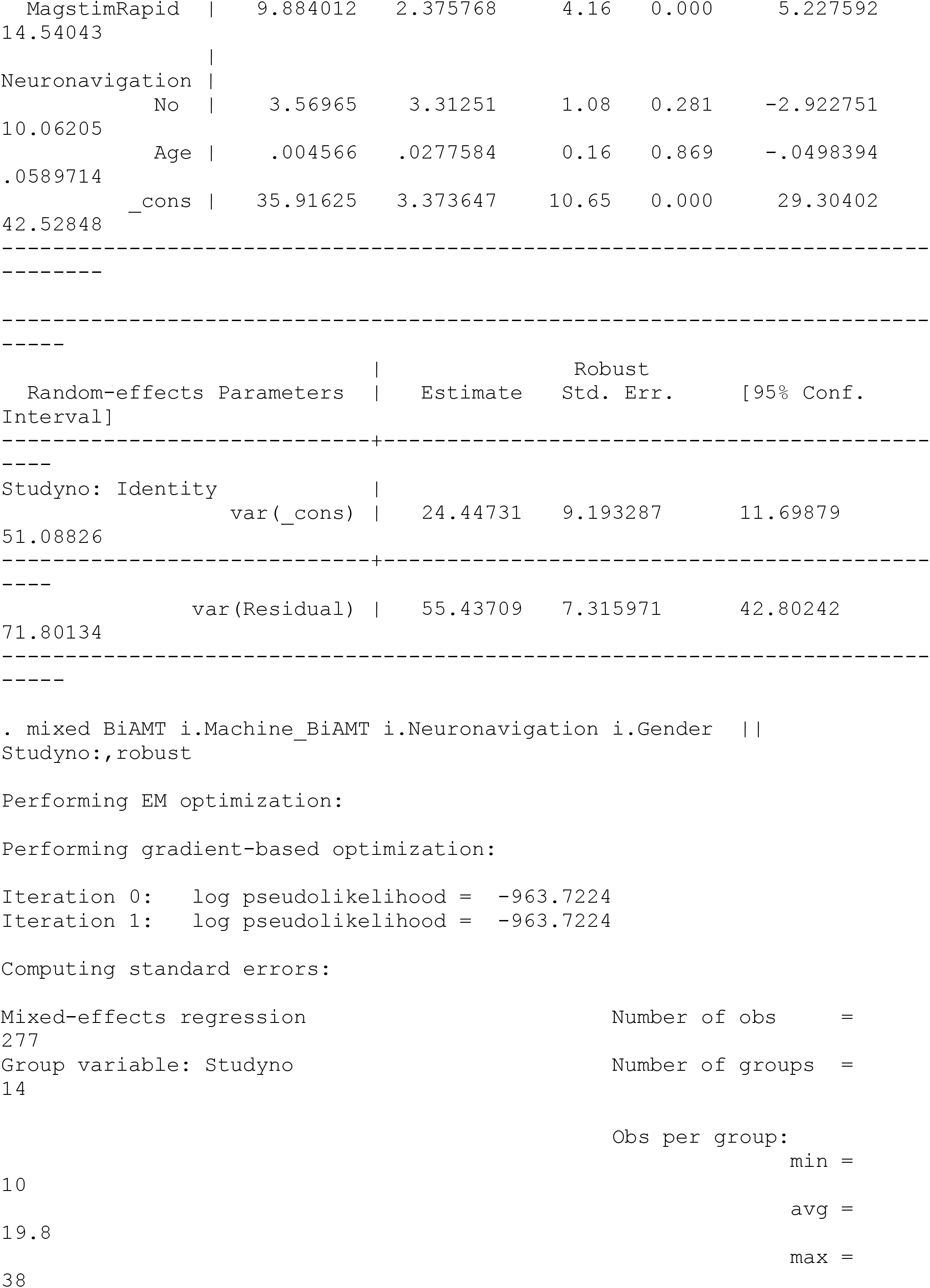
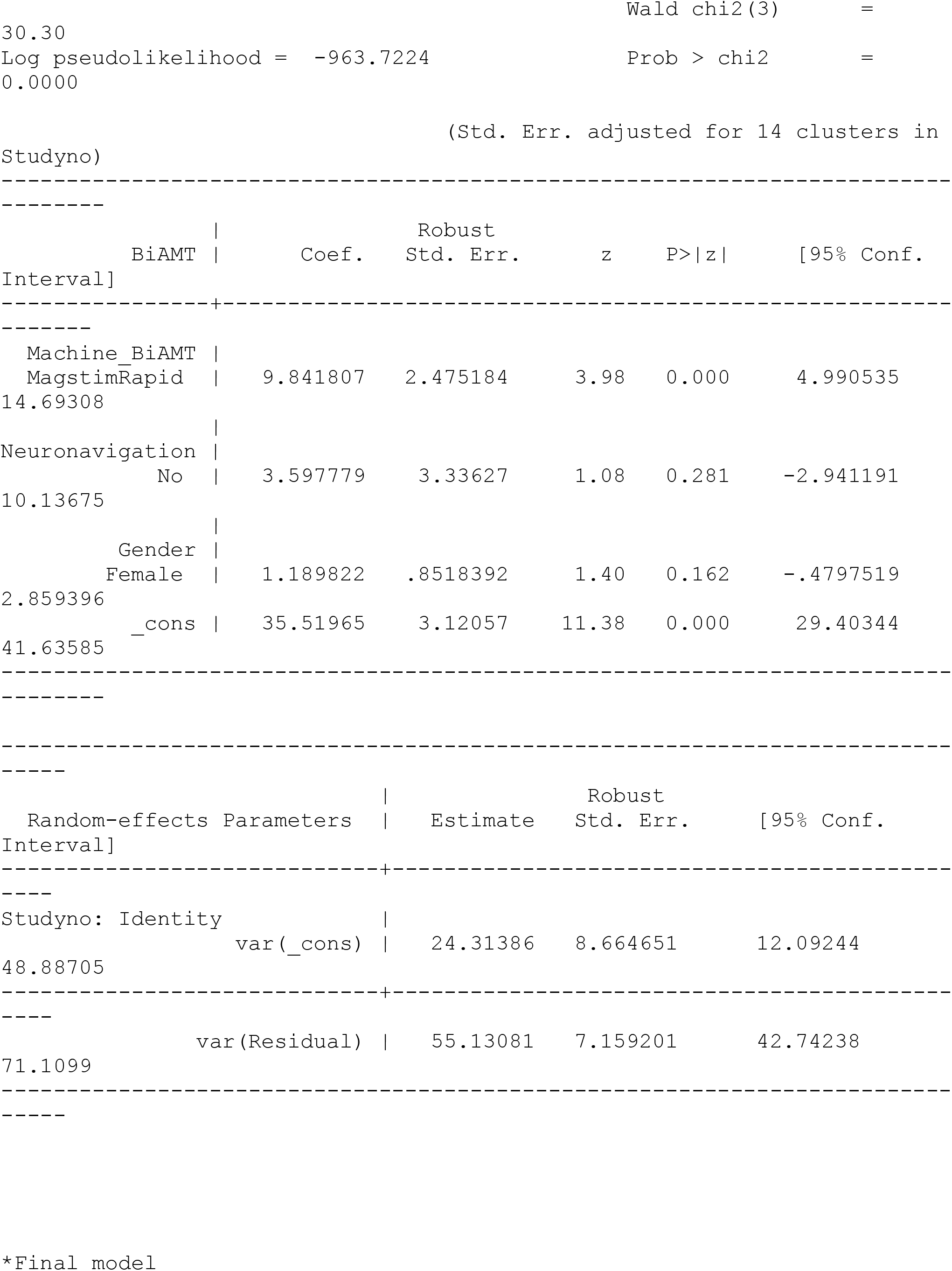
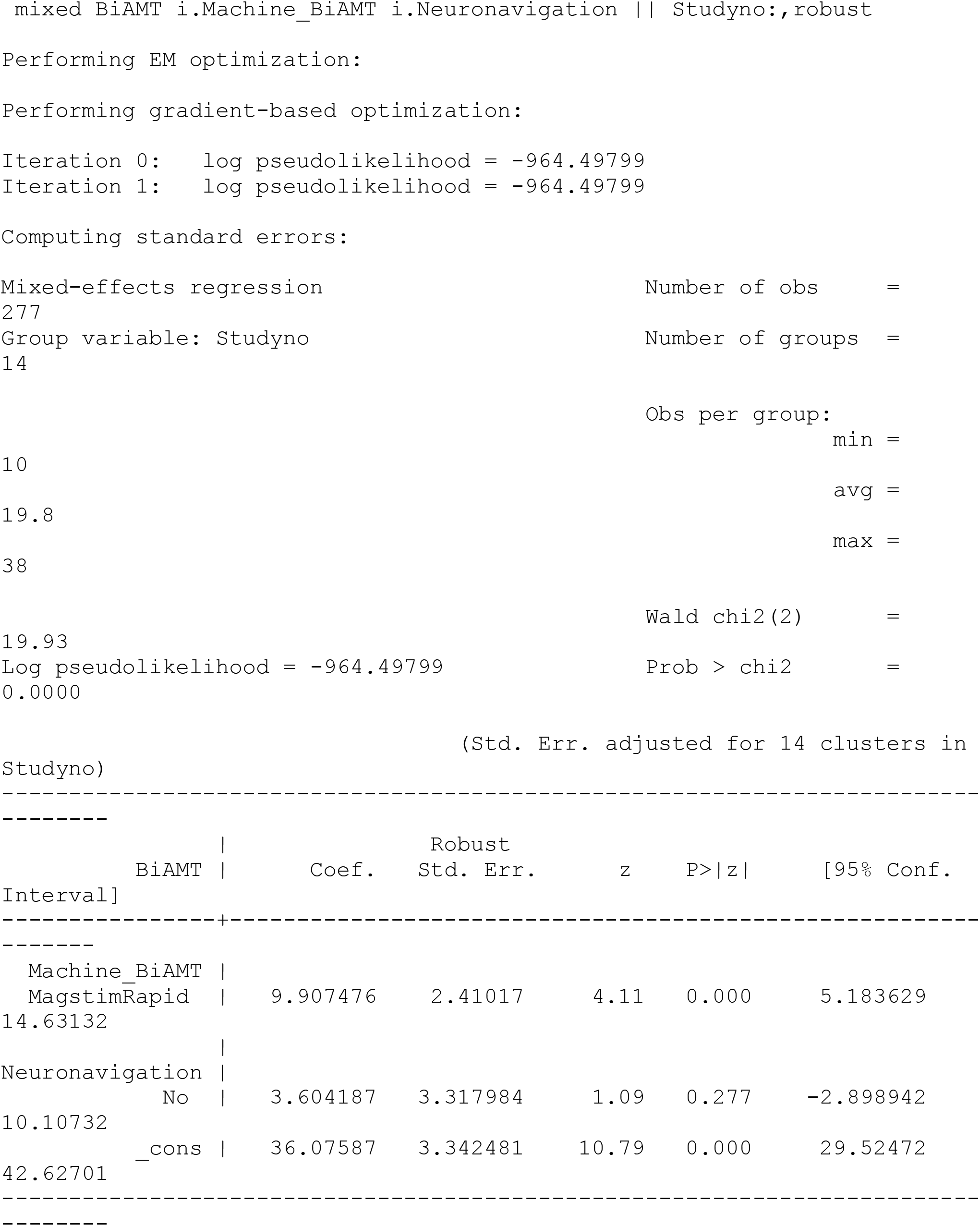

